# Ploidy variation and its implications for reproduction and population dynamics in two sympatric Hawaiian coral species

**DOI:** 10.1101/2021.11.21.469467

**Authors:** Timothy G. Stephens, Emma L. Strand, Hollie M. Putnam, Debashish Bhattacharya

## Abstract

Standing genetic variation is a major driver of fitness and resilience, and therefore of fundamental importance for threatened species such as stony corals. We analyzed RNA-seq data generated from 132 *Montipora capitata* and 119 *Pocillopora acuta* coral colonies collected from Kāneʻohe Bay, Oʻahu, Hawaiʻi. Our goals were to determine the extent of colony genetic variation and to study reproductive strategies in these two sympatric species. Surprisingly, we found that 63% of the *P. acuta* colonies were triploid, with putative independent origins of the different triploid clades. These corals have spread primarily *via* asexual reproduction and are descended from a small number of genotypes, whose diploid ancestor invaded the bay. In contrast, all *M. capitata* colonies are diploid, outbreeding, with almost all colonies genetically distinct. Only two cases of asexual reproduction, likely *via* fragmentation, were identified in this species. We report two distinct strategies in sympatric coral species that inhabit the largest sheltered body of water in the main Hawaiian Islands. These data highlight divergence in reproductive behavior and genome biology, both of which contribute to coral resilience and persistence.

**Significance Statement:** Given the threat posed to coral reef ecosystems by human caused climate change, there is a growing focus on developing strategies for the protection and restoration of these critical marine habitats. These efforts are however limited by our understanding of the diversity of coral survival and reproductive strategies. Our analysis of data from two coral species inhabiting the same Hawaiian bay found that one is a strict sexual outbreeder, whereas the other reproduces predominantly asexually (i.e., clonally) and includes both diploids and triploids. These results broaden our understanding of coral biology, adaptability, and evolution, and underpin future research into the mechanisms of coral resilience that can inform restoration activities.

## Introduction

Given ongoing climate change, it is critical to understand how rapidly changing ocean conditions impact coral population biology and resilience and how the innate adaptability of coral populations may contribute to their persistence (Cant, et al. 2021; Fischer, et al. 2021). For coral reef ecosystems, which depend on the nutritional symbiosis between scleractinian coral hosts and their single celled dinoflagellate (algal) endosymbionts in the family Symbiodiniaceae (LaJeunesse, et al. 2018), thermal stress may lead to dysbiosis and mortality. This phenomenon is known as coral “bleaching”, whereby symbiotic cells and pigments are expelled or lost from the host tissue, leaving the bright white color of the underlying coral animal body and skeleton (van Oppen and Lough 2009). Bleaching is the primary cause of mass coral mortality (Hughes, et al. 2017).

Coral reefs are also threatened by ocean acidification resulting from the increased amount of CO2 in the atmosphere that dissolves in the surface ocean, changing the carbonate chemistry and lowering the pH (Hoegh-Guldberg, et al. 2007). Understanding the mechanisms that underlie the coral response to long-term environmental stress is, however, challenging, given the genetically diverse collection of organisms (cnidarian animal host, algal symbionts, prokaryotic microbiome, fungi and other eukaryotes, and viruses) that comprise the holobiont and contribute to its health and resilience (Veron 2000). Furthermore, corals are impacted by persistent abiotic stresses (e.g., diurnal, and seasonal light and temperature variation) and a plethora of interacting taxa (e.g., algae, fish, viruses) that are of non-holobiont provenance, making these complex models for field studies.

We previously generated high-quality genome assemblies from two Hawaiian coral species (Stephens, et al. 2022). The first is at chromosome-level from the rice coral, *Montipora capitata*, and comprises 14 large scaffolds that likely represent the 14 chromosomes predicted in other *Montipora* species (Kenyon 1997). The second is from the cauliflower coral, *Pocillopora acuta*, and is the first polyploid (i.e., triploid) genome assembly generated from Scleractinia. Whereas the mechanisms that give rise to polyploidy in corals, and its effects on organismal fitness, are unknown, it can result from genome duplication within a species (autopolyploidy), or from hybridization of two different species (allopolyploidy). This process often precipitates drastic changes in cell organization and genome structure and can alter gene expression, genome stability, cell physiology, and the cell cycle (Wertheim, et al. 2013). In some animals, triploidy may be beneficial with respect to improved growth and pathogen resistance (Kang and Rosenwaks 2008). This observation peaks interest in corals with respect to how changes in their genomic configuration may contribute to the evolution of stress resistant genotypes. To advance understanding of ploidy variation in corals, and differences in reproductive strategies of sympatric species, we generated and analyzed RNA-seq data from fragments (i.e., nubbins) of *M. capitata* and *P. acuta* colonies collected from across six reefs in Kāneʻohe Bay, a 45 km² sheltered water body in Oʻahu, Hawaiʻi. Analysis of single-nucleotide polymorphisms (SNPs) in each coral sample was used to investigate genetic diversity, ploidy, and reproductive strategy in these two sympatric species.

## Results

### Ploidy differences in Kāneʻohe Bay corals

Transcriptome data were collected from 119 *P. acuta* (fig. 1A) and 132 *M. capitata* (fig. 1B) coral nubbins (Supplementary tables S1 and S2), each derived from a different coral colony. Colonies were sampled from six reefs in Kāneʻohe Bay, Oʻahu, Hawaiʻi (fig. 1C). Analysis of the *P. acuta* RNA-seq data using the program nQuire predicted (using data pre- and post-denoising) that 44 (37% of the 119 total) samples are derived from diploid genets (i.e., at genomic loci with two alleles, each present in ∼50% of the reads, producing an allele frequency distribution with a single peak at roughly 0.5; Supplementary fig. S1A, B; Supplementary table S3; Supplementary data S1). In contrast, 75 (63%) samples are from triploid genets (i.e., at genomic loci with two alleles, one present in ∼33% and the other in ∼66% of the reads, producing an allele frequency distribution with two peaks at roughly 0.33 and 0.66). The Pacuta_HTHC_TP11_2185 sample (Supplementary fig. S1E), which was predicted to be a triploid, was generated from the same coral colony as the *P. acuta* reference genome (Stephens, et al. 2022).

**Fig. 1.**
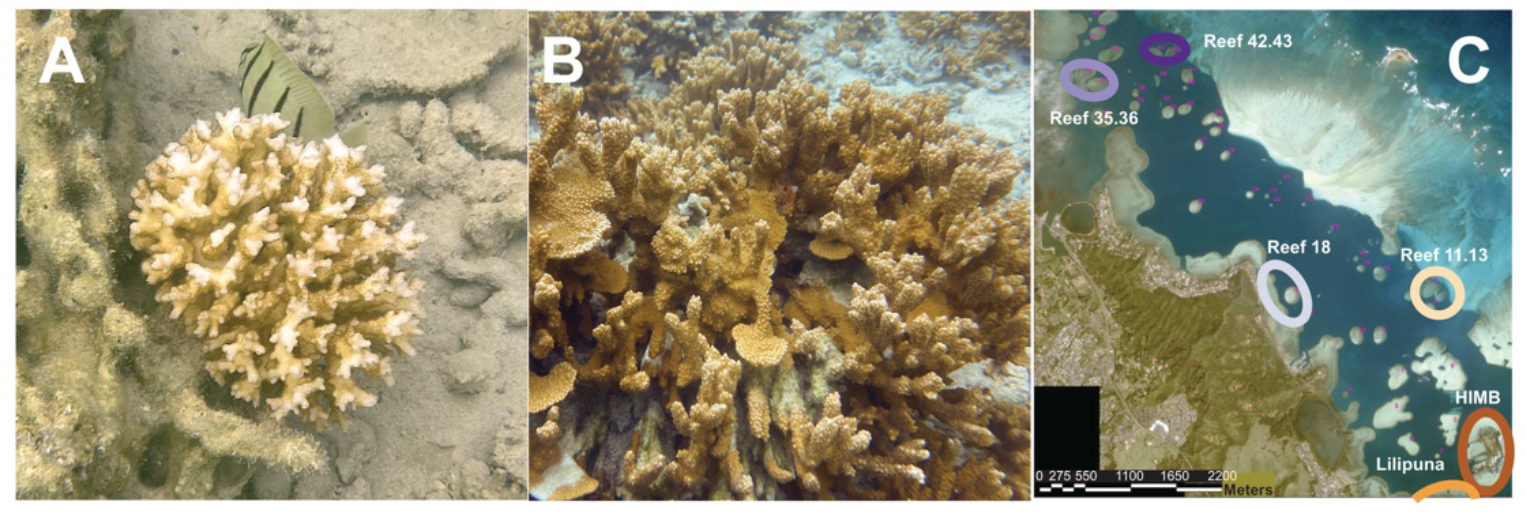
Study corals in Kāne‘ohe Bay, Oʻahu, Hawaiʻi. Colonies of (**A**) *Montipora capitata*, the rice coral, and (**B**) *Pocillopora acuta*, the lace coral. Images taken by D. Bhattacharya. (**C**) Aerial image of Kāne‘ohe Bay (modified from https://dlnr.hawaii.gov/) with the six reefs where the coral samples were collected highlighted using colored circles and labels. The color of the circles surrounding the collection sites corresponds to the colors used in all subsequent main text and supplementary figures. The legend for distance in meters is shown at the bottom of the image.

This genome was shown to be triploid using *k*-mer based methods (see Stephens, et al. 2022). The presence of this sample in our analysis supports the accuracy of our approach for ploidy determination using RNA-seq data. nQuire predicted that all additional Hawaiian *P. acuta* samples (n=32) from sites outside of Kāneʻohe Bay, acquired from the NCBI Sequence Read Archive (SRA; see Materials and Methods), were diploid (Supplementary fig. S1A, B; Supplementary table S3; Supplementary data S2).

In contrast, all the *M. capitata* samples analyzed in this study are diploid (using data pre- denoising), except for Mcapitata_HTAC_TP12_1632 and Mcapitata_ATAC_TP11_1644, that were identified as a tetraploid and a triploid (respectively; Supplementary table S3; Supplementary fig. S1C samples highlighted in red). In the denoised data Mcapitata_HTAC_TP12_1632 remained a putative tetraploid (Supplementary fig. S1D) where as Mcapitata_ATAC_TP11_1644 was identified as a diploid, although the latter did have a higher delta Log-Likelihood value than most of the other samples, showing that the diploid model did not fit this sample as well as it did for the others. Whereas it is possible that Mcapitata_HTAC_TP12_1632 is a tetraploid, the SNP allele frequency distributions (Supplementary data S3; Supplementary fig. S1F) do not strongly support this hypothesis. Tetraploid samples would have three peaks in the distribution at approximately 0.25, 0.5, and 0.75 along the x-axis (Weiss, et al. 2018), which is not observed for this sample (Supplementary fig. S1F). Instead, the distribution has no visible middle peaks, but does have a much higher frequency of alleles at the tails of the distribution. The distribution also has a higher frequency of alleles with vales around 0.1 and 0.9, with an increase in the frequency of alleles occurring around 0.2 and 0.8, which is not observed in any of the other *M. capitata* samples. This contrasts with the other *M. capitata* samples in which there is a single (approximately) uniform peak at 0.5 and the tails of the distribution did not show an increase in allele frequency. The increase in frequency of alleles that occupied the tails of the distribution could result from the samples being a mix of (at least two) different genotypes. If one of the genotypes in the sample was dominant (present at a much higher frequency than the others) then we would see an increase in the frequency of SNP alleles with support towards the ends of the distribution that do not conform to any of the expected peak structures. These SNPs, rather than originating from heterozygous regions of the different haplotypes within a single genotype, represent heterozygous regions between different genotypes, and thus the proportion of reads that support the alleles reflects the relative abundances of the different genotypes and not the ploidy of the sample. It is also possible that such a scenario can explain the variability in the prediction of the ploidy of Mcapitata_ATAC_TP11_1644, although obviously to a lesser degree. *M. capitata* samples (n=27, see methods for criteria) downloaded from SRA to compare with this study were predicted by nQuire to be diploids (Supplementary table S3; Supplementary data S4). One of the *M. capitata* samples (SRR5453755) was identified as a triploid in the non-denoised data, although this is likely caused by the sample being from a colony that is comprised of multiple genets.

### Population structure of *P. acuta* in Kāne‘ohe Bay

Several approaches were used to make pair-wise comparisons of the 119 *P. acuta* RNA- seq samples to assess relatedness and determine if any were derived from clones (i.e., colonies derived from the same genet), given indications of clonal relationships in past studies (Combosch and Vollmer 2013; Gorospe and Karl 2013; Yeoh and Dai 2010).

Sample relationship was initially assessed using the proportion of shared single nucleotide polymorphisms (SNPs; fig. 2A; see Materials and Methods), with a threshold of > 94% used to aggregate samples into groups: i.e., assumed to represent clonal samples. This threshold was chosen based on the distribution of shared SNPs (fig. 2B) between each pairwise combination of samples; the set of pairwise comparisons captured by this threshold (regions shaded bright yellow in Figure 2) is clearly separated from the other distinct sets of comparisons observed in the distribution. This threshold is very close to the 95% similarity threshold applied in another coral study (Locatelli and Drew 2019). In total, there are 8 clonal groups (Groups 1-8) which comprise 113/119 (94.96%) of the *P. acuta* samples (fig. 2). Groups 1-4 are triploids, whereas Groups 5-8 are diploid. Only two triploid and four diploid samples were ungrouped. Generally, the ungrouped samples had higher similarity with samples of the same ploidy, however, the diploid sample Pacuta_HTHC_TP5_1415 had higher similarity to the triploid samples compared with the diploid samples (fig. 2). An additional 32 *P. acuta* samples (collected from locations not in Kāne‘ohe Bay) were downloaded from SRA and incorporated into the SNP analysis. These samples were all derived from a single experiment (BioProject: PRJNA435468; Poquita-Du, et al. 2019) and represent three genotypes. Each genotype was collected from a separate reef near Singapore, and had been fragmented into multiple ramets, each of which underwent RNA sequencing as part of a stress experiment (Supplementary table S4). The proportion of shared SNPs between the samples derived from each of the three genotypes was ∼98%, which is very similar to the values observed between many of the putative clonal samples generated in this study (e.g., ∼97% between the samples in Groups 4, 5, and 6; Supplementary fig. S2; Supplementary table S5).

**Fig. 2.**
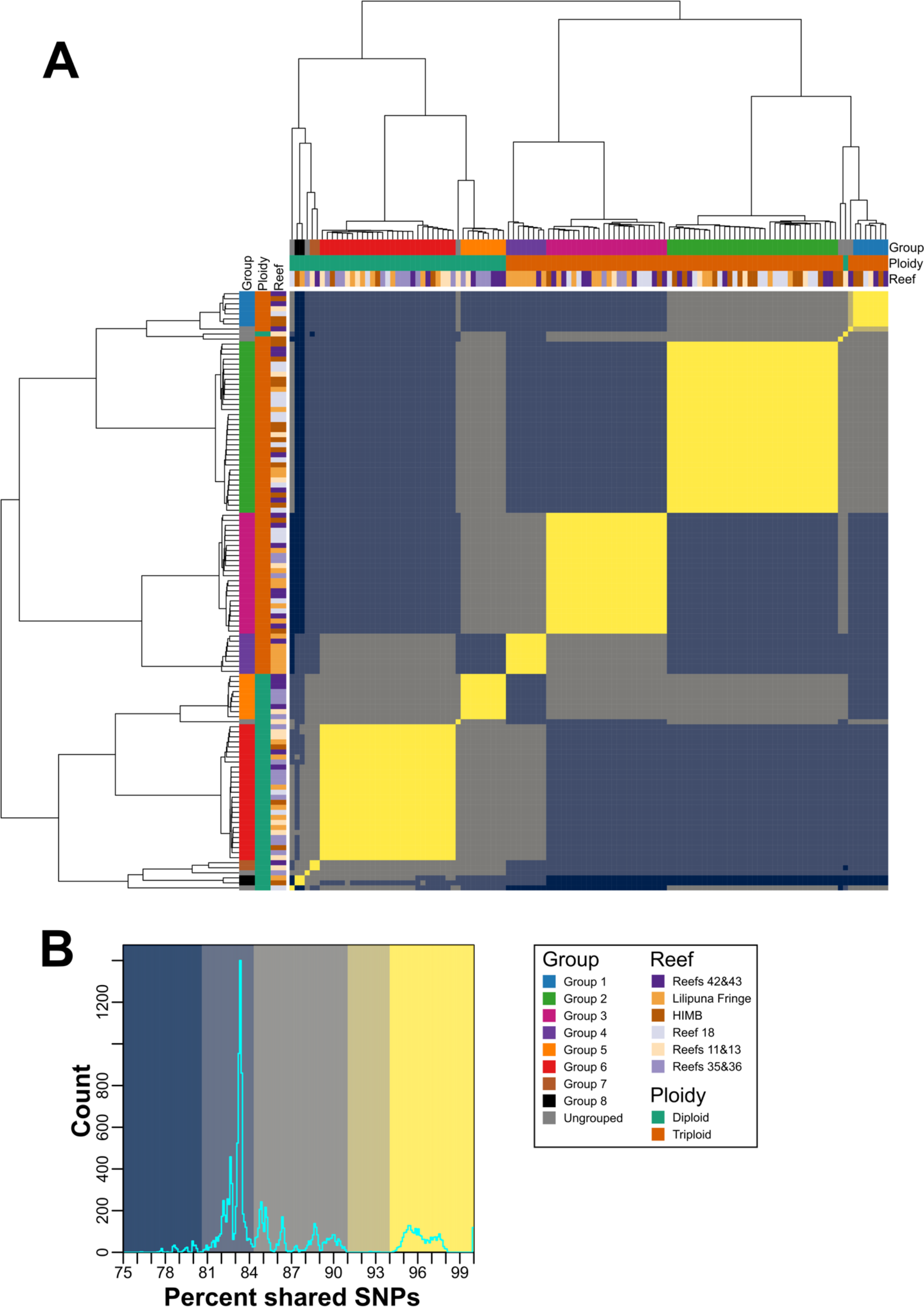
Proportion of shared SNPs between each combination of *P. acuta* samples. (**A**) Heatmap displaying the proportion of shared SNPs between all pair-wise combinations of *P. acuta* RNA-seq samples sequenced in this study. The putative clonal groups identified in this study (>94% shared SNPs) are shown as colored bars along the top and left sides of the heatmap; a legend describing the color of each group is presented on the bottom right side of the image. The order of the column and rows, and the dendrograms presented on the top and left sides of the heatmap, were generated by hierarchical clustering of the proportion of shared SNPs. (**B**) Histogram of the proportion of shared SNPs in the heatmap; the background colors used in the histogram correspond to the colors used in the heatmap presented in (**A**). The colors used in the heatmap and histogram were manually chosen to highlight the distinct sets of proportions of shared SNPs observed in the histogram presented in (**B**).

The relatedness metric produced by vcftools (originally proposed by Manichaikul, et al. 2010; see Materials and Methods) agrees with the relationship between samples established in Figure 2, with clear grouping within, and separation of samples between, the identified groups (Supplementary fig. S3). In addition, the samples in each of the groups all have relatedness values > 0.43; values of 0.5 denote samples that are monozygotic twins (i.e., perfect clones), suggesting that the samples in each of these groups are clones, albeit not identical, because each colony has accumulated some segregating variants (Vasquez Kuntz, et al. 2022). Furthermore, the majority of *P. acuta* samples have a relatedness of around 0.25 (equivalent to the relatedness of parent and offspring, or full siblings), with almost all samples having a relatedness > 0.06 (3rd degree relatives; Supplementary fig. S3). It is also notable that within each of the groups (in particular, the larger Groups 2, 3, and 6), there appears to be subgroups of samples that have slightly higher similarity with each other (Supplementary fig. S3; Supplementary table S6). This suggests an uneven rate of spread of the clonal lineages throughout Kāne‘ohe Bay, although it is unclear if the spread of each clonal group is linked across time. *P. acuta* samples from each of the genotypes included from SRA had relatedness values of ∼0.47, which is very similar to what we observe between each of the putative clonal groups. This result supports our hypothesis that these groups represent samples from colonies that have spread through the bay *via* asexual reproduction. Furthermore, the relatedness between the three SRA genotypes, and between the SRA genotypes and the samples generated in this study, are at, or close to 0 (i.e., are unrelated individuals).

The program PCAngsd identified four ancestral populations of *P. acuta* in Kāne‘ohe Bay. The admixture results (fig. 4) are consistent with the groups shown in Figure 2, with uniformity of ancestral population profiles within each group, and separation of profiles between different groups. Interestingly, the ancestry of Pacuta_HTHC_TP5_1415 (the diploid sample that clustered with triploids, based on the SNP similarity scores) is derived from the same ancestral populations as triploid Group 1, albeit with an increased abundance of the 2nd ancestral population. This analysis demonstrates that the ancestry of each clonal group is distinct and that they have all likely arisen from separate asexual propagation events that occurred in different ancestral lineages. Principal component analysis (PCA) performed by PCAngsd using the Kāne‘ohe Bay *P. acuta* samples (fig. 5) supports the pairwise similarity and admixture results (figs. 2 and 3). That is, the samples in each clonal group form clusters along both PC1 and PC2 and the clonal groups are separated from each other. Whereas the triploid clonal groups (Groups 1-4) are clearly separated from diploid clonal groups (Groups 5-8) along PC1 and PC2, the difference between the largest triploid groups (Groups 2, 3) is roughly the same as that between these groups and the largest diploid group (Group 6). Reanalysis using a single sample per clonal group with the highest read mapping rate to the reference genome is consistent with the results produced using all samples (Supplementary fig. S4). The representative samples have congruent ancestral population profiles (albeit with only ancestral population inferred and not four; possibly due to the significantly reduced number of samples used in the analysis) and relative positions in the inferred PCA plots. These results reinforce our hypothesis that the majority of Kāne‘ohe Bay *P. acuta* samples are derived from colonies that have arisen *via* asexual reproduction, and that these events are likely to have occurred in separate related (i.e., previously mixing) lineages over an extended (currently unknown) period of time.

**Fig. 3.**
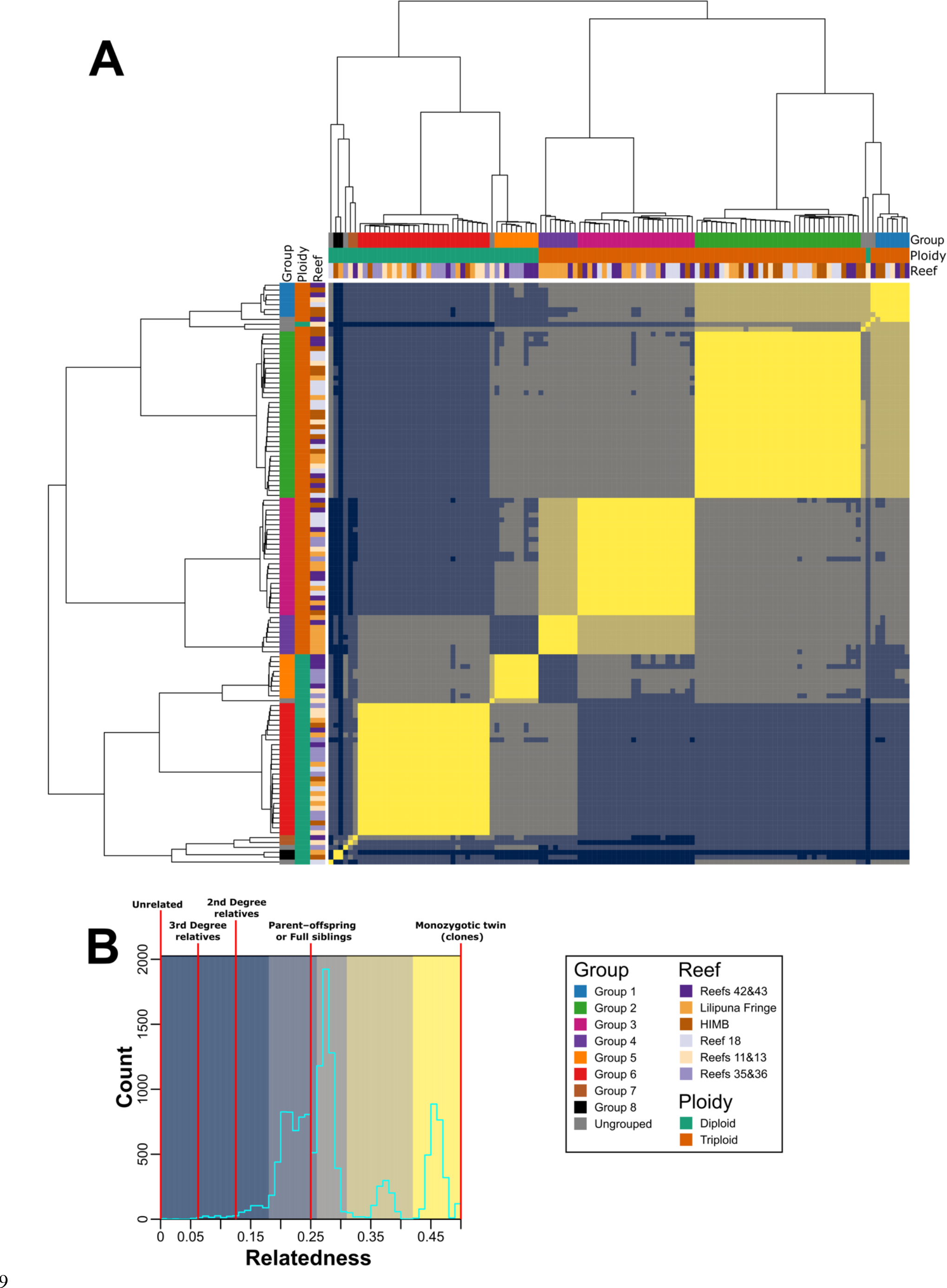
Pair-wise relatedness between each combination of *P. acuta* samples. (**A**) Heatmap displaying the relatedness scores (proposed by Manichaikul, et al. 2010) of all pair-wise combinations of *P. acuta* RNA-seq samples sequenced in this study. The putative clonal groups identified in this study are shown as colored bars along the top and left sides of the heatmap; a legend describing the color of each group is presented on the bottom right side of the image. The order of the column and rows, and the dendrograms presented on the top and left sides of the heatmap, are based on the dendrogram used in Figure 2. (**B**) Histogram of relatedness scores in the heatmap; the background colors used in the histogram correspond to the colors used in the heatmap presented in (**A**). The relatedness values that correspond to a particular relationship between samples (described in Manichaikul, et al. 2010) are annotated on the histogram using vertical red lines. The colors used in the heatmap and histogram were manually chosen to highlight the distinct sets of relatedness scores observed in the histogram presented in (**B**).

**Fig. 4.**
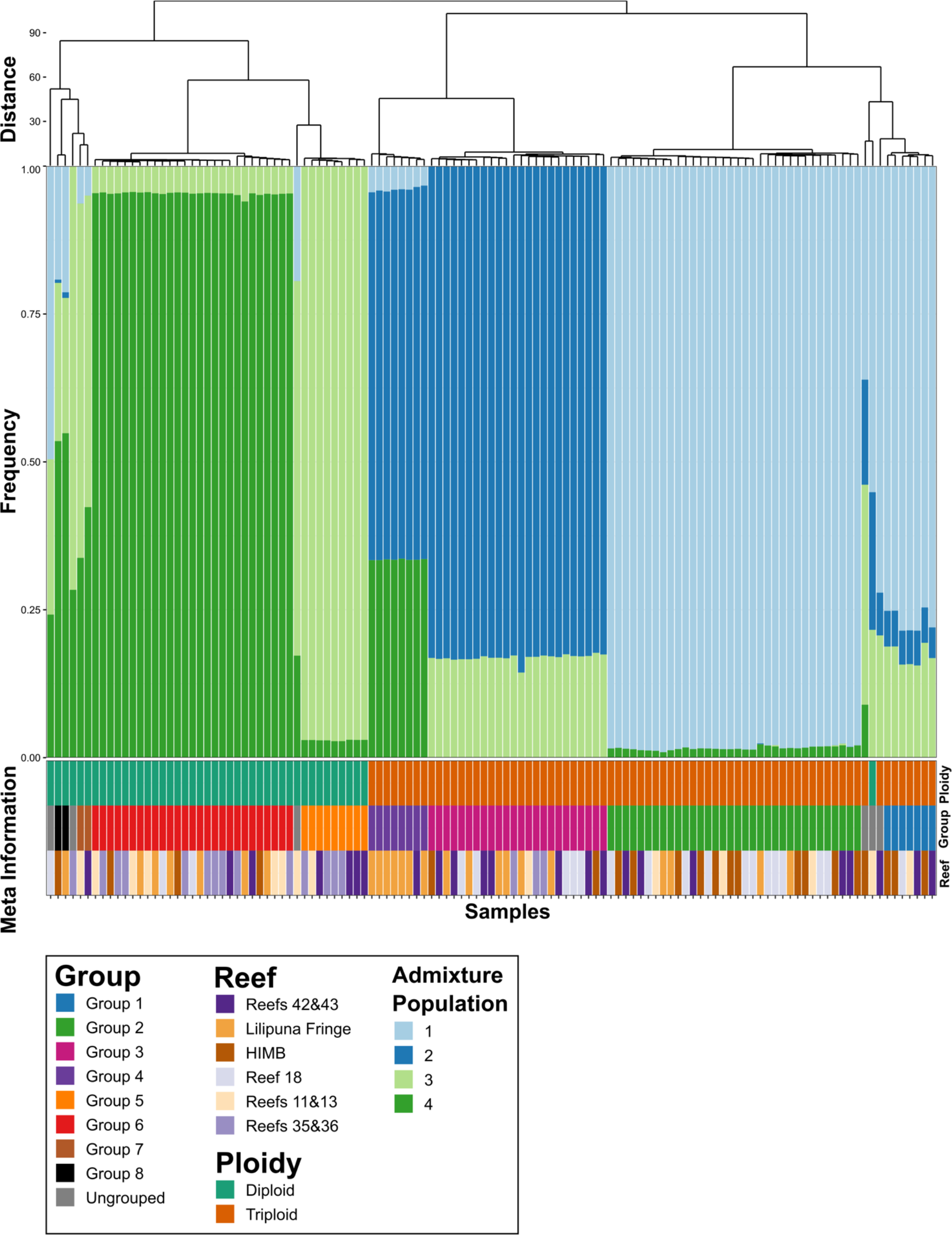
Admixture analysis of *P. acuta* samples. A stacked bar chart showing the proportion of each *P. acuta* sample’s ancestry that is from each of the four estimated (by PCAngsd) ancestral populations. The ploidy, putative clonal group, and reef that the sample was collected from are shown as colored bars at the bottom of the plot; a legend describing each of the colors used in the figure is presented at the bottom of the plot. The dendrogram and order of the columns is based on the hierarchical clustering of the proportion of shared SNPs presented in Figure 2.

**Fig. 5.**
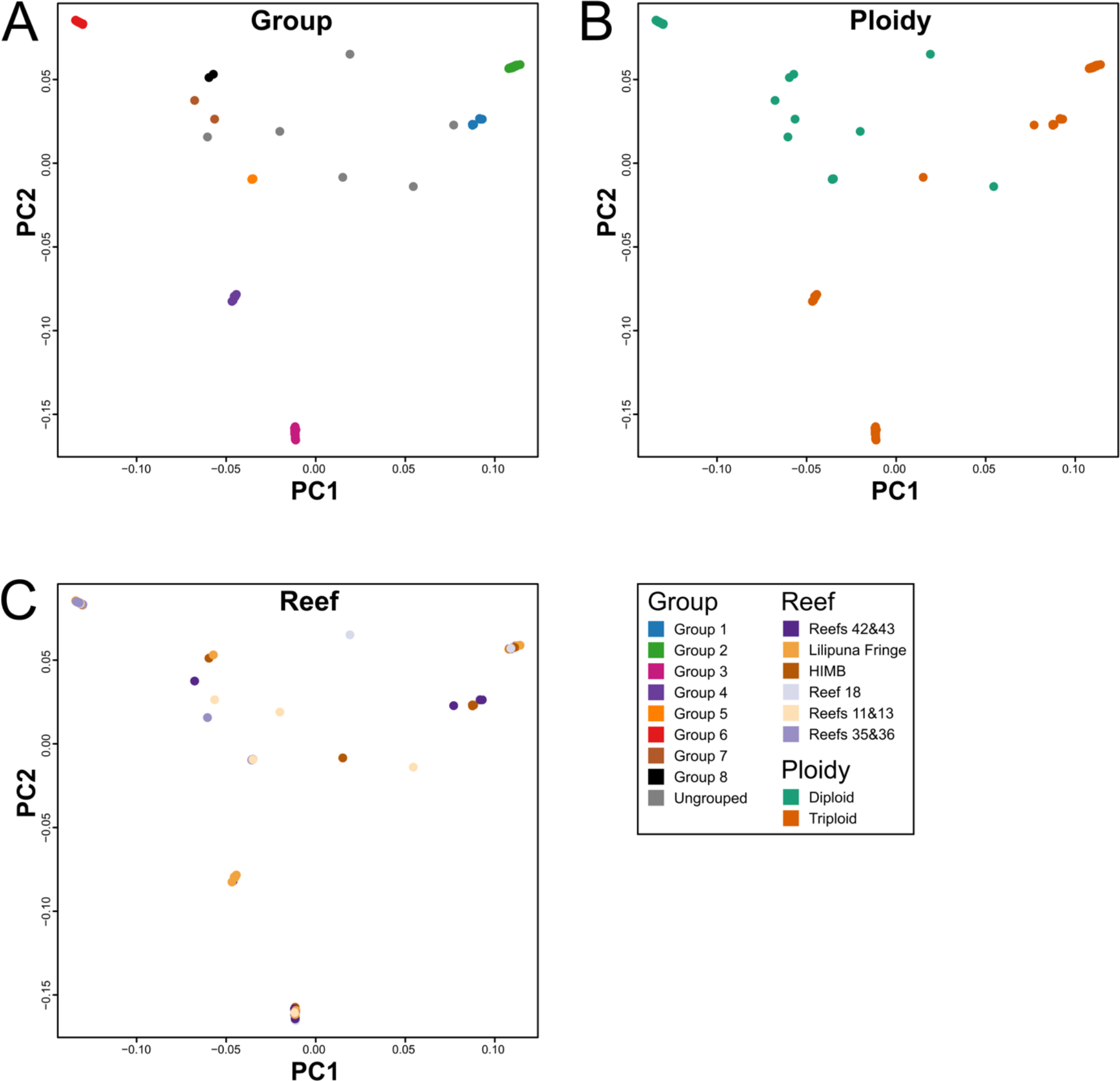
PCA of *P. acuta* sample relatedness. PCA of *P. acuta* samples colored by (**A**) group, (**B**) ploidy, and (**C**) reef. A legend describing each color is presented in the bottom right of the figure. Plots are based on the covariance matrix produced by PCAngsd with estimated individual allele frequencies and show PC1 (17.54% variance explained) and PC2 (14.66%).

### Population structure of *M. capitata* in Kāne‘ohe Bay

The same approaches used to analyze *P. acuta* were applied to the 132 *M. capitata* RNA-seq datasets. Here, we identified only two clonal *M. capitata* sampes (Groups 1, 2; fig. 6). Each clonal group comprises just two individuals: i.e., there were only four clonal samples (3% of the 132 samples analyzed). The overwhelming majority of samples had ∼85% pair-wise proportion of shared SNPs and < 0.15 relatedness (figs. 6 and 7). A significant number of samples have relatedness values of 0. Of the 27 samples of *M. capitata* downloaded from SRA, all were from the same project (BioProject: PRJNA377366; Frazier, et al. 2017), with some derived from different regions of the same colony (i.e., ramets). The patterns of relatedness and proportion of shared SNPs between the SRA samples and those sequenced in this study are the same as observed for *P. acuta*. That is, the two groups of *M. capitata* clonal samples from this study have similar values to the SRA ramet samples (Supplementary figs. S5 and S6; Supplementary tables S7 and S8). PCAngsd estimated two ancestral populations for Kāne‘ohe Bay *M. capitata* and overall, there were no obvious patterns in these results that would indicate strong population structure (Supplementary fig. S7). In addition, PCA performed by PCAngsd does not suggest a strong grouping of samples, with them distributed across PC1 (Supplementary fig. S8). Notably, the clonal samples from Group 1 separate from all other samples along PC2. These results all suggest that *M. capitata* in Kāne‘ohe Bay is a panmictic collection of sexual outbreeders, with very low relatedness between the analyzed samples.

**Fig. 6.**
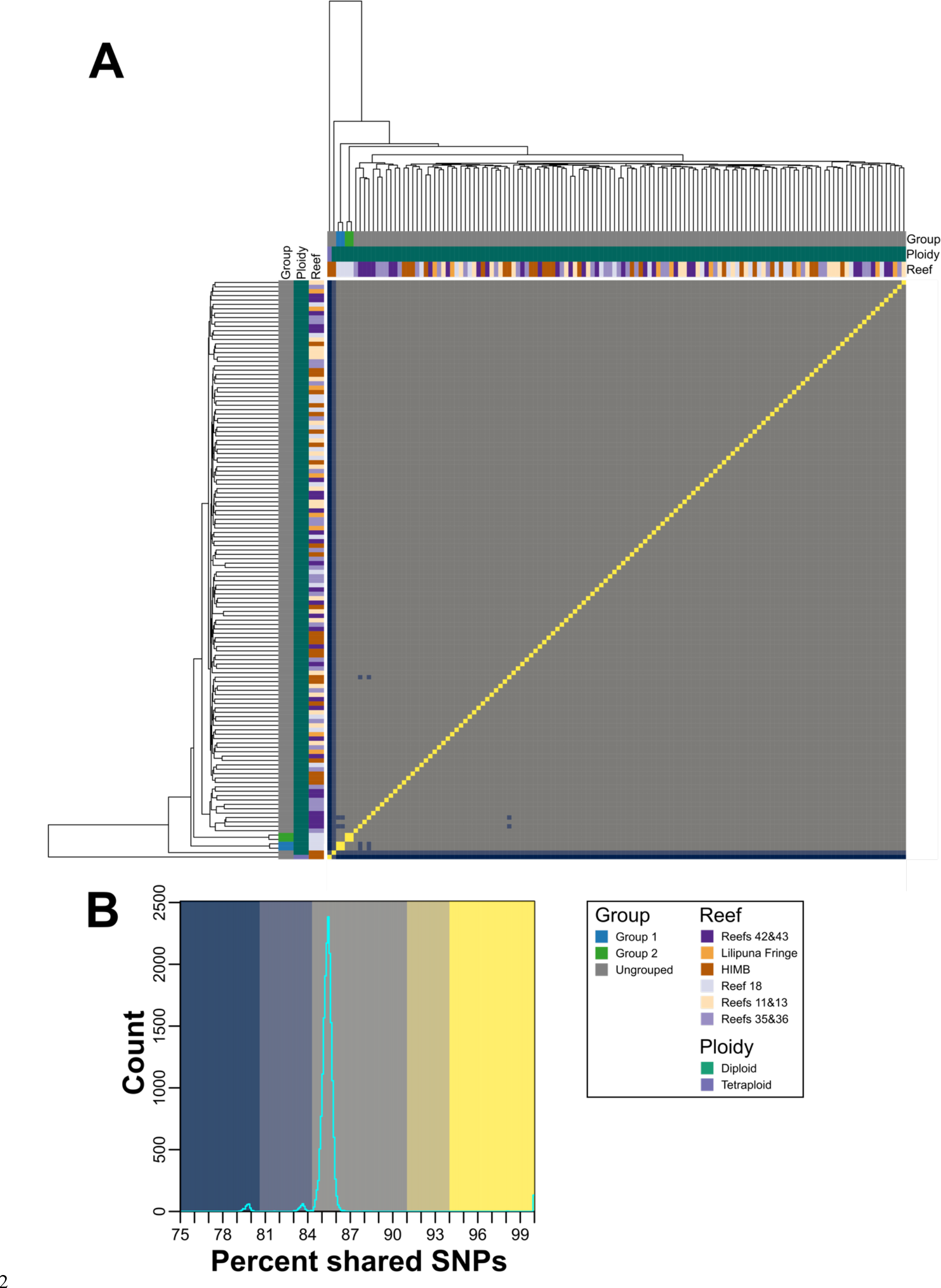
Proportion of shared SNPs between each combination of *M. capitata* samples. (**A**) Heatmap displaying the proportion of shared SNPs between all pair-wise combinations of *M. capitata* RNA-seq samples sequenced in this study. The putative clonal groups identified in this study (>94% shared SNPs) are shown as colored bars along the top and left sides of the heatmap; a legend describing the color of each group is presented on the bottom right side of the image. The order of the column and rows, and the dendrograms presented on the top and left sides of the heatmap, were generated by hierarchical clustering of the proportion of shared SNPs. (**B**) Histogram of the proportion of shared SNPs in the heatmap; the background colors used in the histogram correspond to the colors used in the heatmap presented in (**A**). The colors used in the heatmap and histogram were manually chosen to highlight the distinct sets of proportions of shared SNPs observed in the histogram presented in (**B**).

**Fig. 7.**
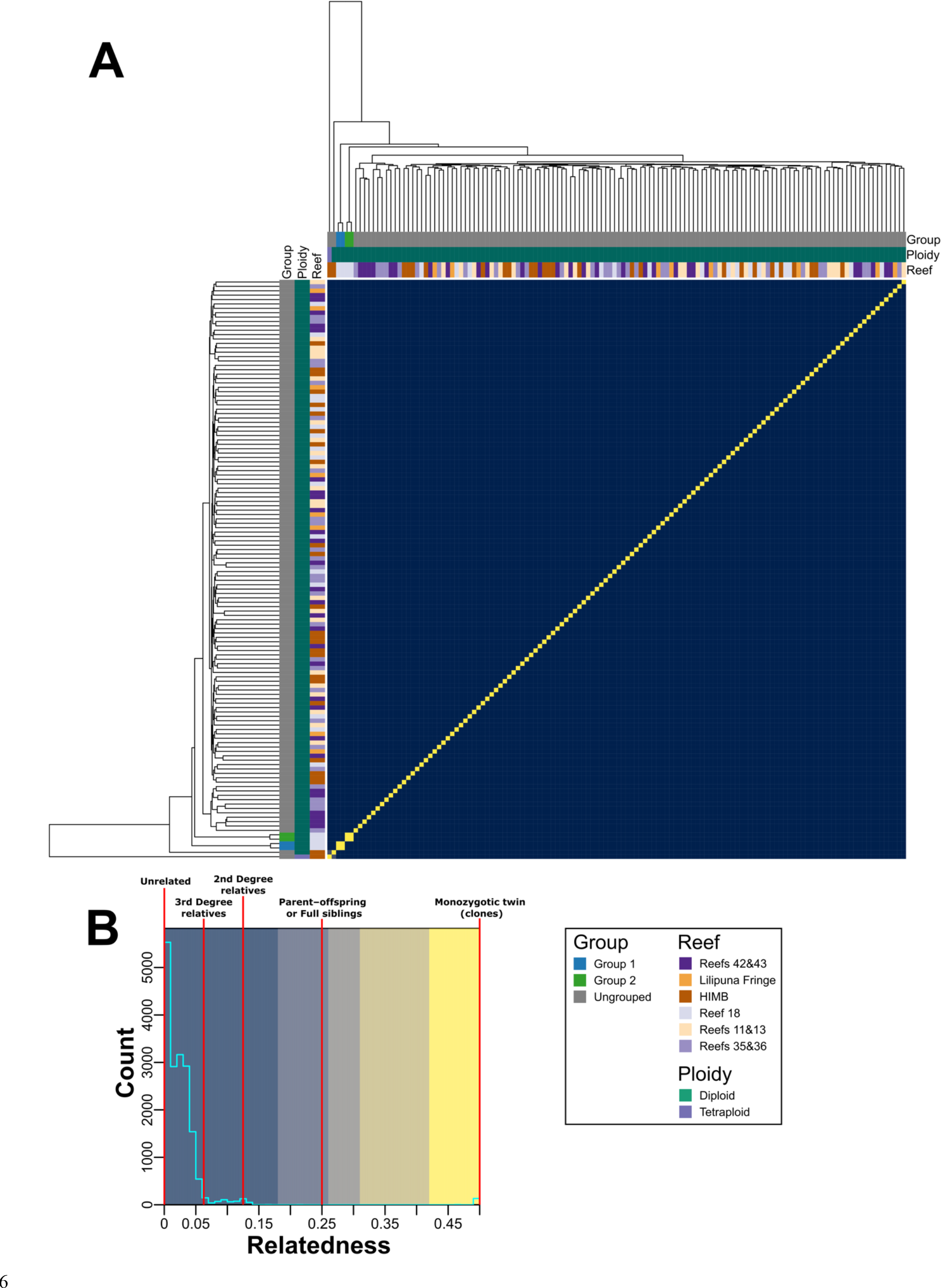
Pair-wise relatedness between each combination of *M. capitata* samples. (**A**) Heatmap displaying the relatedness scores (proposed by Manichaikul, et al. 2010) of all pair-wise combinations of *M. capitata* RNA-seq samples sequenced in this study. The putative clonal groups identified in this study are shown as colored bars along the top and left sides of the heatmap; a legend describing the color of each group is presented on the bottom right side of the image. The order of the column and rows, and the dendrograms presented on the top and left sides of the heatmap, are based on the dendrogram used in Figure 6. (**B**) Histogram of relatedness scores in the heatmap; the background colors used in the histogram correspond to the colors used in the heatmap presented in (**A**). The relatedness values that correspond to a particular relationship between samples (described in Manichaikul, et al. 2010) are annotated on the histogram using vertical red lines. The colors used in the heatmap and histogram were manually chosen to highlight the distinct sets of relatedness scores observed in the histogram presented in (**B**).

## Discussion

In this study, we generated RNA-seq data from colonies of *P. acuta* and *M. capitata* collected from six reefs distributed across Kāneʻohe Bay, Oʻahu, Hawaiʻi. We report significant differences in ploidy and reproduction strategies between the two sympatric species, with *P. acuta* derived from a mix of diploid and triploid clonal lineages, and *M. capitata* being a highly heterozygous, panmictic outbreeder.

### The adaptive advantage of triploidy in corals is unknown

The role of triploidy (or any form of polyploidy) in corals is not well understood. Although triploids are rare in wild populations, they occur frequently in commercially farmed plants and animals, such as oysters and some banana cultivars, often conferring beneficial commercial traits such as improved growth, pathogen resistance, and through infertility, protection of superior, adapted genotypes (Kang, et al. 2013). Triploids may also enhance the rate of autotetraploid formation (Husband 2004). Triploids occur in the coral *Acropora palmata* and may be a path to generating different ploidy levels in different members of this genus (Baums, et al. 2005; Kenyon 1997). Our results show that triploidy is common in Kāneʻohe Bay *P. acuta* (Supplementary fig. S1; Supplementary table S3) and has a higher abundance (63% at the sites sampled) than diploids (only 37%). This stands in clear contrast to *M. capitata*, which is completely (baring a single, possibly, chimeric sample) diploid. All methods for assessing sample relatedness (i.e., shared SNPs, relatedness metrics, PCA, and admixture analysis) predict that diploid samples are different from triploids, that is, there is clear separation between these groups (figs. 2, 3, 4, and 5). The only exception is a single diploid sample (Pacuta_HTHC_TP5_1415) that has higher similarity to triploid than diploid samples (although not high enough to be considered part of the closely related triploid Group 1). This individual could be an example of reversion (i.e., from triploid to a diploid), or be the extant member of the progenitor lineage of triploid Group 1. Our results suggest that the diploid *P. acuta* are both sexual outbreeders and generate asexual brooded larvae, as previously described (Combosch and Vollmer 2013; Gorospe and Karl 2013; Nakajima, et al. 2018; Richmond and Jokiel 1984; Schmidt-Roach, et al. 2014; Yeoh and Dai 2010). Triploidy may have arisen from self-fertilization of a *P. acuta* egg, followed by fertilization by a foreign sperm, or one of the two gametes was diploid and provided two closely related sets of alleles. Alternatively, failure of the ovum to extrude the second polar body after fertilization could lead to triploidy. These are the most common mechanisms for generating triploid plants and animals (Carson, et al. 2018; Rosenbusch 2008). The evolution of triploid genotypes in *P. acuta* may be explained by adaptation to local conditions in Kāne‘ohe Bay, possibly allowing them to outcompete ancestral diploid genotypes.

It is plausible that SNP allele frequency distributions, which are the basis of our estimation of ploidy, are explained by chimeric *P. acuta* colonies. Evidence exists for chimerism in corals through fusion of two or more genetically distinct individuals (Oury, et al. 2020; Rinkevich, et al. 2016; Willis, et al. 2006) as well as mosaicism *via* somatic cell mutations (Schweinsberg, et al. 2015; Willis, et al. 2006). Nonetheless, the two-peaked SNP distributions of *P. acuta* are difficult to explain under chimerism because the fused colonies would have to comprise roughly equal amounts of one haploid and one diploid individual to generate this result. Which is unlikely to have occurred in so many closely related colonies from across the bay. In addition, *k*-mer analysis of the reference triploid genome of *P. acuta* from Kāne‘ohe Bay (Stephens, et al. 2022) provides results that are consistent with our current findings. Given these results, we hypothesize that the most likely scenario to explain our data is triploidy in many *P. acuta* individuals, rather than fused/mixed diploids.

### Differences reproductive strategies of sympatric coral species

Our results also suggest that *P. acuta* in Kāne‘ohe Bay almost exclusively undergoes asexual reproduction, with only a few genets giving rise to colonies in the bay. This “genotypes everywhere” result has previously been found for Kāneʻohe Bay *P. damicornis* populations (Combosch and Vollmer 2013; Gorospe and Karl 2013; Yeoh and Dai 2010). Using microsatellite data, these authors studied a single reef and found that >70% of the colonies comprised seven genotypes with high clonal propagation.

Neighboring reefs however conformed to a genetic panmixia model with no inter-reef genetic structure. Our results support this model, showing the existence of at least eight groups of *P. acuta* samples (with each group representing a genet that has given rise to multiple independent colonies [ramets]) with broad distribution across Kāneʻohe Bay.

The presence of a limited number of genets in the bay, and the absence of isolation-by-distance, even when individual reefs show genetic structure is puzzling. This result may be explained by microhabitat variability that selects for particular genotypes that occupy specific niches in each reef (Gorospe and Karl 2011). These locally adapted genotypes disperse *via* asexual reproduction given that no major barriers exist for larval dispersal in Kāneʻohe Bay. This result might also be explained by a genetic bottleneck. A recent natural event, such as severe bleaching that caused mass coral mortality (e.g., the 2014- 2015 Kāneʻohe Bay bleaching event [Bahr, et al. 2015a]), may have removed much of the *P. acuta* from this region. The subsequent repopulation of Kāneʻohe Bay by surviving corals, or recolonization from different regions, coupled with asexual reproduction, would result in the observed, low genetic diversity. In addition, this would also explain why all the *P. acuta* samples analyzed in this study, even those not in the same clonal group, have relatively high relatedness. With the majority of samples (even between diploids and triploids) having relatedness values around 0.25 (i.e., the relatedness expected between parent and offspring or full sibling; fig. 3, Supplementary fig. S3, and Supplementary table S6). In contrast, the majority of relatedness values of *M. capitata* samples were close to zero, which indicates unrelated individuals (fig. 7, Supplementary fig. S6, and Supplementary table S8). This result for *M. capitata* has been previously reported (Caruso, et al. 2021; Concepcion, et al. 2014). A recent survey of nearly 600 colonies of this species in Kāneʻohe Bay found very few clonal individuals and no evidence of isolation by distance (Caruso, et al. 2021). Colonies that were potentially derived from the same genet were almost exclusively found at the same collection site, consistent with our observations.

### Study limitations

Our study makes extensive use of RNA-seq data that was originally generated as part of a mesocosm experiment not relevant to the results of this research. Whereas RNA-seq data are not commonly used in population genetics, in this case, we believe that it provides invaluable insights into coral biology that can inform follow-up DNA-based sequencing projects. We acknowledge however that ploidy is more challenging to interpret using RNA-seq data. We have previous described, using DNA sequencing data, a triploid *P. acuta* genet from Kāneʻohe Bay (Stephens, et al. 2022), which was included in this study and was identified using RNA-seq data as a triploid. To the best of our knowledge, all bioinformatic approaches for ploidy determination (such as nQuire and visualization of allele frequencies, which were used by this study) are designed for use with DNA, not RNA data. Thus, we cannot fully discount allele specific expression (ASE) as an alternative explanation for the patterns that we observe. However, we believe it is unlikely that ASE has affected our results, for the following four reasons: (1) we have clear DNA evidence for triploidy from one of the samples (Stephens, et al. 2022). (2) If ASE is affecting our results, it would have to be strongly affecting some groups of sample and not others (i.e., ASE is only occurring in putative triploid and not diploid lineages, and not in the triploid with DNA evidence). (3) ASE typically occurs at different rates across the genome, that is, ASE produces an uneven distribution of expression ratios. Our results suggest that all loci are being affected at the same rate, which would support variation in chromosome copy number and not locus-specific allelic expression modification. And (4), coral genomes have relatively low rates of methylation (Trigg, et al. 2022), with 11.4% of CpG sites in the *M. capitata* genome being methylated, with this value being 2.9% in *P. acuta*. Given that methylation would be the most obvious mechanism for ASE, the low methylation rate in *P. acuta* makes ASE a less likely explanation for the ploidy results.

Regarding the analysis of samples with mixed ploidy, few of the available population genetic techniques accept non-diploid data, and none that we are aware of accept data with mixed ploidies. As a result, all samples were treated as diploid, which may adversely affect results from the putative triploid samples because it would bias our analysis to just biallelic sites. However, given that we expect most variant sites in the genome to be biallelic (because multiple mutations occurring at a single site to create a multiallelic site is less likely than a single mutation to create a biallelic site), we believe that our approach is valid given the current techniques and data available. Furthermore, given that a variety of data analysis tools were used, and all led to the same conclusions, we believe our results to be robust. These insights should prove valuable for designing DNA-based studies that focus not only on generating additional population genetic data from Kāneʻohe Bay, but also from other locations in the Hawaiian Islands. There is currently very little data available for *P. acuta*, preventing us from comparing our results to other populations or studies done in this region.

### Final remarks

The data presented in this study underline how selection can act in a divergent manner to forge ecologically successful lineages. We find that two sympatric species living in a sheltered Hawaiian bay that has been severely impacted by human activity, including warming events, freshwater incursion, and impacts of dredging (Bahr, et al. 2015b), follow diametrically opposed strategies to allow persistence. *M. capitata* relies on strict outbreeding to generate high standing genetic variation as a “defense” against changing local environments. In contract, *P. acuta* relies on periodic polyploidization events, perhaps triggered by local stress, to putatively generate fitter, clonal groups to reestablish populations in Kāneʻohe Bay through asexual reproduction. Next steps to expand our understanding of these patterns include studying the response of Hawaiian corals with divergent genotypes to environmental stress, to ascertain the connection between the two.

## Materials and Methods

### Sample processing

The coral samples (one ∼5x5cm fragment per colony) were collected from six reef areas ranging across the north to south span and fringing to patch reefs of Kāneʻohe Bay (Lilipuna Fringe: 21°25’45.9”N 157°47’28.0”W; HIMB: 21°26’09.8”N 157°47’12.7”W; Reef 18: 21°27’02.9”N 157°48’40.1”W; Reef 11, 13: 21°27’02.9”N 157°47’41.8”W; Reef 35, 36: 21°28’26.0”N 157°50’01.2”W; Reef 42, 43: 21°28’37.9”N 157°49’36.8”W; fig. 1C) under Hawaiʻi Department of Aquatic Resources Special Activity Permit 2019-60, between 4-10 September 2018. RNA was extracted from the snap frozen nubbins and stored at -80 °C. A small piece was clipped off using clippers sterilized in 10% bleach, deionized water, isopropanol, and RNAse free water, and then placed in 2 mL microcentrifuge tube containing 0.5mm glass beads (Fisher Scientific Catalog. No 15- 340-152) with 1000 μL of DNA/RNA shield. A two-step extraction protocol was used to extract RNA, with the first step as a “soft” homogenization to reduce shearing RNA. Tubes were vortexed at high speed for 1 and 2 minutes for *P. acuta* and *M. capitata* fragments, respectively. The supernatant was removed and designated as the “soft extraction”. Second, 500 μL of DNA/RNA shield was added to the bead tubes and placed in a Qiagen TissueLyser for 1 minute at 20 Hz. The supernatant was removed and designated as the “hard extraction”. Subsequently, 300 μL of sample from both soft and hard homogenate was extracted with the Zymo Quick-DNA/RNA Miniprep Plus Kit. RNA quality was measured with an Agilent TapeStation System. RNA-Seq samples were sequenced by GENEWIZ (Azenta; https://www.genewiz.com) using the Illumina NovaSeq 6000 platform.

### Statistical Analysis

#### RNA base quality and adapter trimming

Adapters and low-quality regions were trimmed from the RNA-seq data generated in this study using Cutadapt v2.9 (Martin 2011) (--nextseq-trim 10 --minimum-length 25 -a AGATCGGAAGAGCACACGTCTGAACTCCAGTCAC -A AGATCGGAAGAGCGTCGTGTAGGGAAAGAGTGTA; Supplementary table S2). A second round of trimming with Cutadapt, using the output from the first round, was used to remove poly-G regions from the 5′-ends of the second read in each pair (-G G(20) -e0.0 -n 10 --minimum-length 25). Read quality was assessed at each stage using FastQC v0.11.7 (default parameters; http://www.bioinformatics.babraham.ac.uk/projects/fastqc/) and MultiQC (Ewels, et al. 2016) (v1.9).

#### Alignment of RNA-Seq data against reference genomes

RNA-seq reads were aligned against the *M. capitata* (Version 3; http://cyanophora.rutgers.edu/montipora/; Stephens, et al. 2022) and *P. acuta* (Version 2; http://cyanophora.rutgers.edu/Pocillopora_acuta/; Stephens, et al. 2022) reference genomes following the GATK (v4.2.0.0) (McKenna, et al. 2010) framework, adhering to their best practices workflow (https://gatk.broadinstitute.org/hc/en-us/articles/360035531192?id=4067) whenever possible. For each sample, the Cutadapt trimmed RNA-Seq reads were aligned against the appropriate reference genome using STAR (v2.7.8a; --sjdbOverhang 149 --outSAMtype BAM SortedByCoordinate -- twopassMode Basic) (Dobin, et al. 2013). The trimmed reads were converted into unaligned-SAM (uSAM) format using gatk FastqToSam (setting --SAMPLE_NAME to be the name of the input SAM file); read-group information was extracted from the read names from the uSAM file by rgsam (v0.1; https://github.com/djhshih/rgsam; --qnformat illumina-1.8) and added to the aligned reads using gatk MergeBamAlignment (-- INCLUDE_SECONDARY_ALIGNMENTS false --VALIDATION_STRINGENCY SILENT).

Duplicate reads (that had originating from the same DNA fragment) were identified and annotated using gatk MarkDuplicates (--CREATE_INDEX true -- VALIDATION_STRINGENCY SILENT) before reads that spanned intron-exon boundaries were split using gatk SplitNCigarReads (default parameters). The resulting BAM files were used as the input for downstream population structure, sample relatedness, and ploidy analysis.

#### Processing of coral data not generated in this study

Additional RNA-seq samples were acquired on 15th June 2022 from NCBI’s SRA database for use as outgroups in downstream analysis. A list of all sequencing “Runs” from Scleractinian species were acquired by searching the NCBI’s SRA database using the following search term: “Scleractinia[Organism]” (without the double quotes). The resulting list of 19,050 entries was filtered, keeping only Runs that were generated on an Illumina platform, that had a library strategy of “RNA-Seq”, a library layout of “PAIRED”, and >1.5 billion “bases”. These filters were chosen to keep the types of samples selected from SRA uniform with the samples generated in this study (i.e., paired-end RNA-seq reads generated on an Illumina platform). The 1.5 Gbp threshold was chosen as it is roughly half the minimum number of bases (∼3.2 Gbp; Supplementary table S2) generated for a single sample from this study before base quality filtering was applied; samples with less than this number of bases would likely have far fewer sites with sufficient coverage for downstream analysis so would be less informative and harder to interpret and integrate with the existing samples if included. Sample derived from colonies identified as *P. acuta* and *M. capitata* were extracted from the resulting list of filtered Runs using the species name listed in the “ScientificName” column. The “geographic_location” of each Run was extracted from it associated BioSample and used to identify Runs generated from samples collected from Hawaiʻi. Runs generated from colonies collected in Kāneʻohe Bay were excluded, as were Runs without a listed geographic location. As there were no Runs listed on SRA of Hawaiian *P. acuta* colonies that were not from Kāneʻohe Bay. Runs from other non-Hawaiian locations were selected. This resulted in a total of 27 *M. capitata* and 32 *P. acuta* Runs (samples) that were used for downstream analysis (Supplementary table S4).

The selected samples were retrieved from SRA using fasterq-dump (v2.9.6-1; https://github.com/ncbi/sra-tools) and trimmed using Cutadapt v2.9 (-q 10 --minimum-length 25 -a AGATCGGAAGAGCACACGTCTGAACTCCAGTCA -A AGATCGGAAGAGCGTCGTGTAGGGAAAGAGTGT). The samples were aligned against the *P. acuta* (V2) or *M. capitata* (V3) reference genomes using the same workflow as the RNA-Seq samples generated in this study, with the only exception being that read-group information was added manually to the reads (using gatk AddOrReplaceReadGroups; setting read-group platform to be “illumina” and the read- group library, platform unit, sample name, and ID to be the SRA ID of the sample) instead of using rgsam to extract the information automatically from the read names.

This was done because read-group information needs to be set for some of the downstream tools (such as gatk MarkDuplicates) but this information is not required or preserved during upload of read data to SRA, so cannot be reliably extracted from the downloaded samples. The resulting BAM files were used as the input for downstream population structure, sample relatedness, and ploidy analysis.

#### Identification of variant sites in each sample

Variants (single-nucleotide polymorphisms [SNPs] and insertion and deletion variants [INDELs]) were identified across the RNA-seq samples from each species (both generated in this study and from SRA) using the GATK (McKenna, et al. 2010) (v4.2.0.0) framework, adhering to their best practices workflow (https://gatk.broadinstitute.org/hc/en-us/articles/360035531192?id=4067) whenever possible. The base quality recalibration steps suggested in the GATK best practices workflow could not be applied to our samples because a set of expected high confidence variants (which is required for the recalibration process) is not available for either of the species being studied. Haplotypes were called using the aligned post-processed reads (i.e., after alignment using STAR with read-group information added, duplicate reads marked, and reads spanning exon boundaries split) using gatk HaplotypeCaller (-dont-use-soft-clipped-bases -ERC GVCF). For each species, the VCF files produced by HaplotypeCaller (one per sample) were combined using gatk CombineGVCFs before being jointly genotyped using gatk GenotypeGVCFs (-stand-call-conf 30 --annotation AS_MappingQualityRankSumTest --annotation AS_ReadPosRankSumTest). For the gatk analysis, the ploidy of each sample was treated as diploid (even when the samples were believed to be triploid). This was done because the downstream tools are incapable of processing triploid variants or datasets with mixed diploid and triploid variants.

#### Analysis of sample ploidy

The ploidy of the RNA-Seq samples generated in this study and acquired from SRA were assessed using nQuire (Weiss, et al. 2018) (retrieved on 7/7/2021 from https://github.com/clwgg/nQuire), which was run using the BAM files produced by gatk SplitNCigarReads (i.e., aligned RNA-seq reads that have had duplicates removed and that have been split if they span an intron-exon boundary). The aligned BAM files were converted into “BIN” files, filtering for reads with a minimum mapping quality of 20 and sites with a minimum coverage of 20 (“nQuire create -q 20 -c 20 -x”). Denoised BIN files were created using the “nQuire denoise” command run on the initial BIN files. The delta Log-Likelihood values for each ploidy model was calculated by the “nQuire lrdmodel” command for each of the initial and denoised BIN files (Supplementary table S3). A second round of analysis using nQuire was conducted to generate, for each sample, a distribution of the proportion of reads which support each allele at biallelic sites. Briefly, “BIN” files were created, filtering for reads with a minimum mapping quality of 20, sites with a minimum coverage of 20, and a minimum fraction of reads supporting an allele of 5% (“nQuire create -f 0.05 -q 20 -c 20 -x”). Denoised BIN files were created using the “nQuire denoise” command. The “nQuire view” command was used to extract the biallelic sites from the denoised BIN files. The number of aligned reads reported by nQuire that support each of the alleles at the biallelic sites were used to generate a distribution, for each of the samples, of the proportion of reads which support each of the alleles; this distribution was used to visually confirm the ploidy of each sample (Supplementary data S1, S2, S3, and S4).

#### Exploration of population structure

The population structure of the *P. acuta* and *M. capitata* samples collected from Kāne‘ohe Bay and SRA was assessed using multiple approaches. The genotype likelihoods of the samples from each species were estimated by ANGSD (v0.935; “-GL 2 -doGlf 2 -doMajorMinor 1 -SNP_pval 1e-6 -doMaf 1”) (Korneliussen, et al. 2014) and used with PCAngsd (v1.10) (Meisner and Albrechtsen 2018) to perform Principle Component Analysis (PCA; with estimates for individual allele frequencies; default parameters), with the resulting covariance matrix used for visualization. The genotype likelihoods produced by ANGSD were also used with PCAngsd to perform an admixture analysis (“--admix --admix_alpha 50”) of the samples from each species.

The variants produced by “gatk GenotypeGVCFs” were filtered using vcftools (v0.1.17; “--remove-indels --min-meanDP 10 --max-missing 1.0 --recode --recode-INFO-all”) to remove indels, variants with low average reads coverage across all samples, and sites which do not have called genotypes across all samples. A minor allele frequency (MAF) threshold was not applied to the data, because we knew *a priori* from preliminary analysis of the data that some of the *P. acuta* clonal groups are comprised of very few samples. That is, the use of a MAF (e.g., 0.05, which would remove alleles that appear in < 5% of samples) would disproportionately remove variants that segregate the small clonal groups of samples (e.g., *P. acuta* Groups 7 & 8 which are each comprised of two samples [1.7% of the 119 samples]). This would reduce our ability to resolve the smaller groups of clonal samples in our dataset, biasing our analysis towards resolving only larger clonal groups. By not applying a MAF threshold we increase the chances of incorporating false positive variants into our analysis, however, this would likely only slightly reduce the relatedness between the samples and obscure their ploidy. Thus, it would only result in increased noise in the data, not inflation of sample relatedness or ploidy. Vcftools (“--relatedness2”) was also used to compute the relatedness statistic developed by Manichaikul, et al. 2010. Negative relatedness values produced by vcftools were converted to zero before downstream analysis. The number of SNPs shared between each pair of samples from each species was assessed using the “vcf_clone_detect.py” script from https://github.com/pimbongaerts/radseq (retrieved 12/06/2022), which was run using the vcftools filtered VCF file. A threshold of >94% shared SNPs, chosen based on the distribution of shared SNPs across all pair-wise combinations of samples (fig. 2), was used to aggregate samples into groups that were used for all downstream analysis.

For each species, two sets of analyses using ANGSD, PCAngsd, vcftools, and “vcf_clone_detect.py” were performed, one using only the RNA-seq samples produced in this study and the other using the RNA-seq samples from this study and the samples downloaded from SRA.

## Acknowledgments

This work was supported by the USDA National Institute of Food and Agriculture Hatch Formula project accession number 1017848 awarded to HMP; USDA National Institute of Food and Agriculture Hatch Formula project accession number NJ01180 awarded to DB; National Science Foundation grant NSF-OCE 1756616 awarded to DB; Catalyst Science Fund grant 2020-008 awarded to DB; National Aeronautics and Space Administration grant 80NSSC19K0462 awarded to DB; National Science Foundation grant NSF-OCE 1756623 awarded to HMP. We acknowledge support of the research and administrative staff at the Hawaiʻi Institute of Marine Biology where this work was done.

## Author contributions

Conceptualization: DB, HMP, TGS

Methodology: TGS, ES

Supervision: DB, HMP

Writing—original draft: TGS, DB

Writing—review & editing: TGS, EM, HMP, DB

## Data availability

The RNA-Seq data generated in this study are available from NCBI’s SRA repository (PRJNA731596). The code used to analyze the data is available in the GitHub repository (https://github.com/TimothyStephens/Kaneohe_Bay_coral_2018_PopGen.git). The genome assemblies and predicted genes used in this study are available from http://cyanophora.rutgers.edu/montipora/ (Version 3)and http://cyanophora.rutgers.edu/Pocillopora_acuta/ (Version 2). All other data needed to evaluate the conclusions in the paper are present in the paper and/or the Supplementary Materials.

---

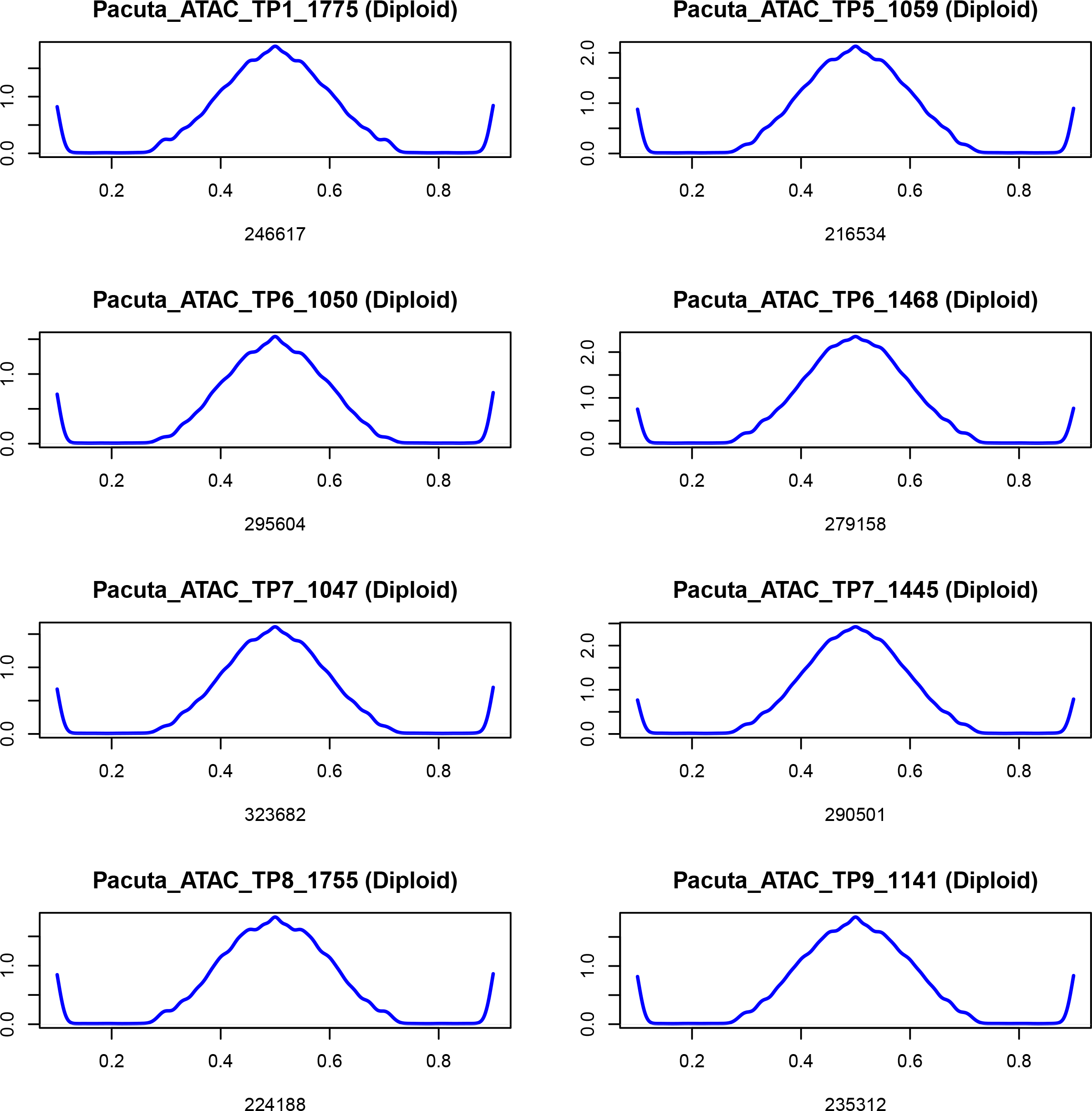
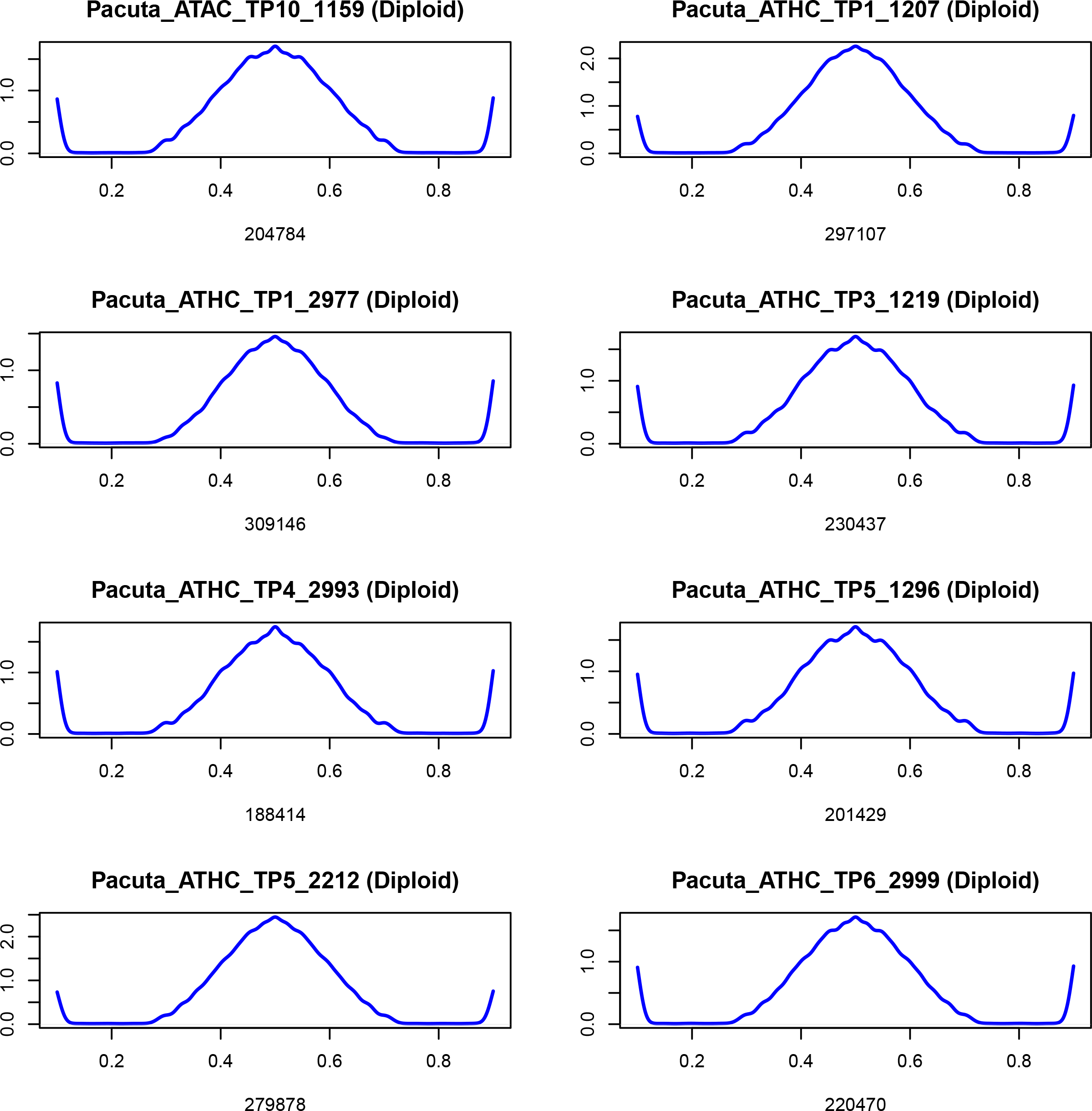
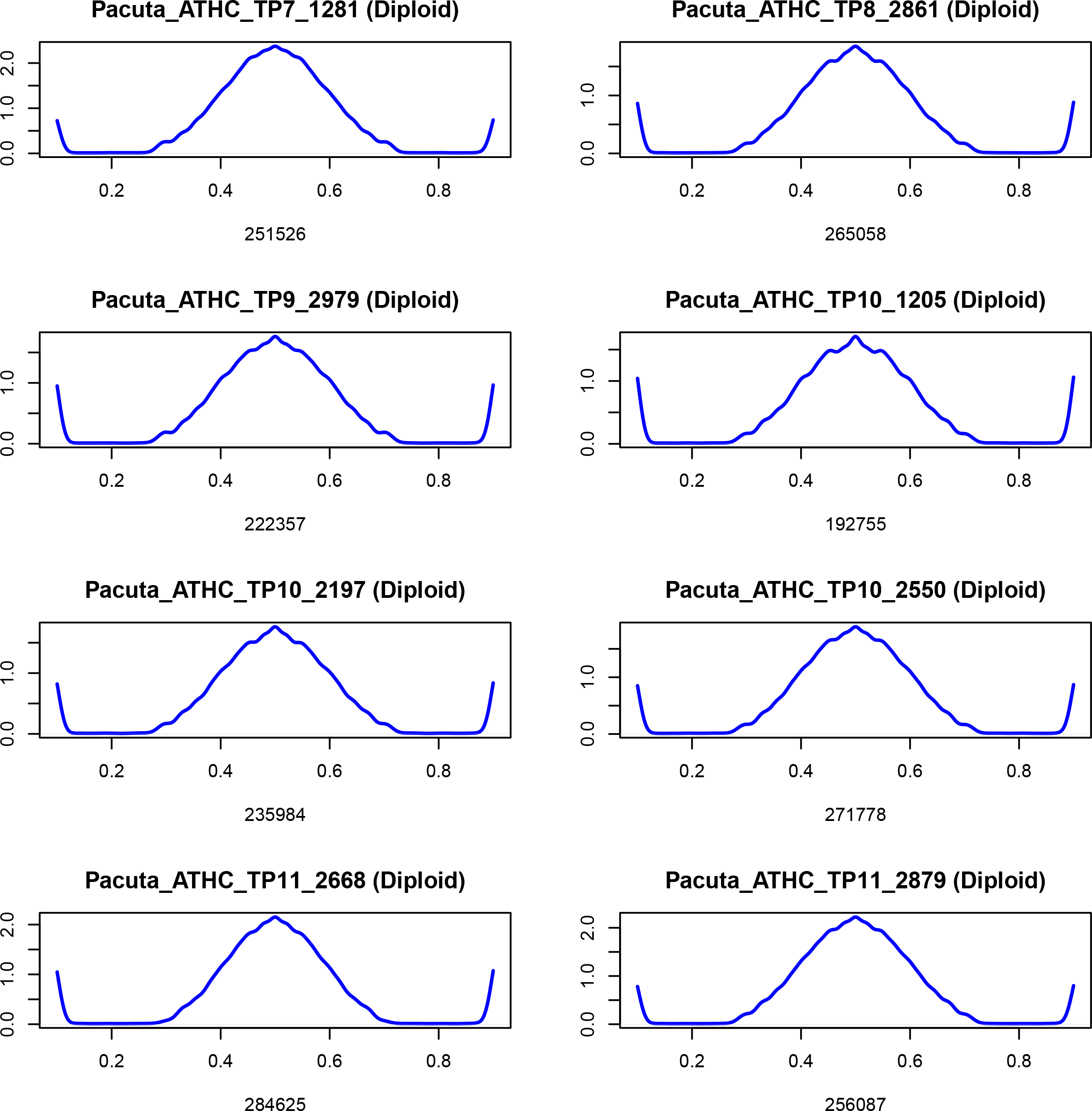
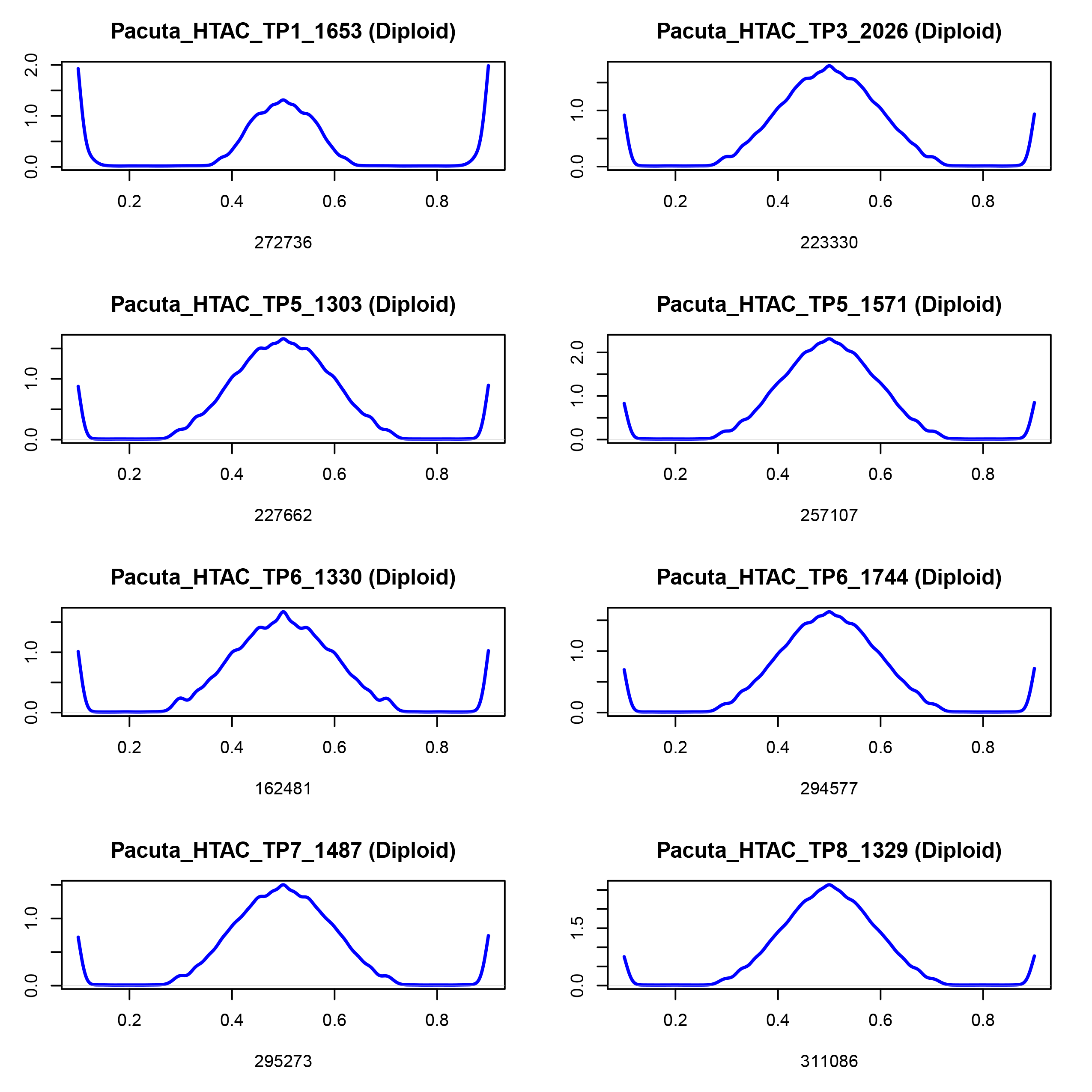
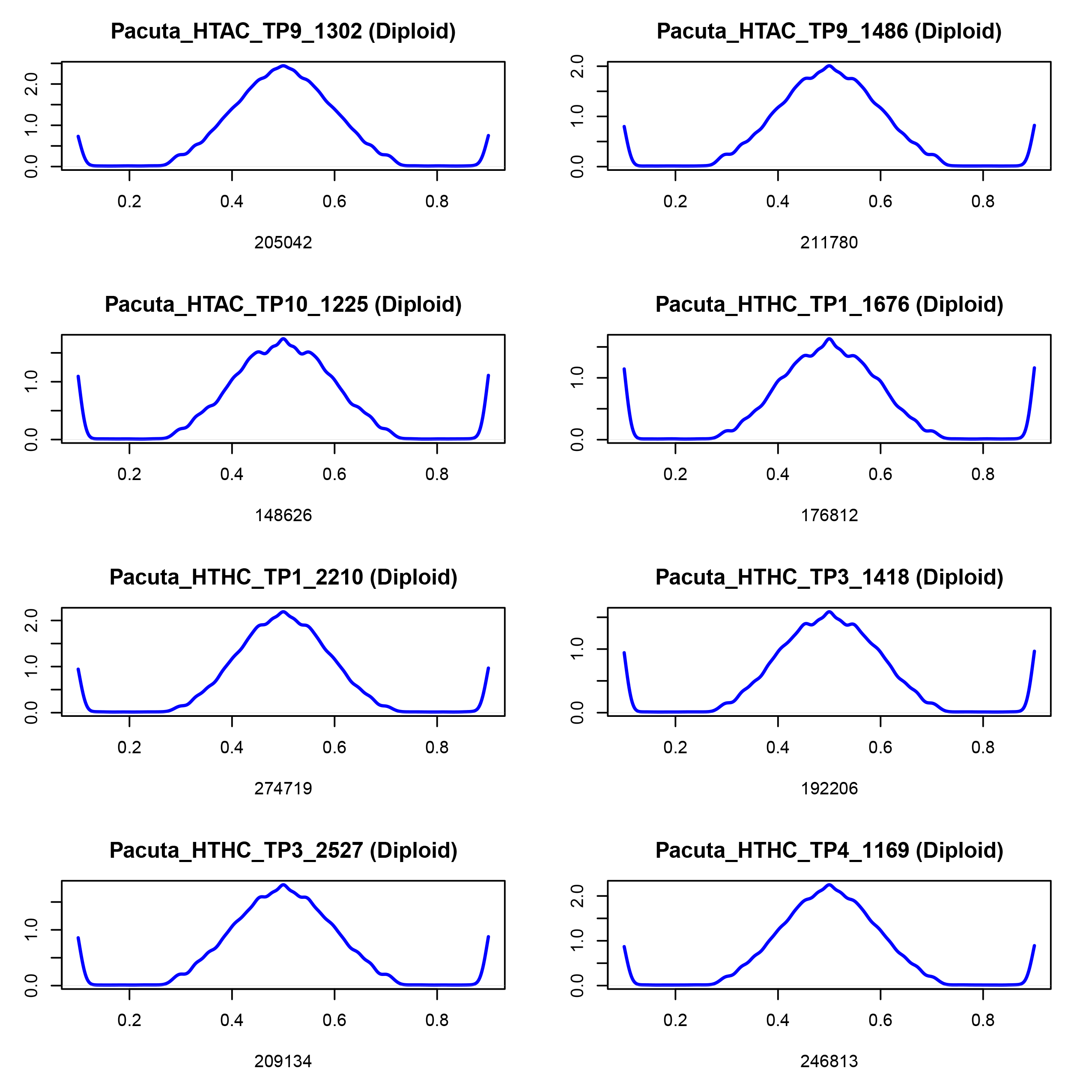
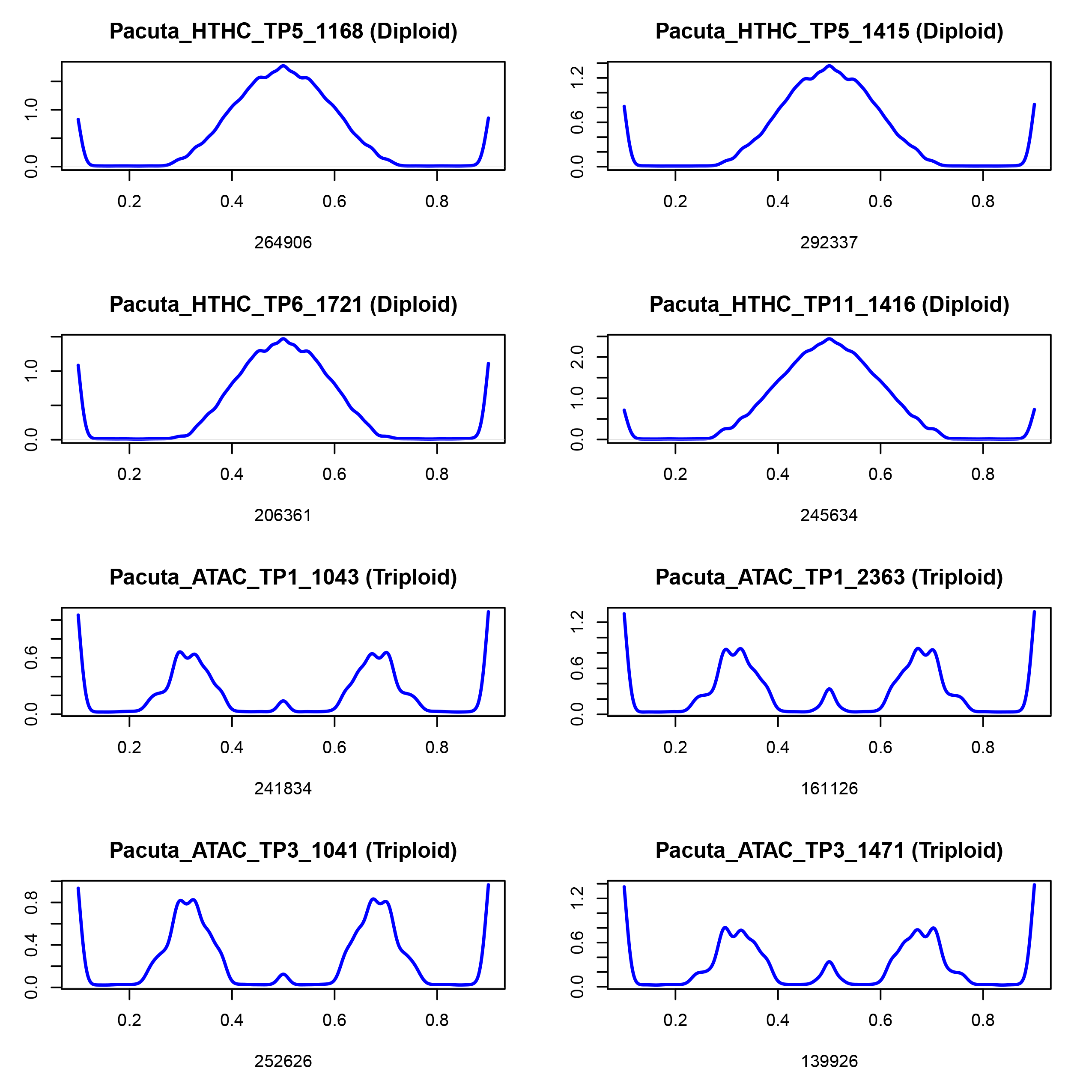
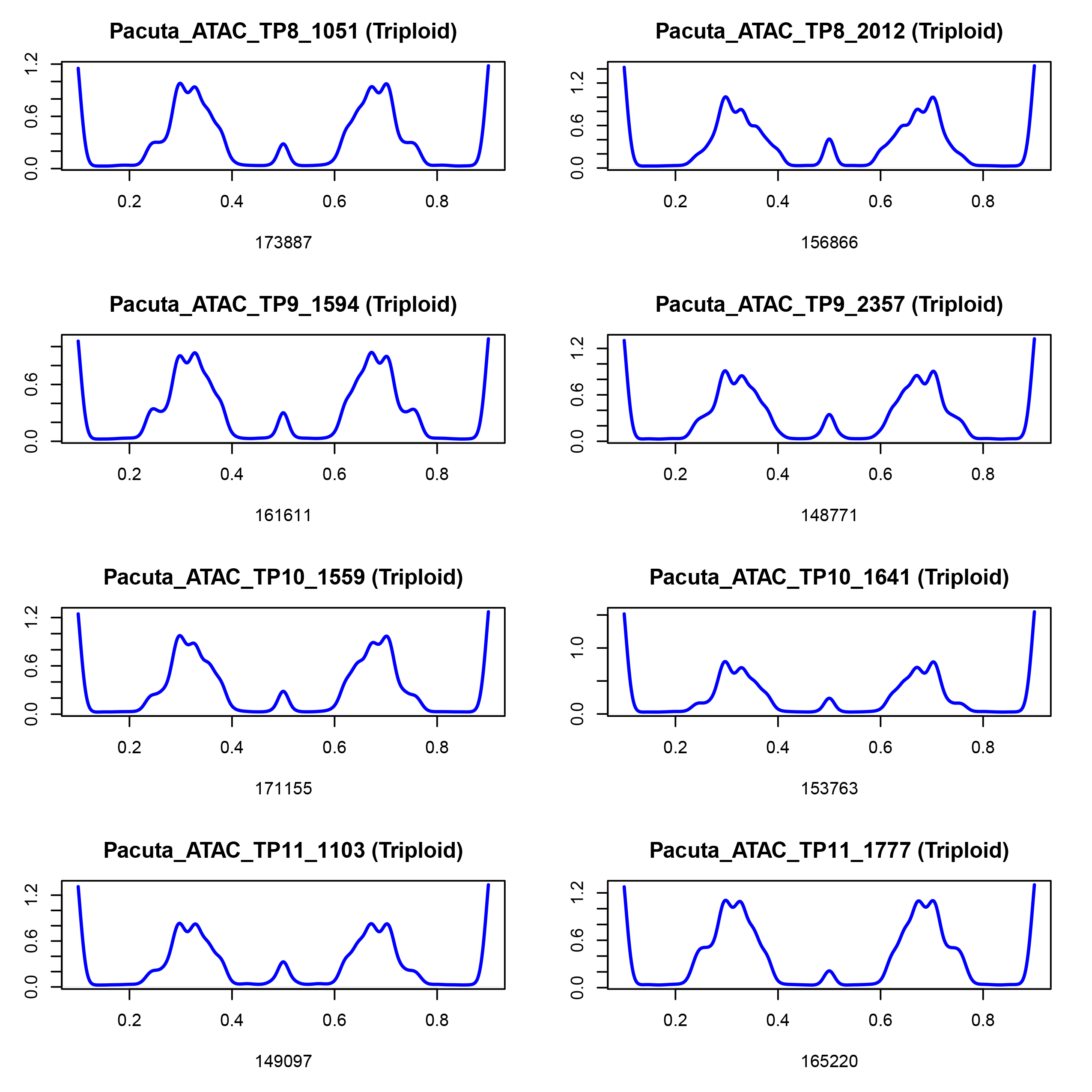
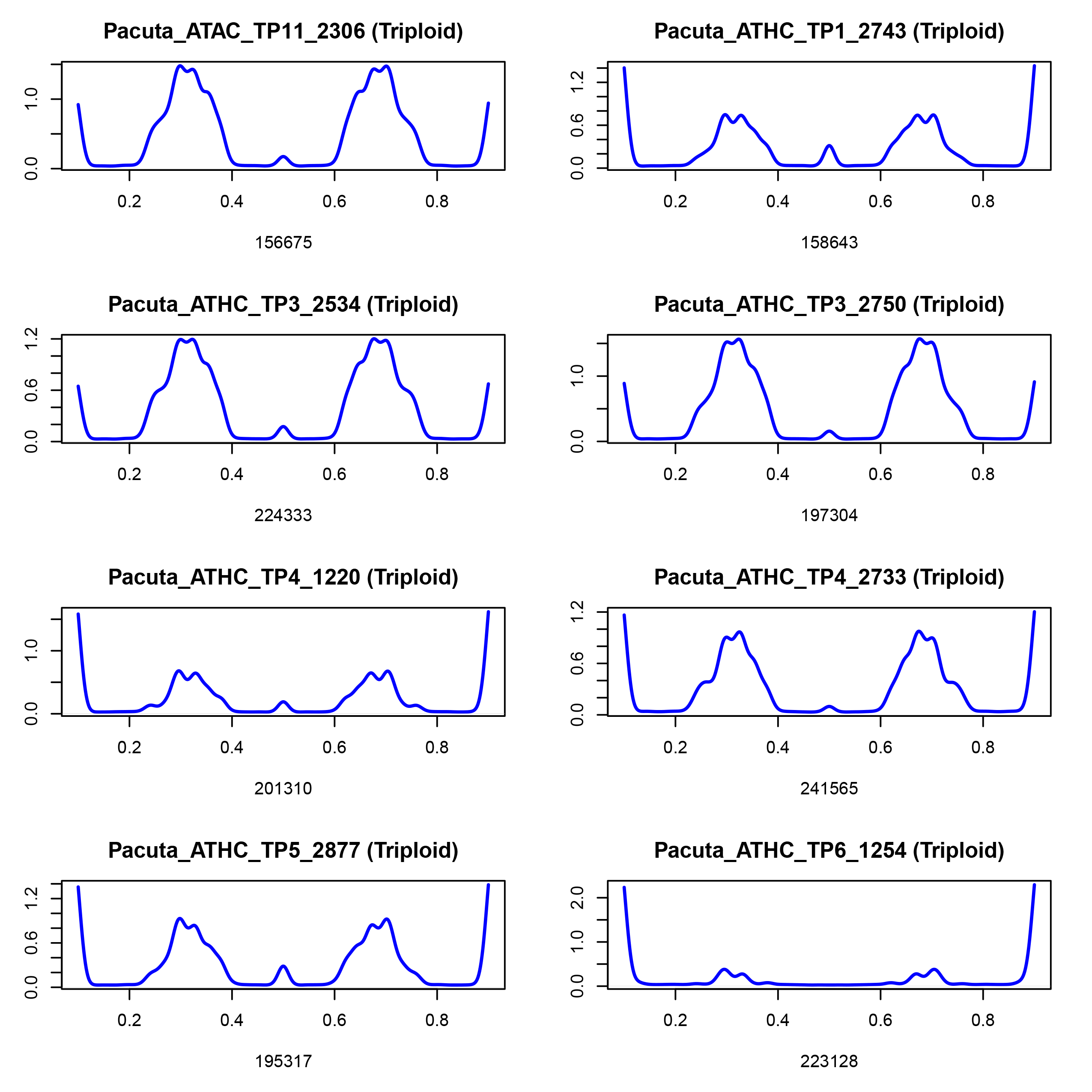
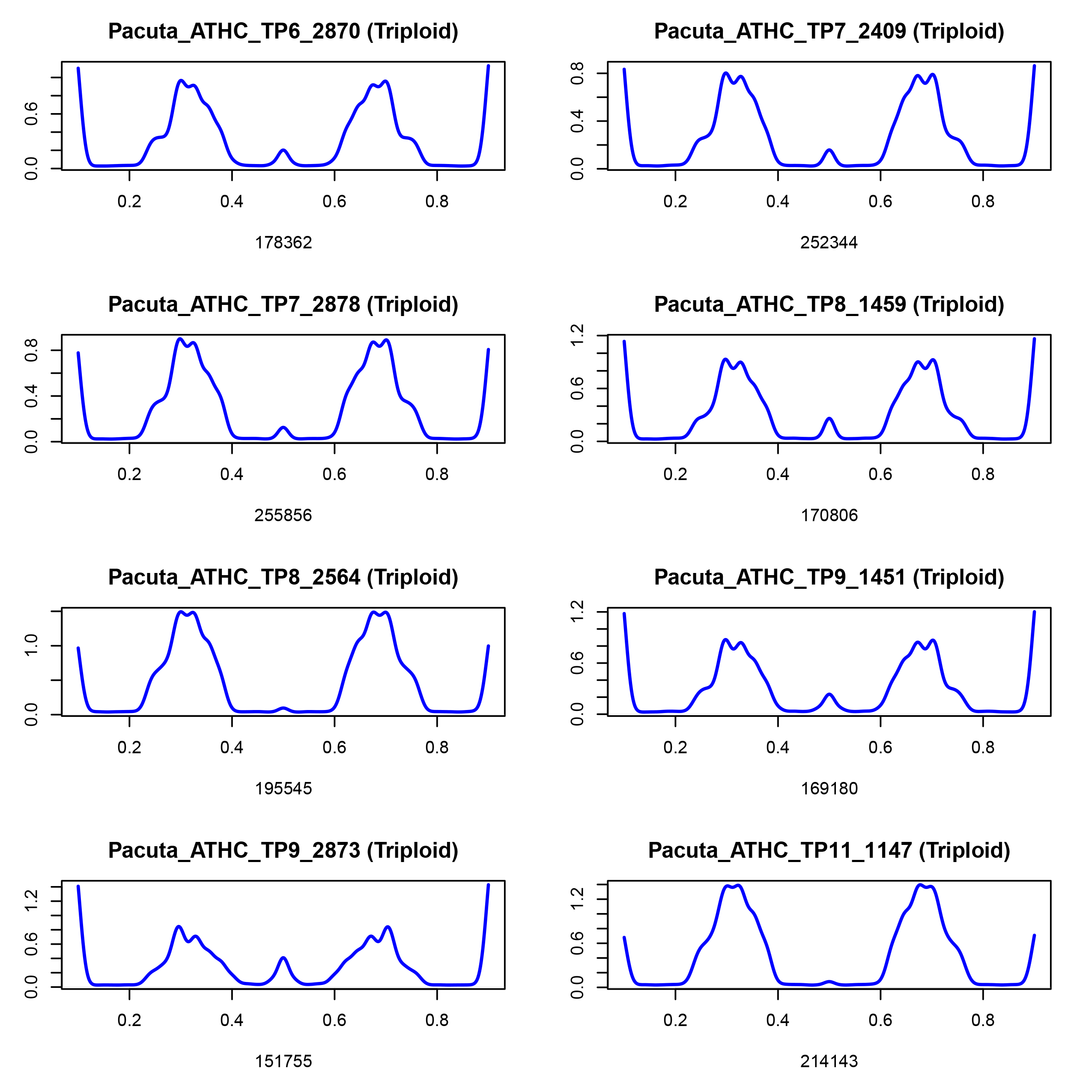
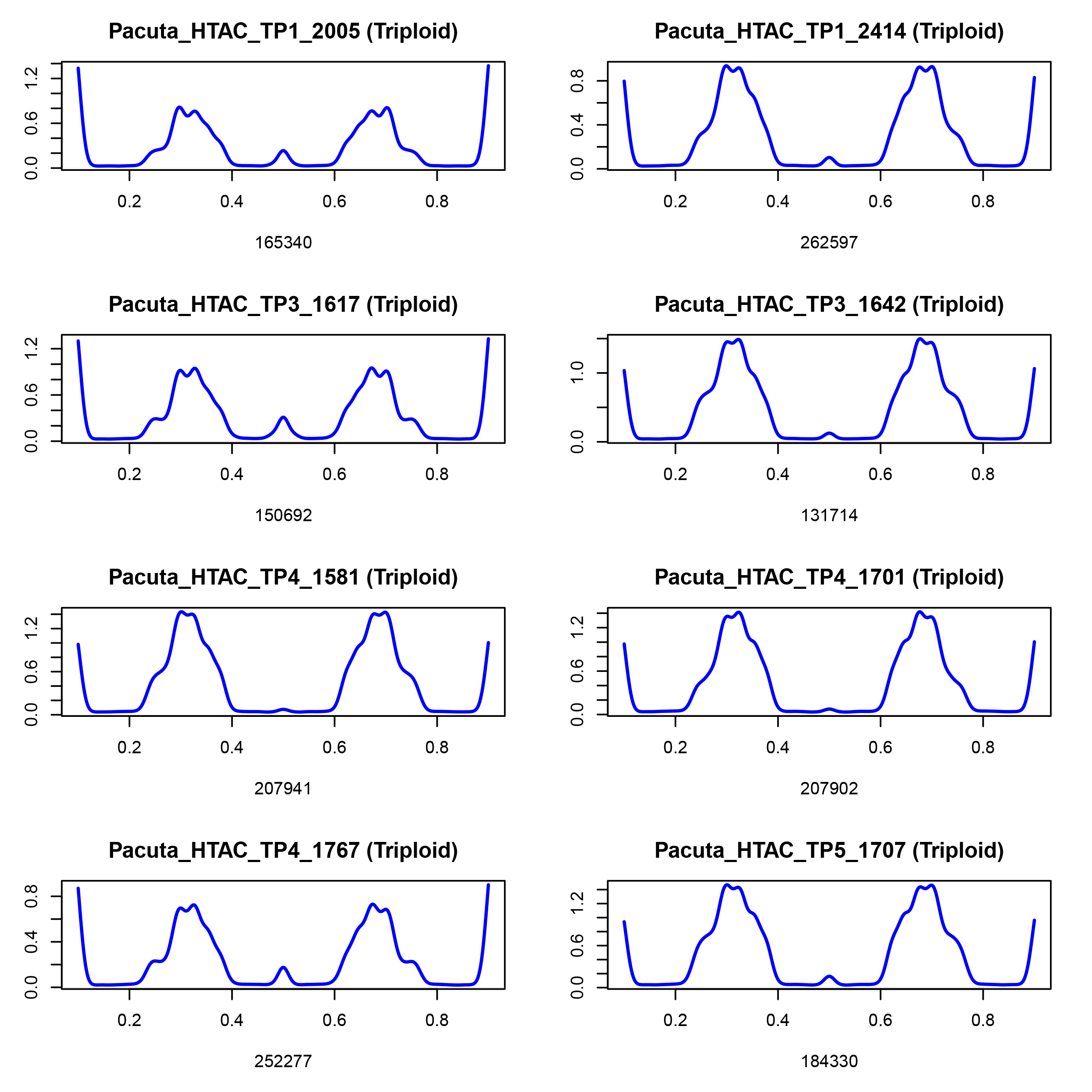
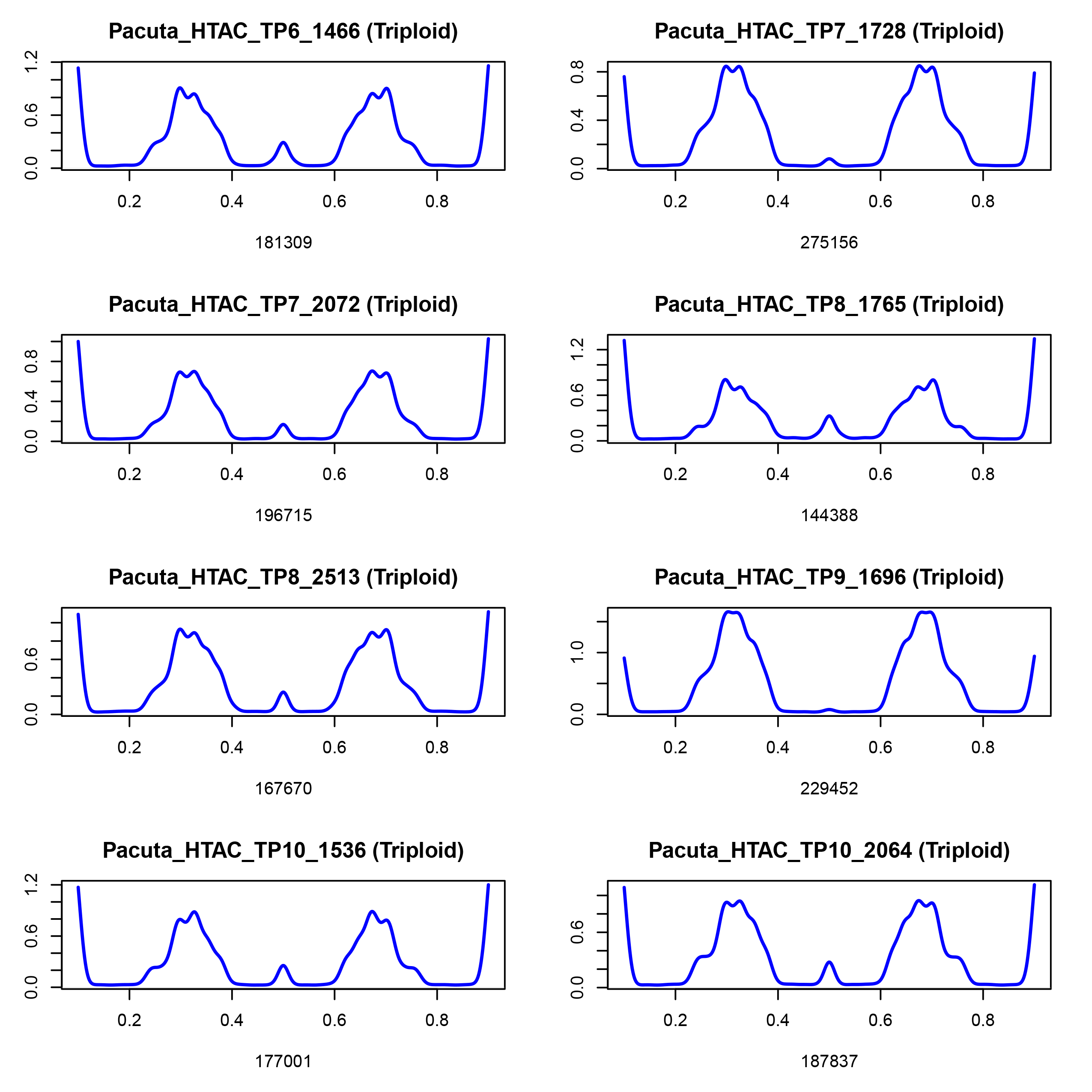
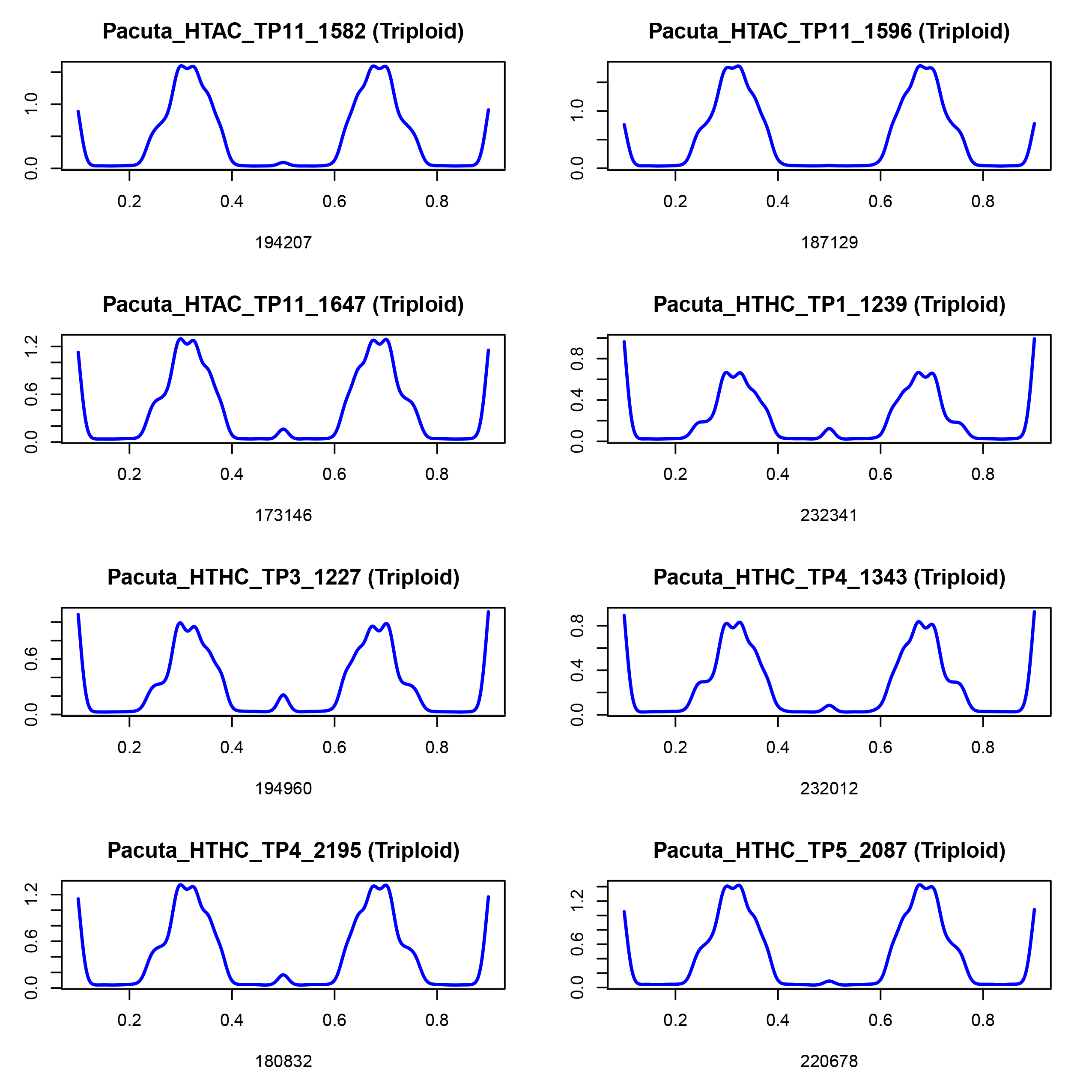
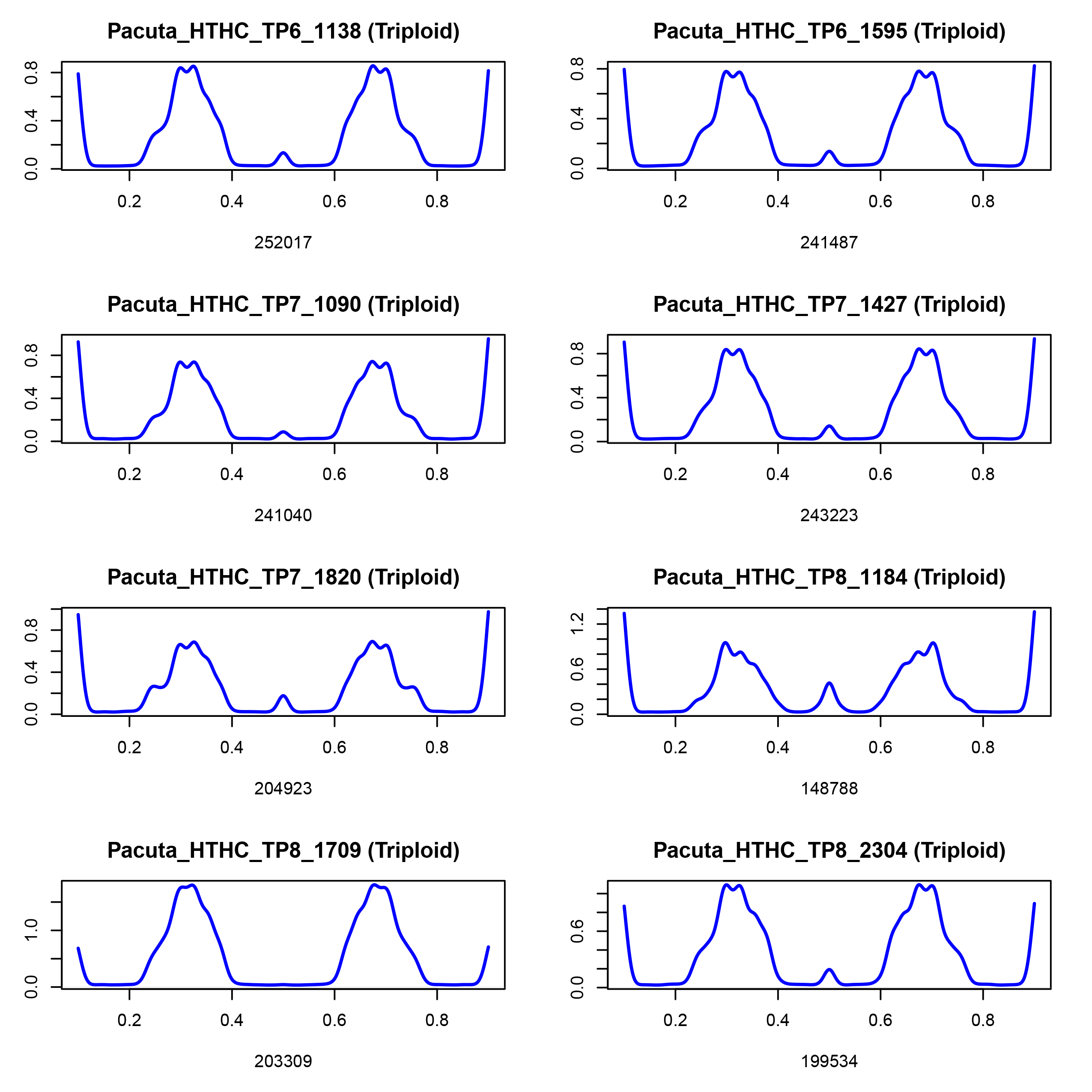
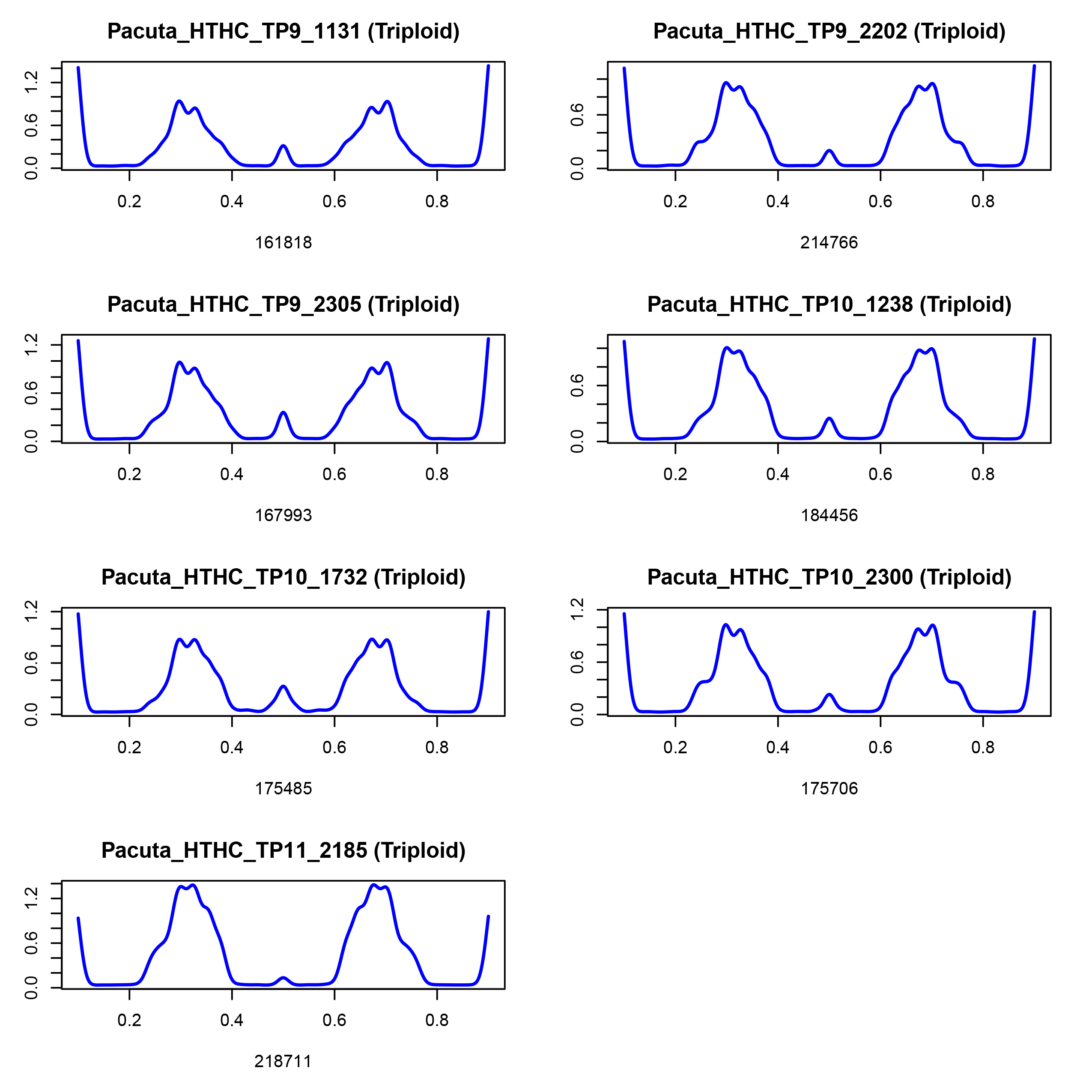
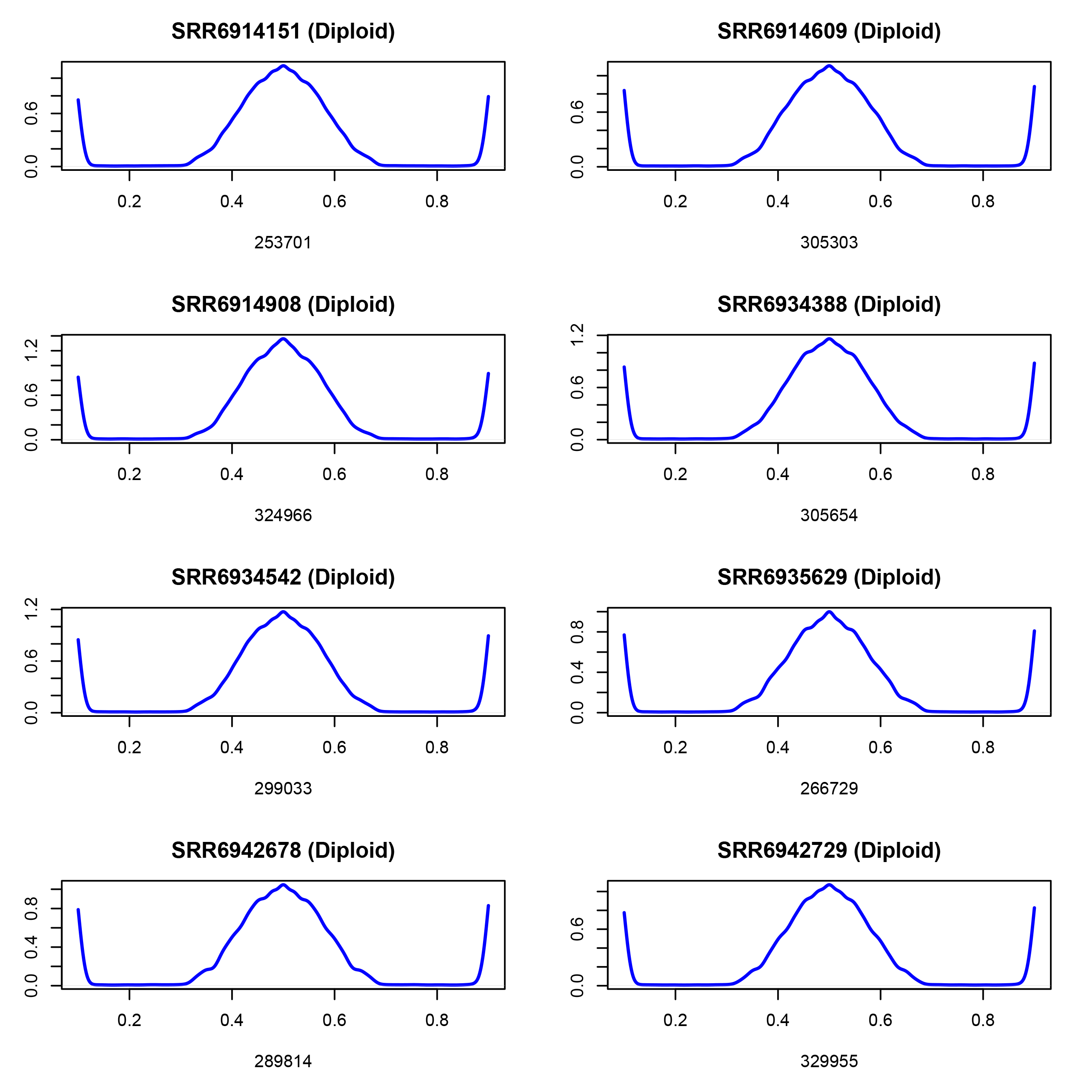
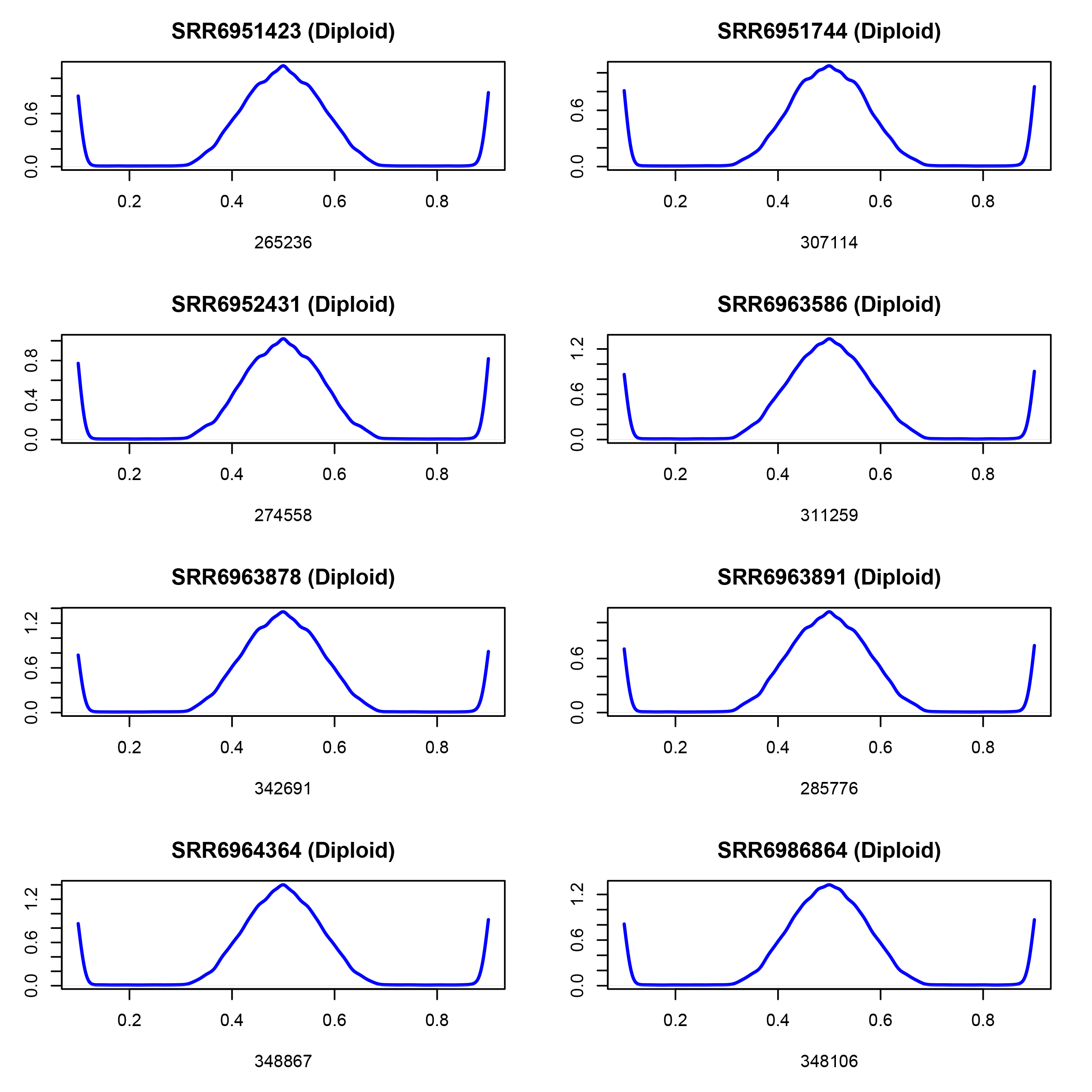
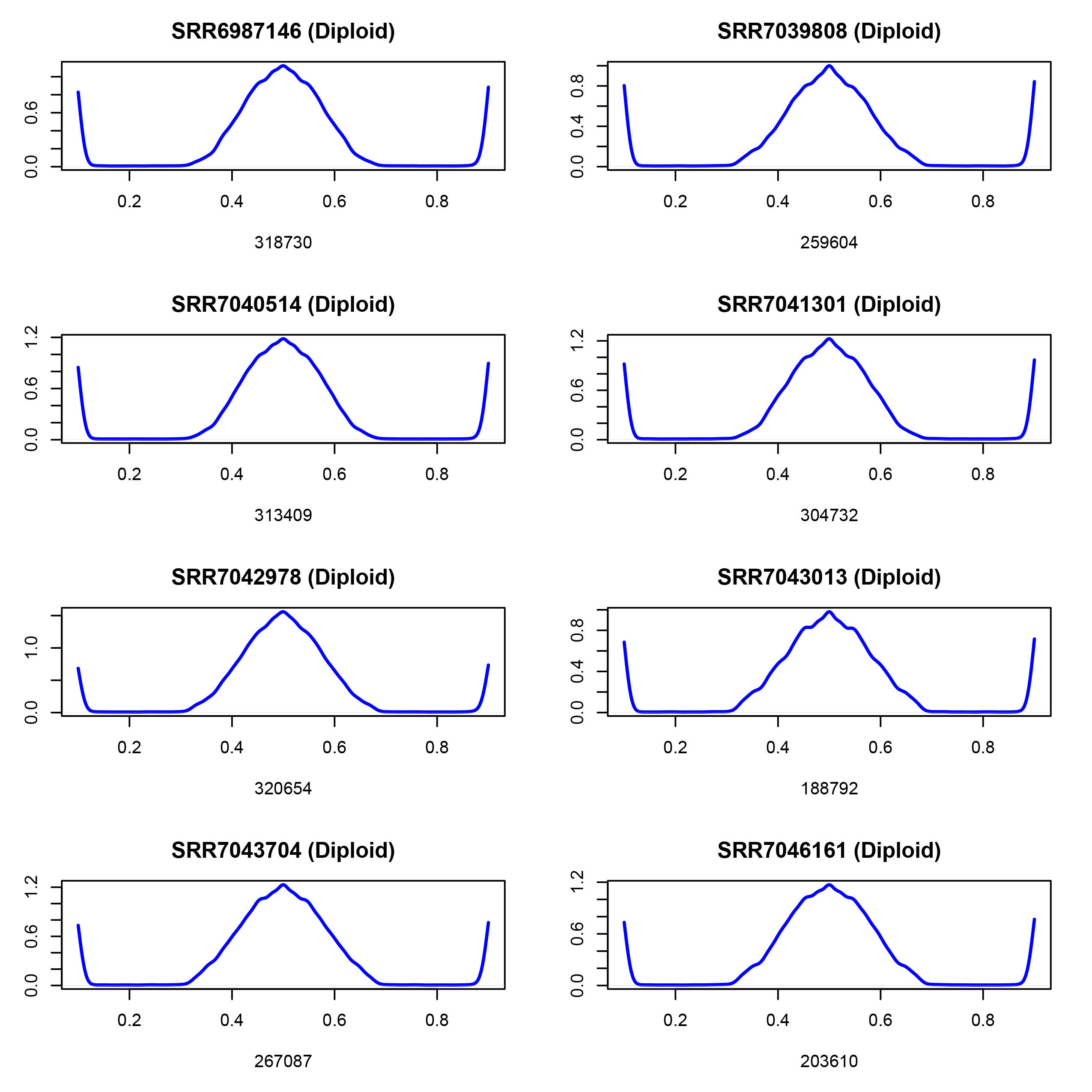
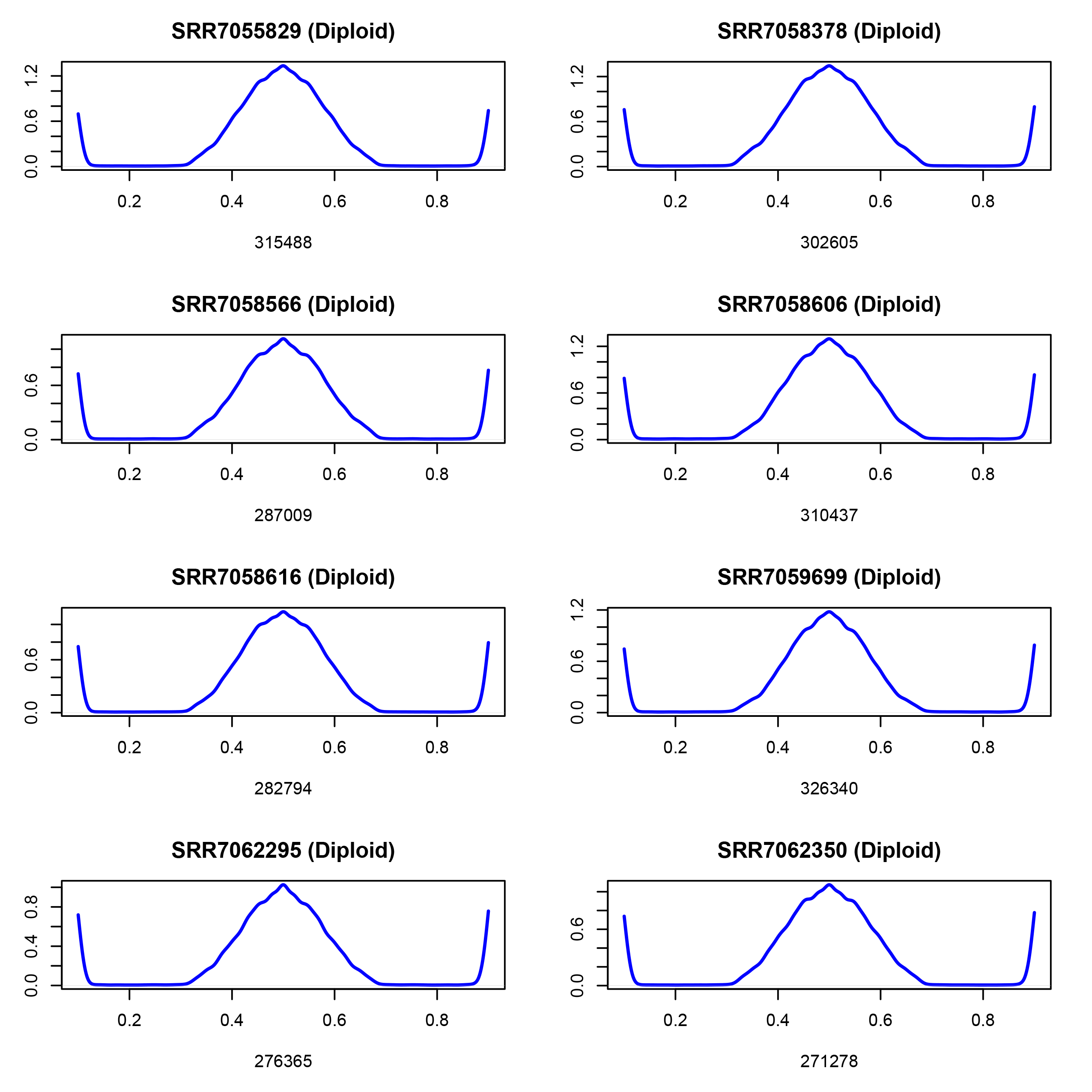
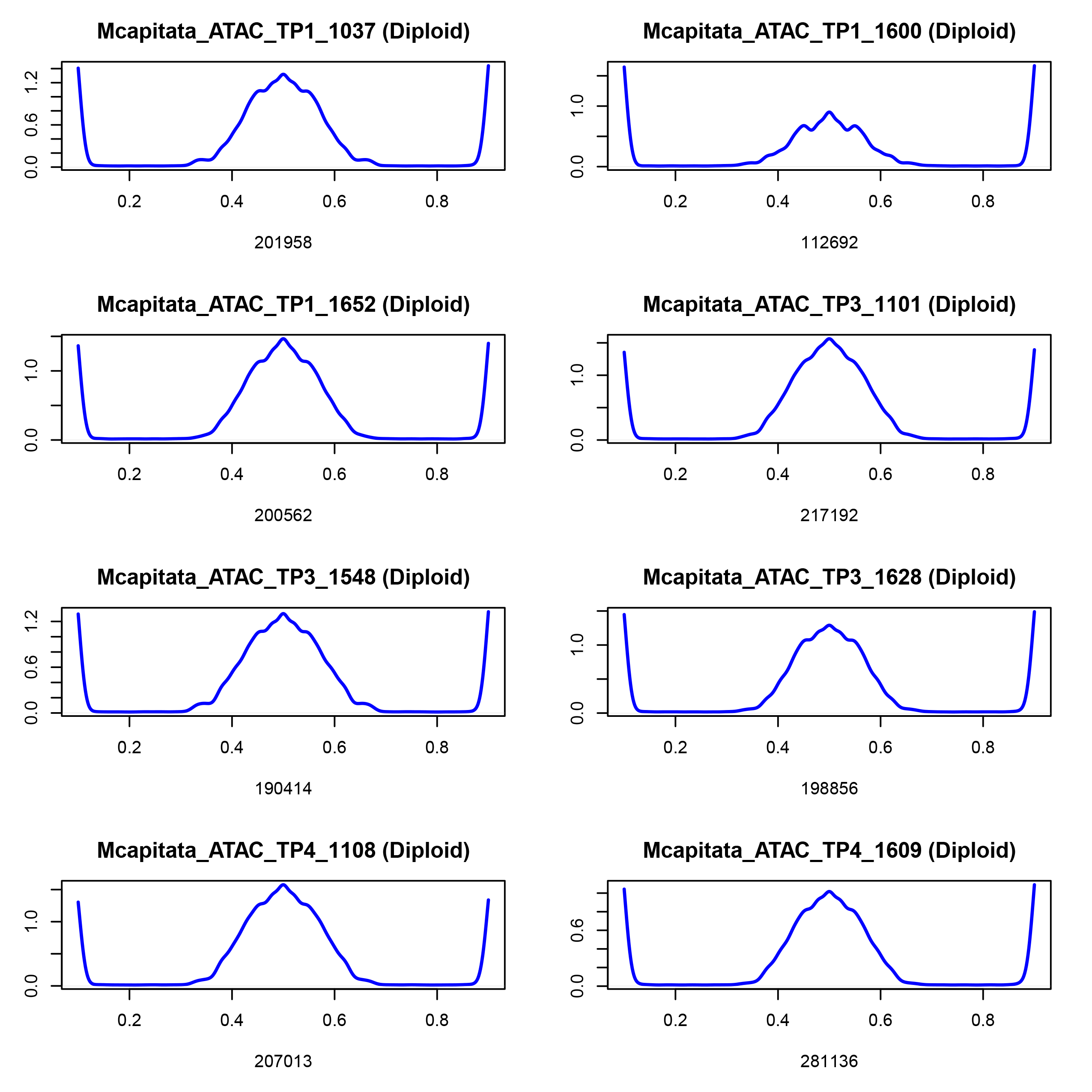
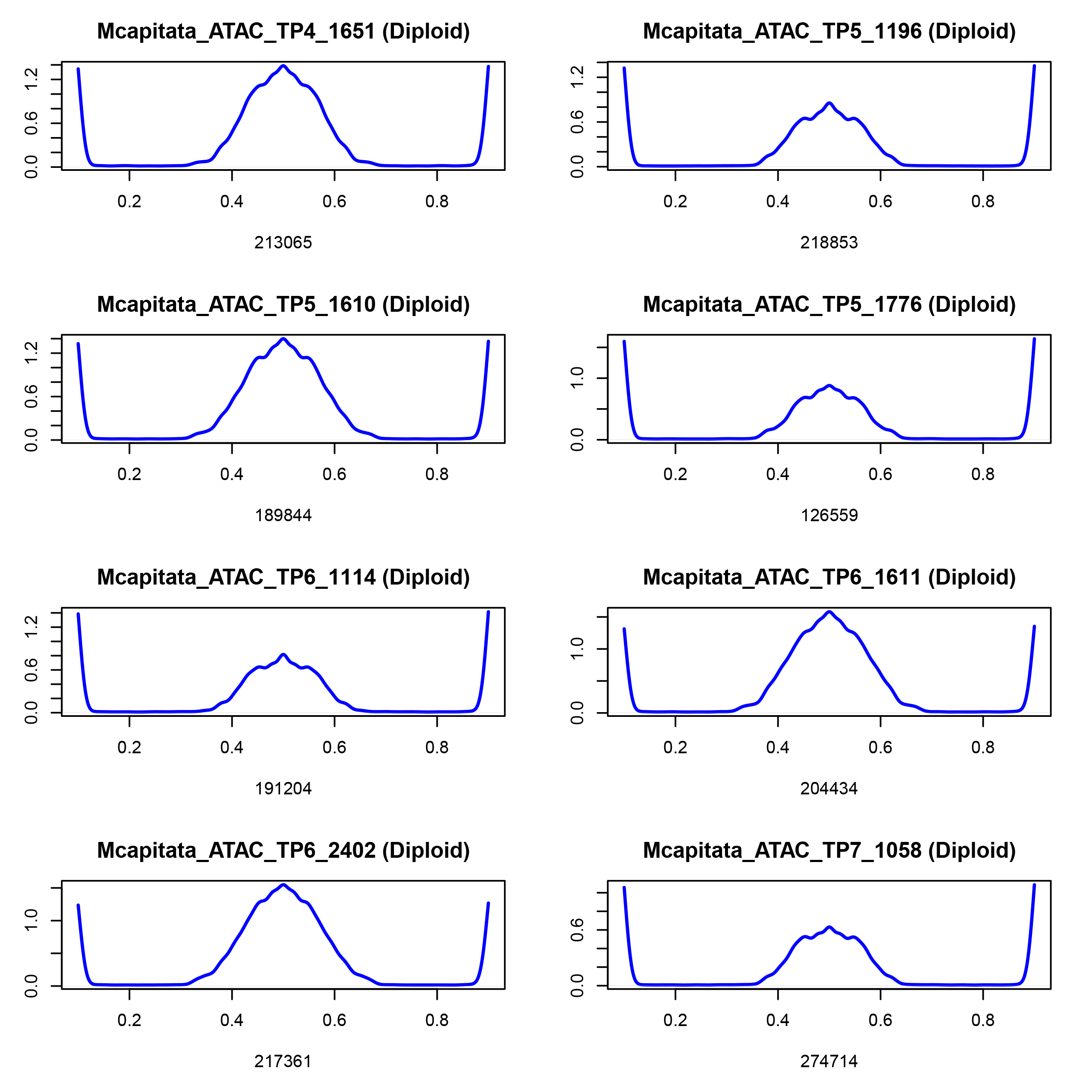
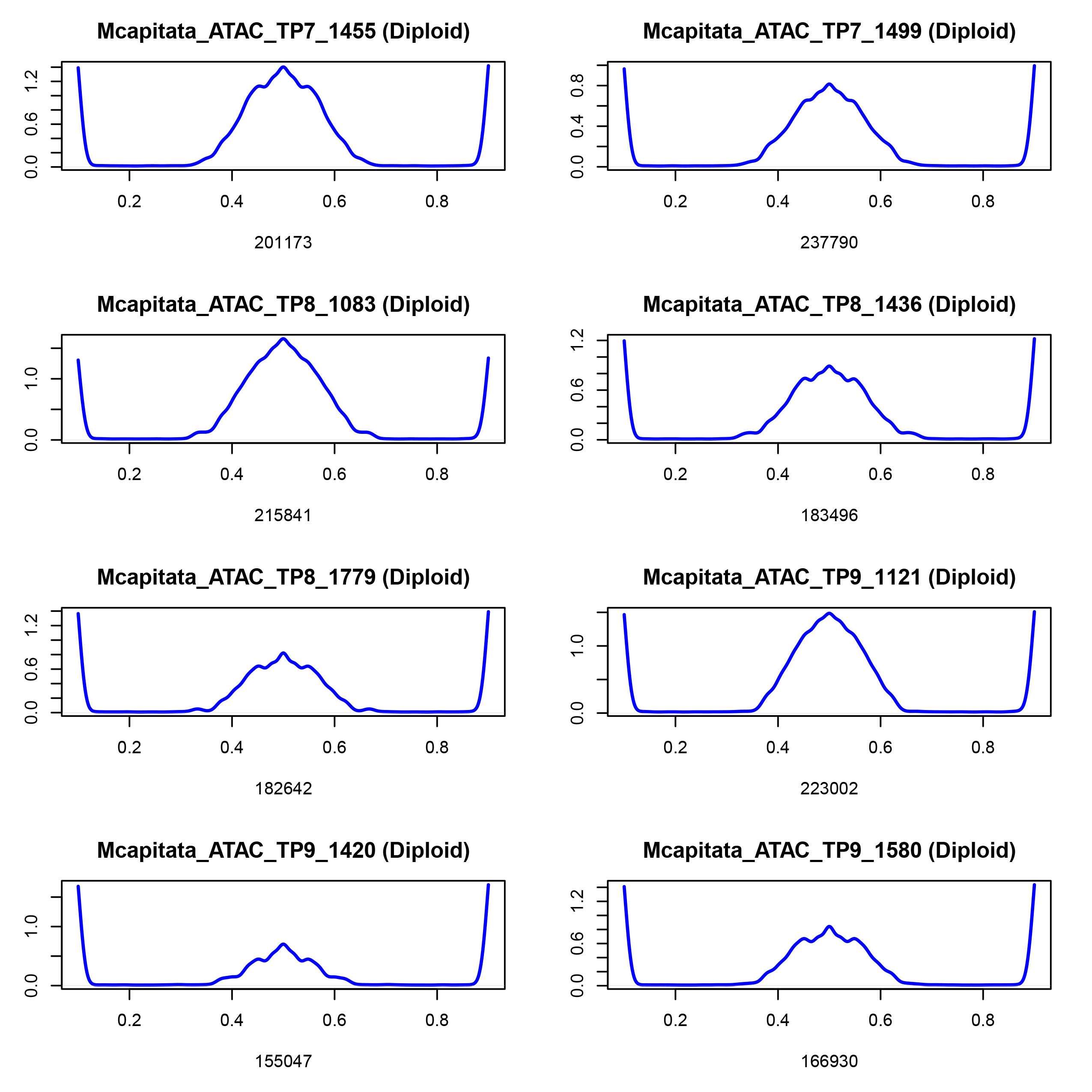
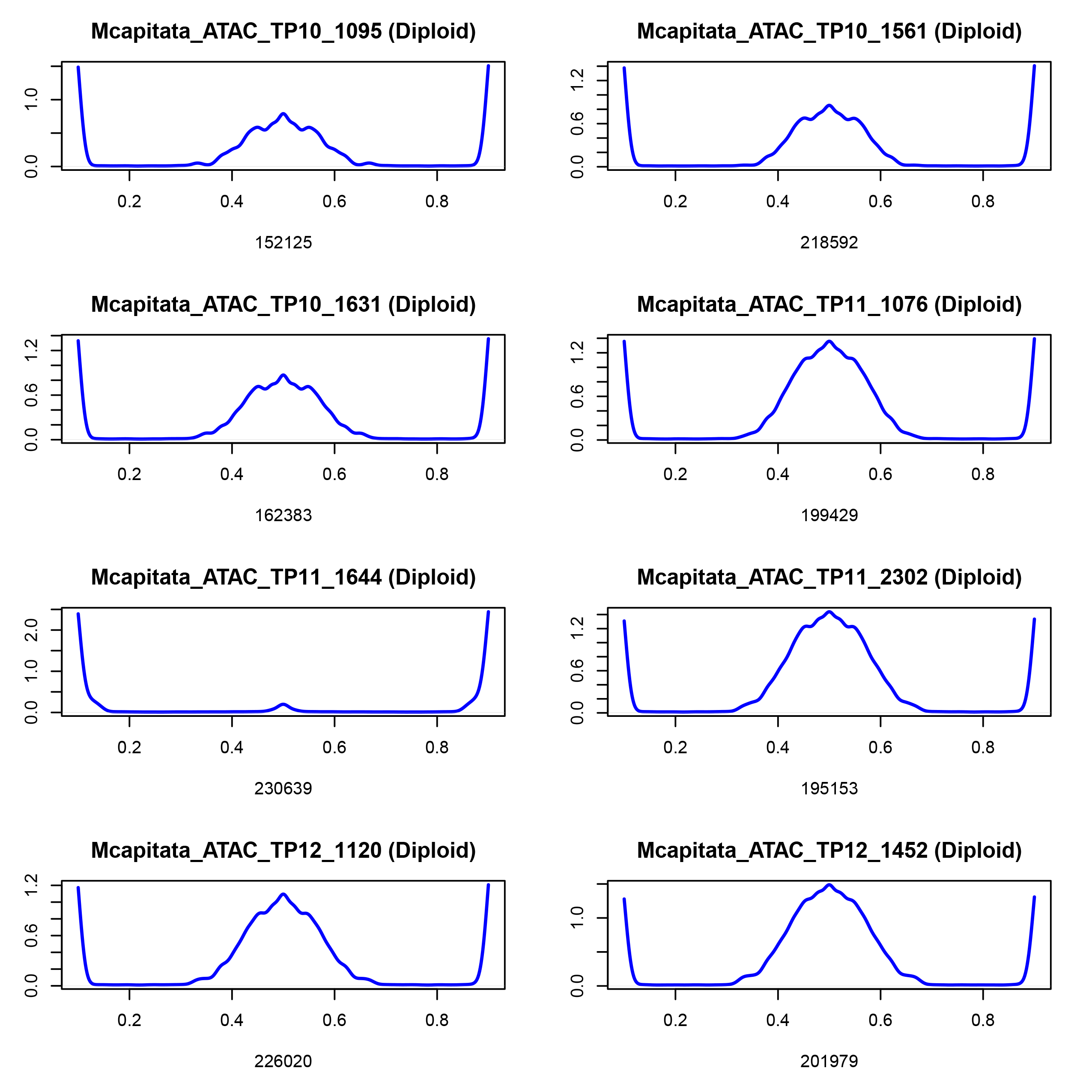
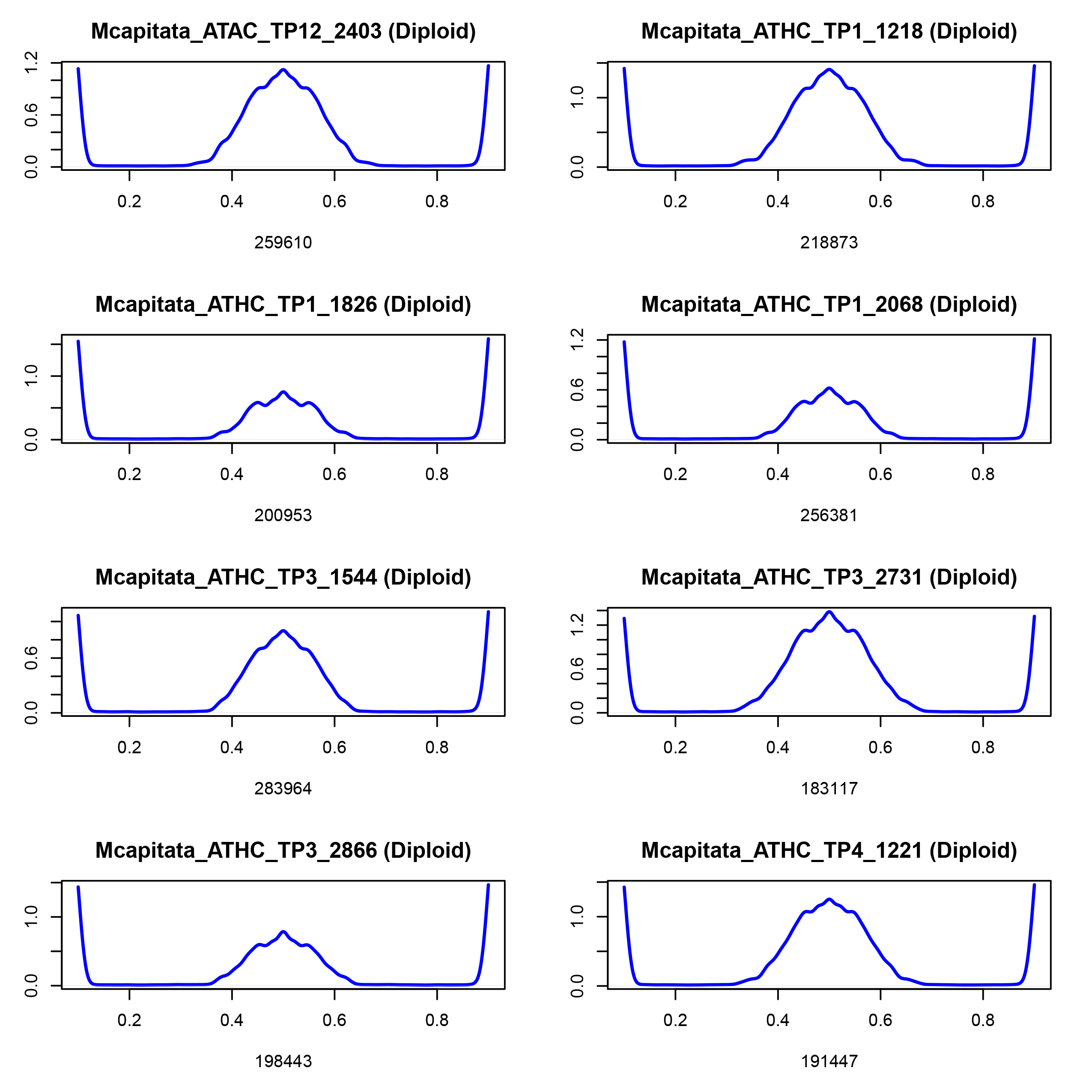
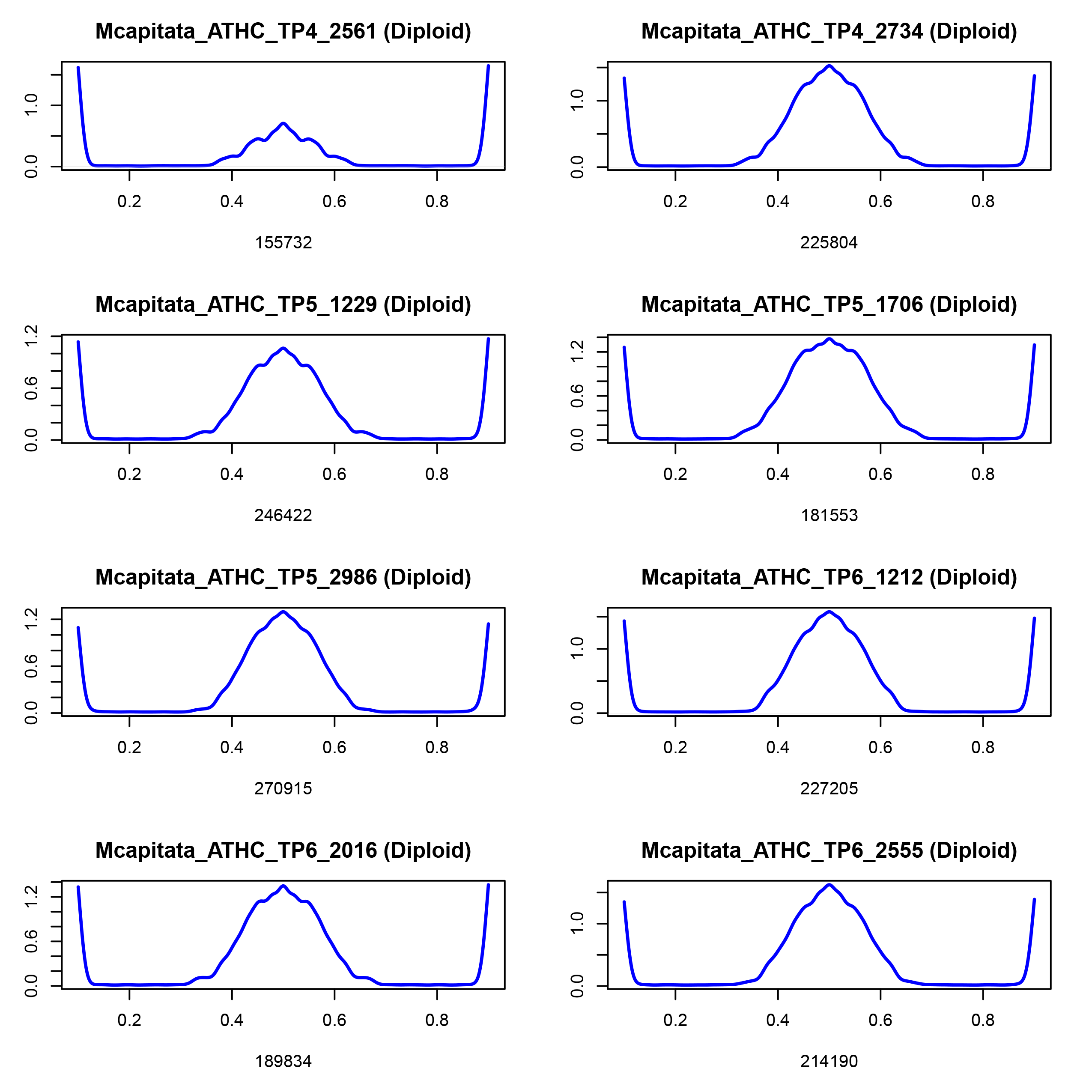
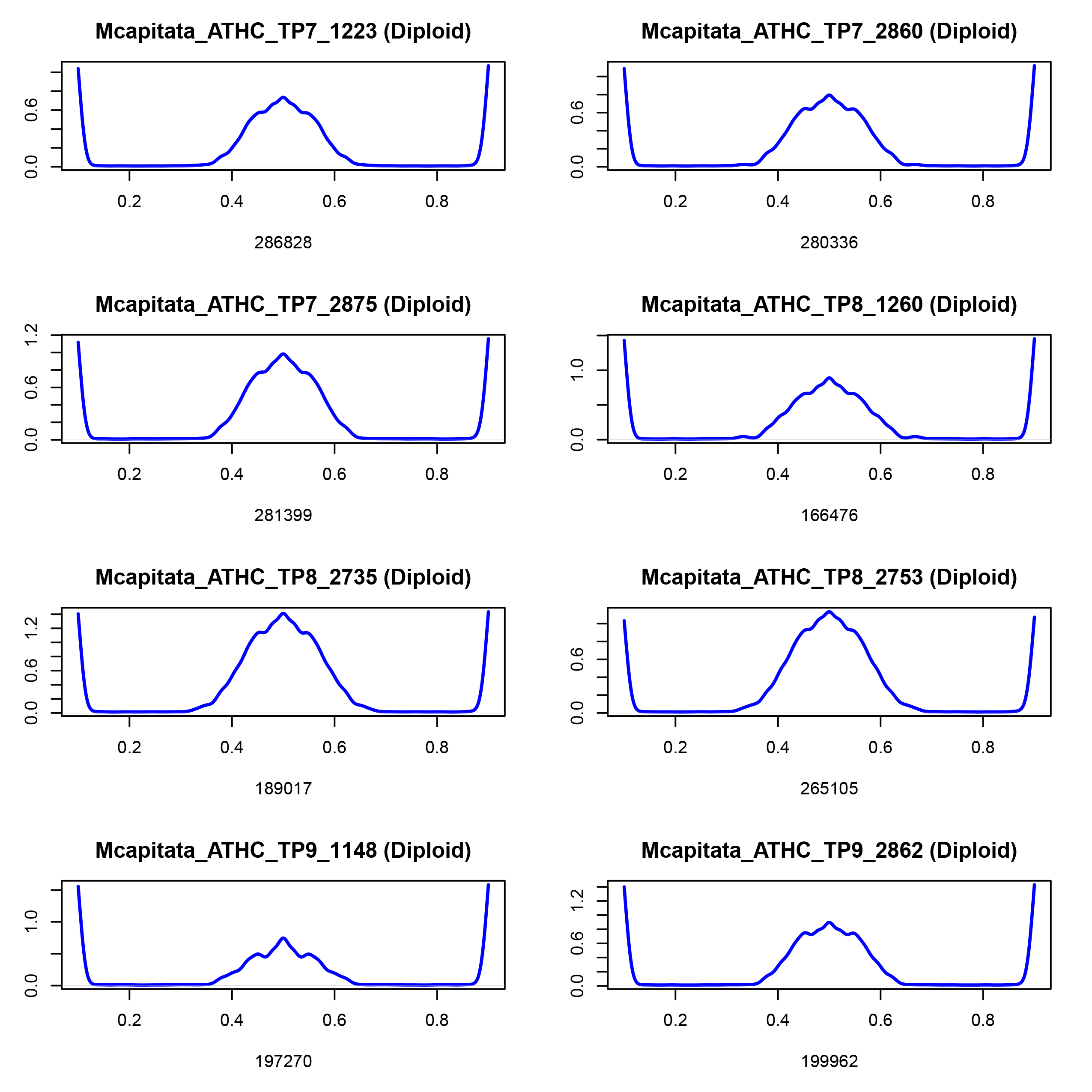
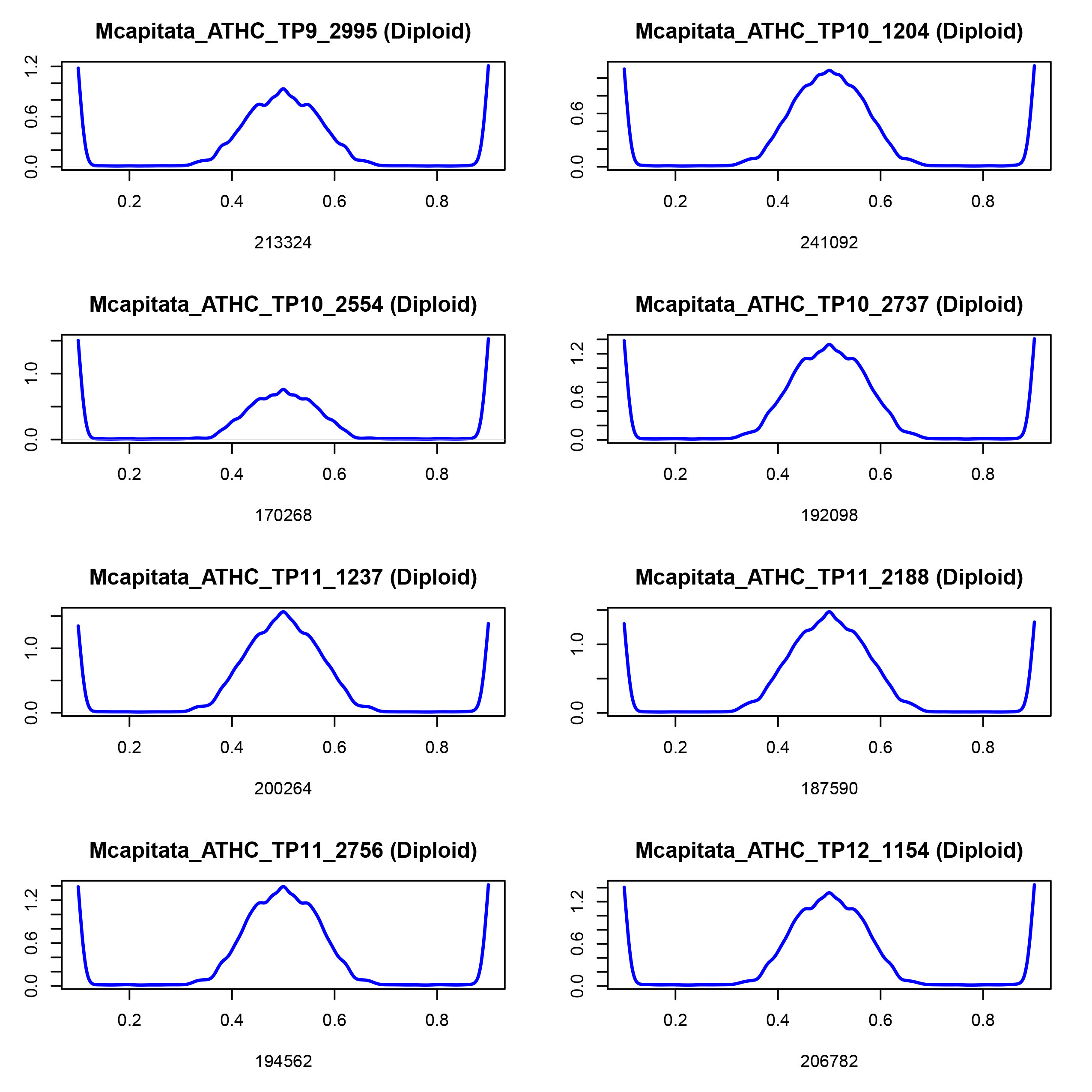
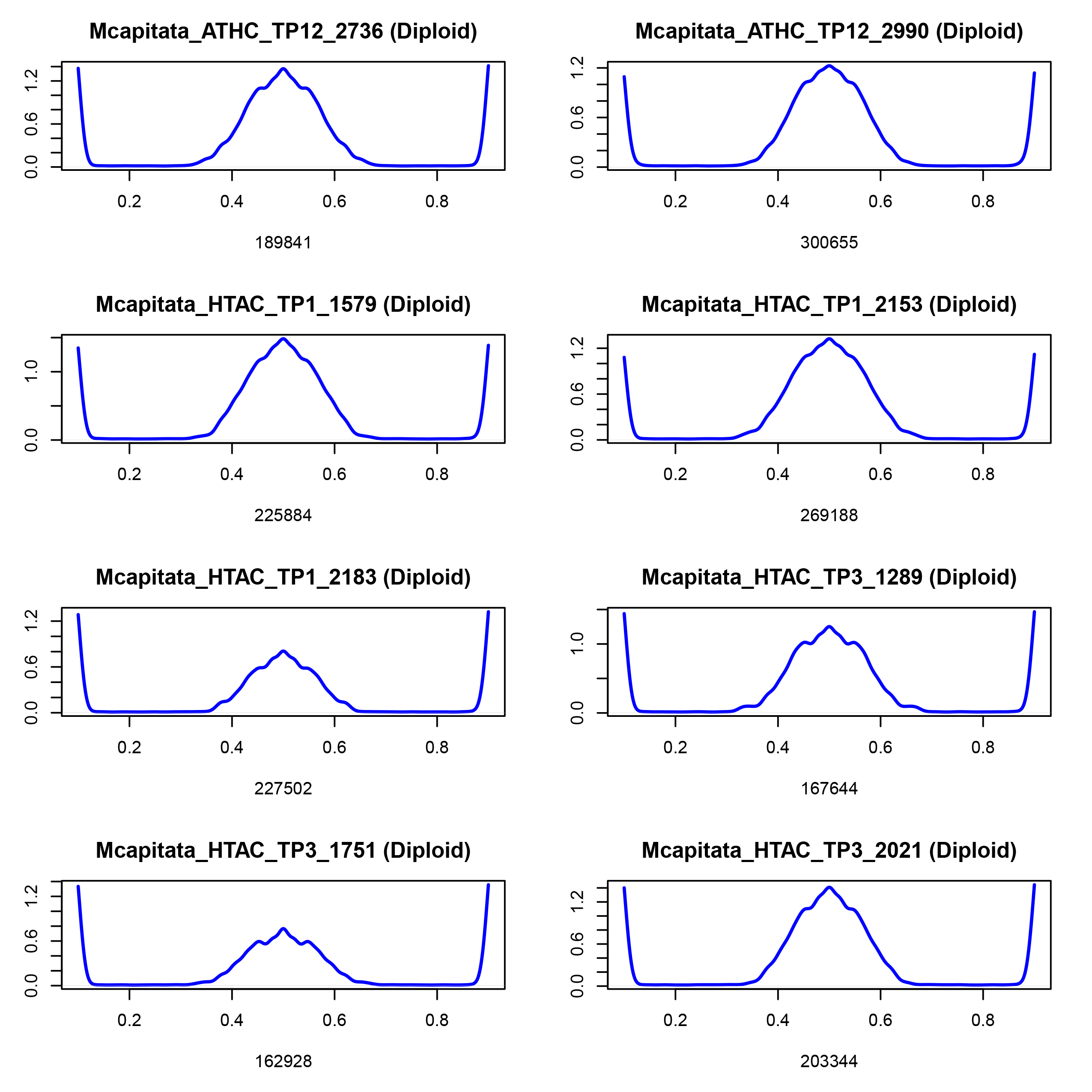
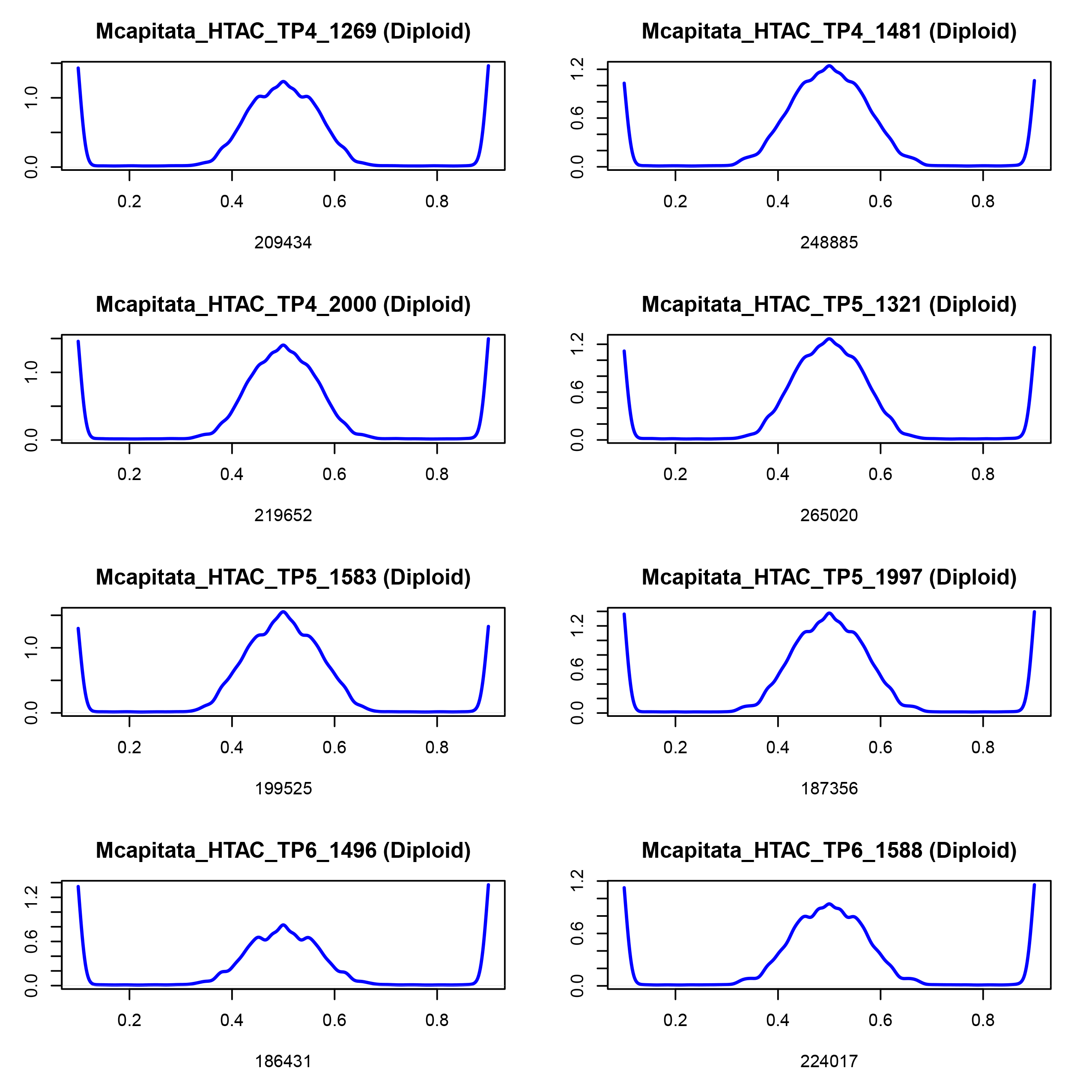
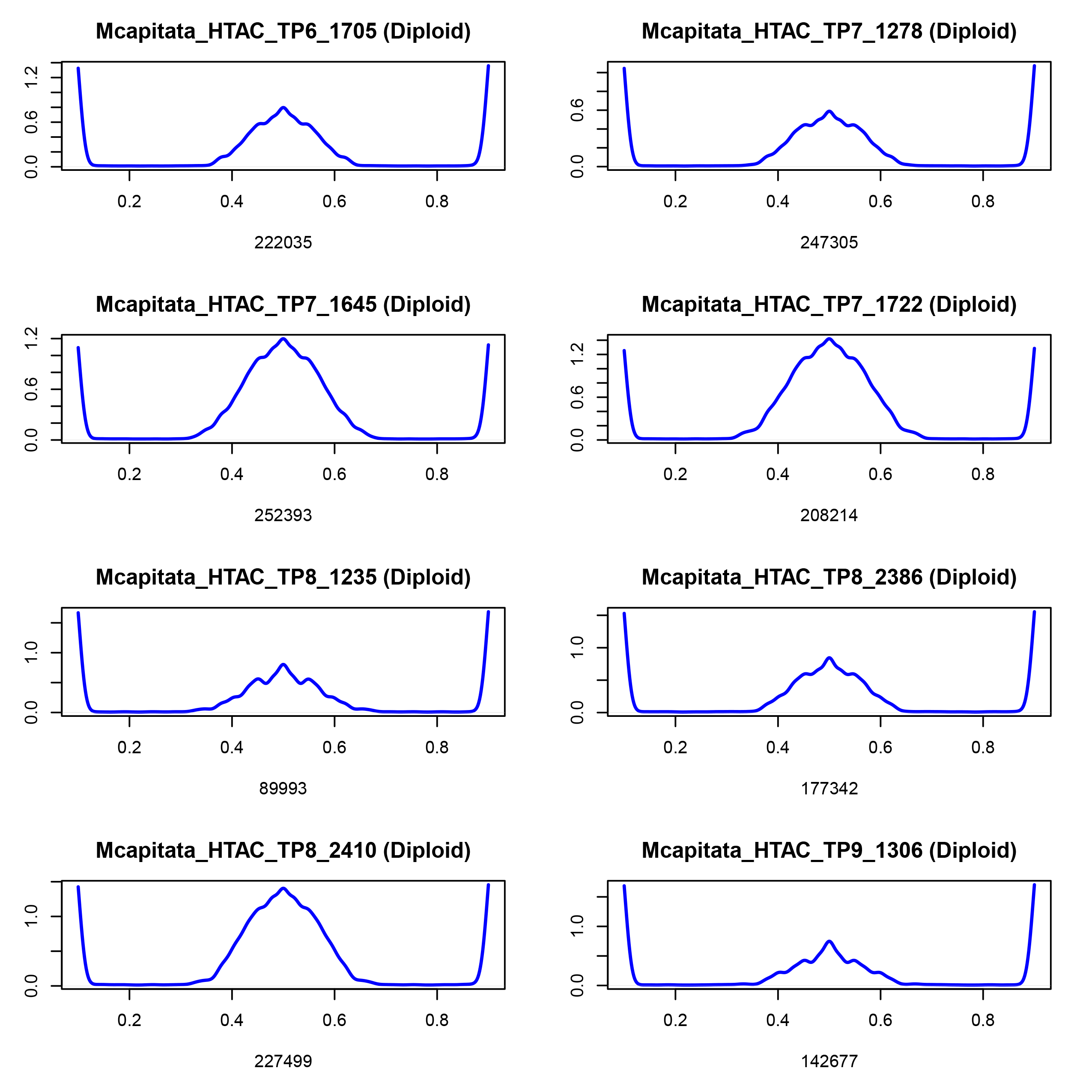
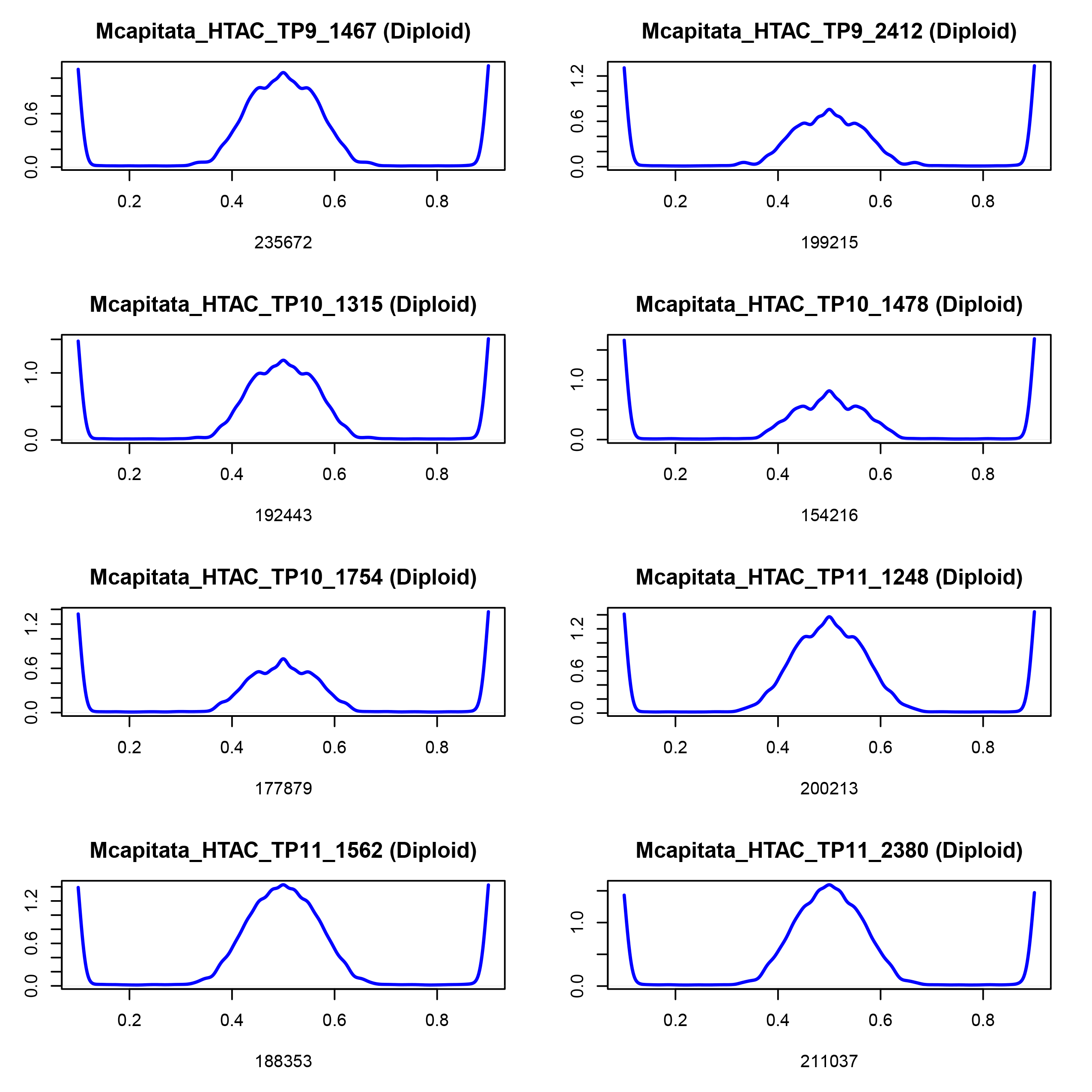
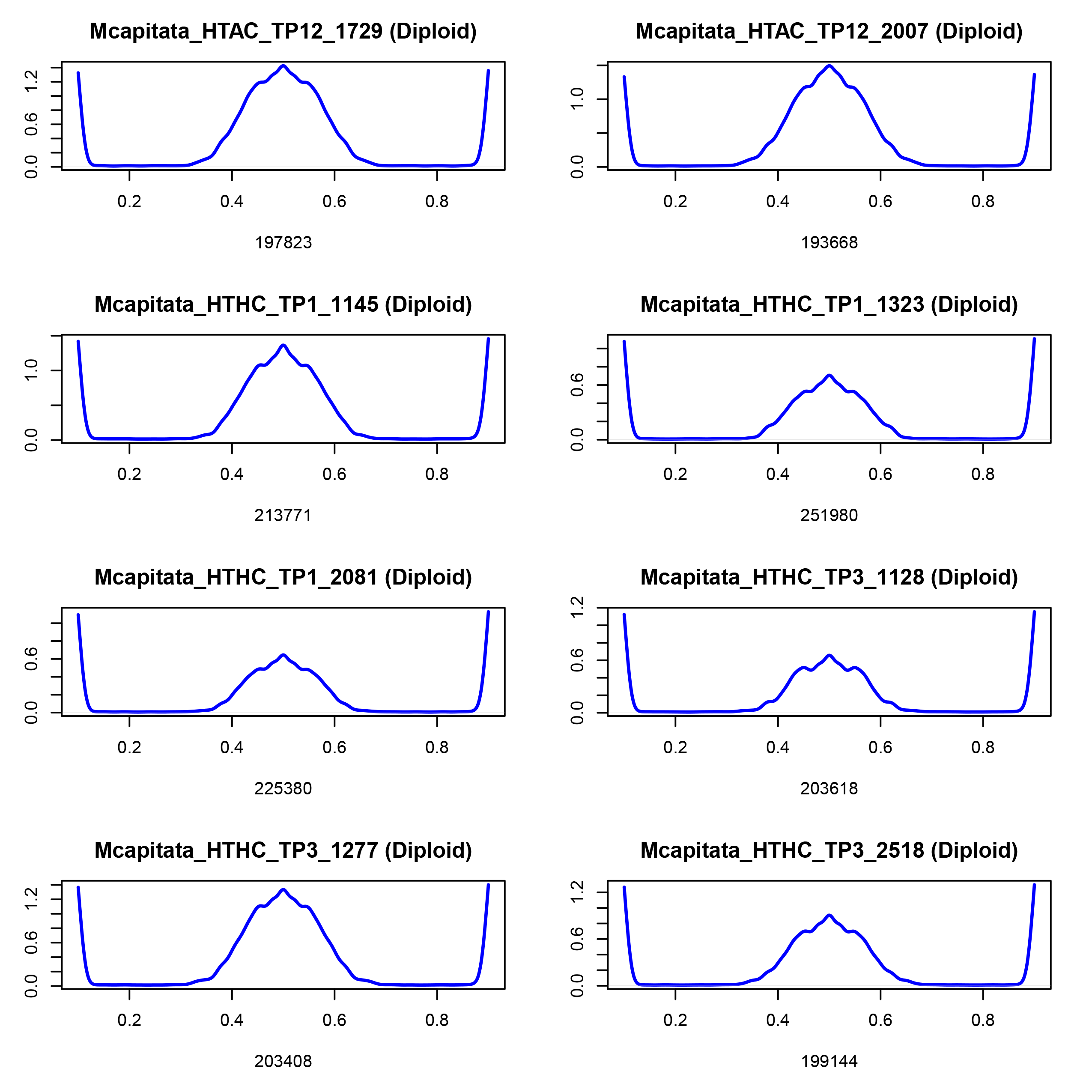
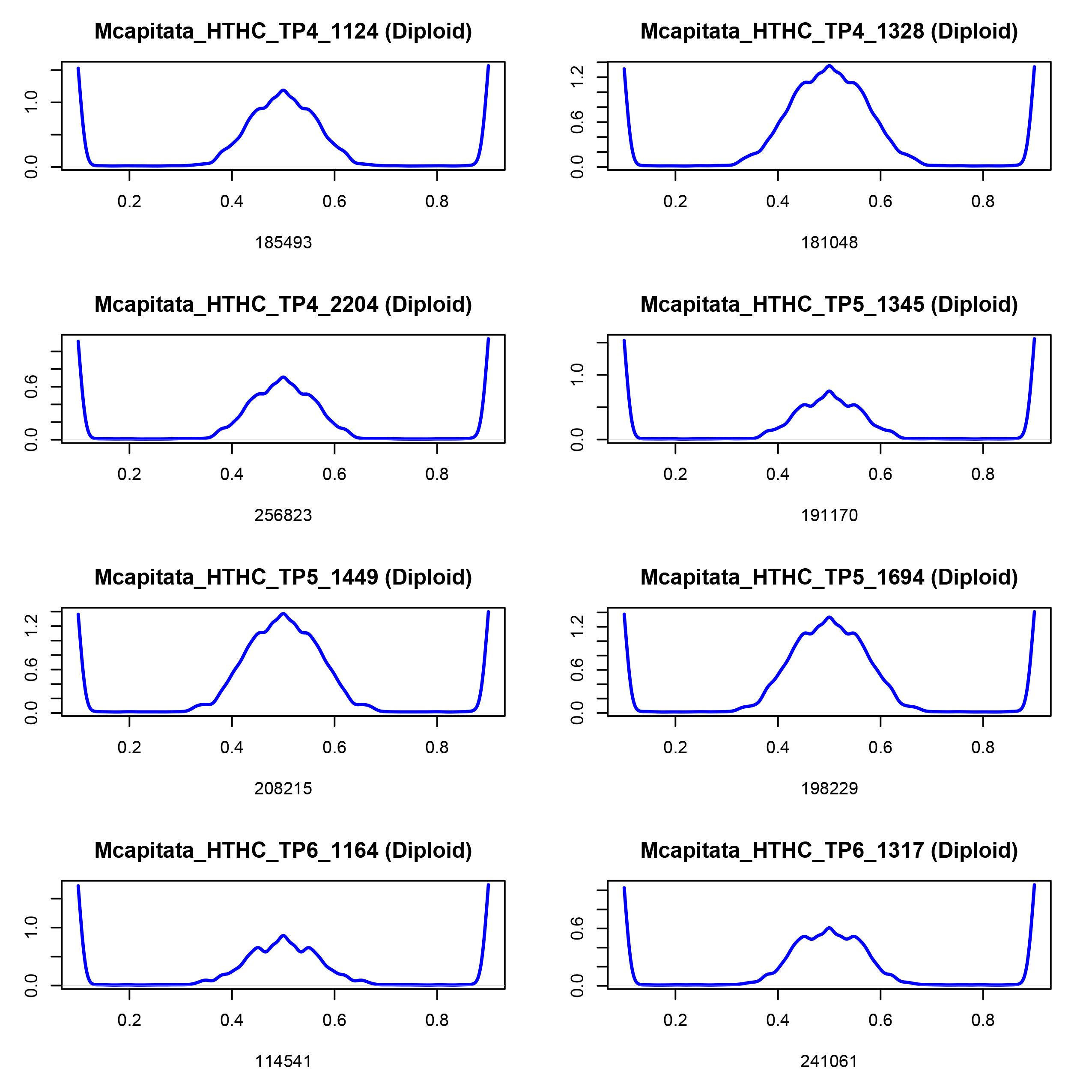
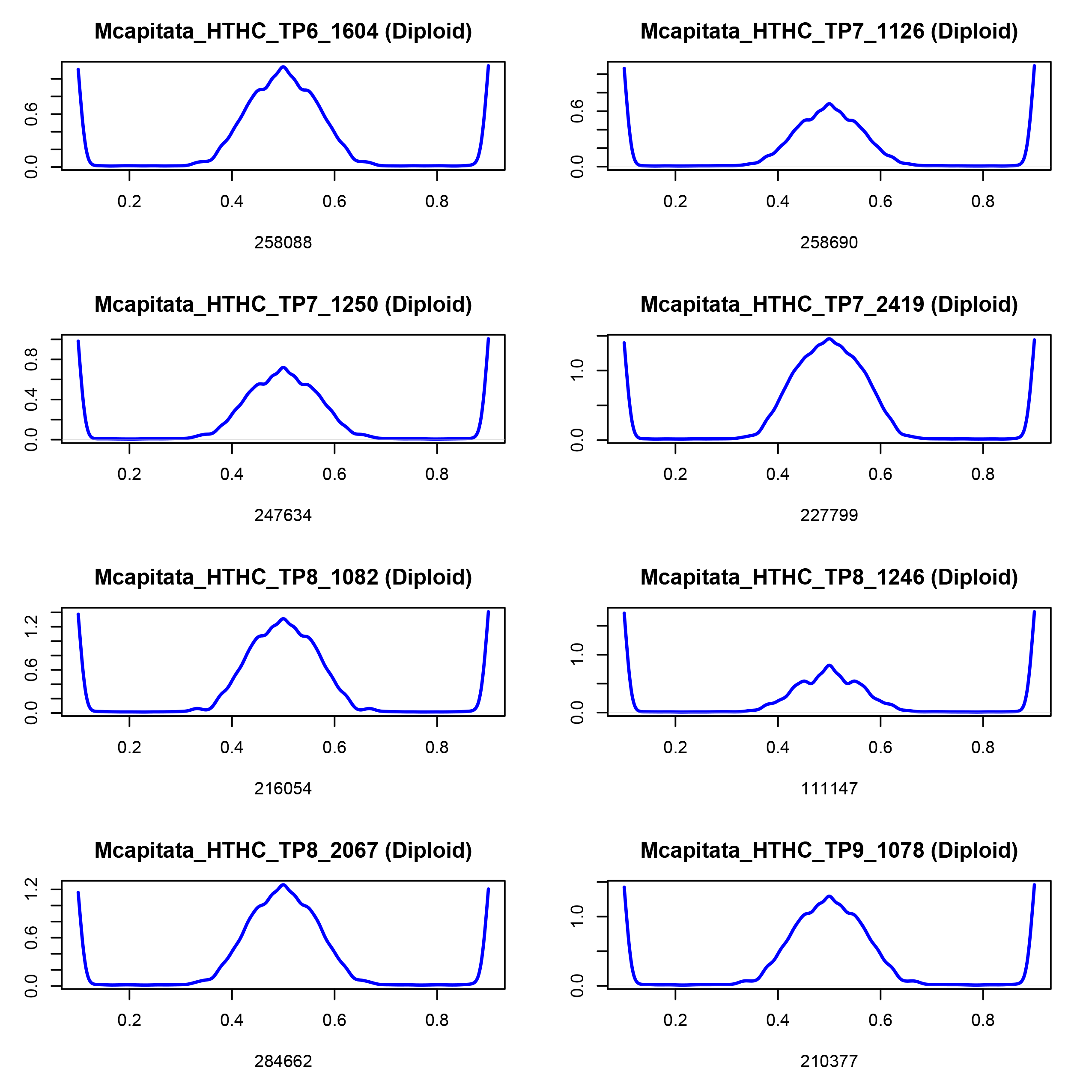
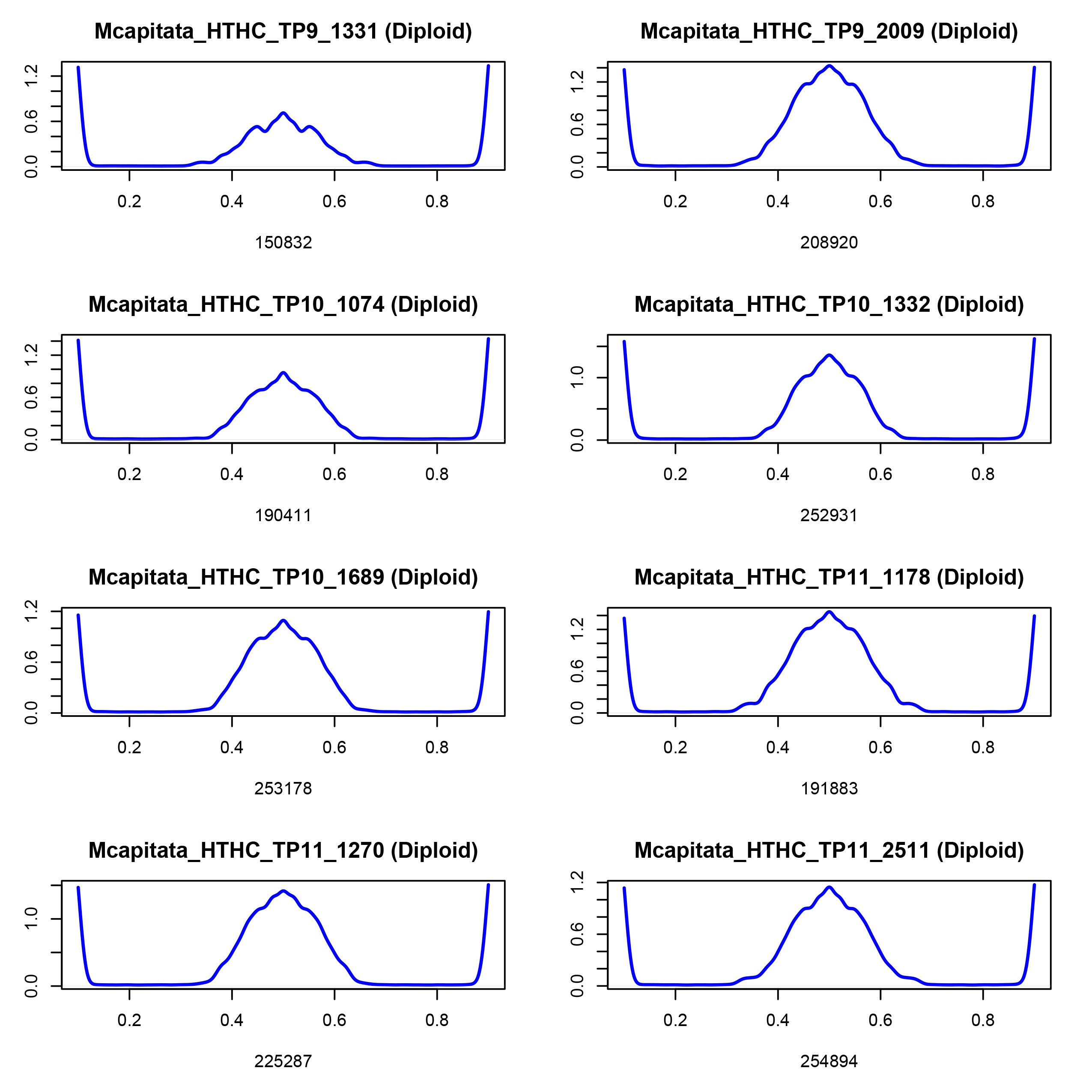
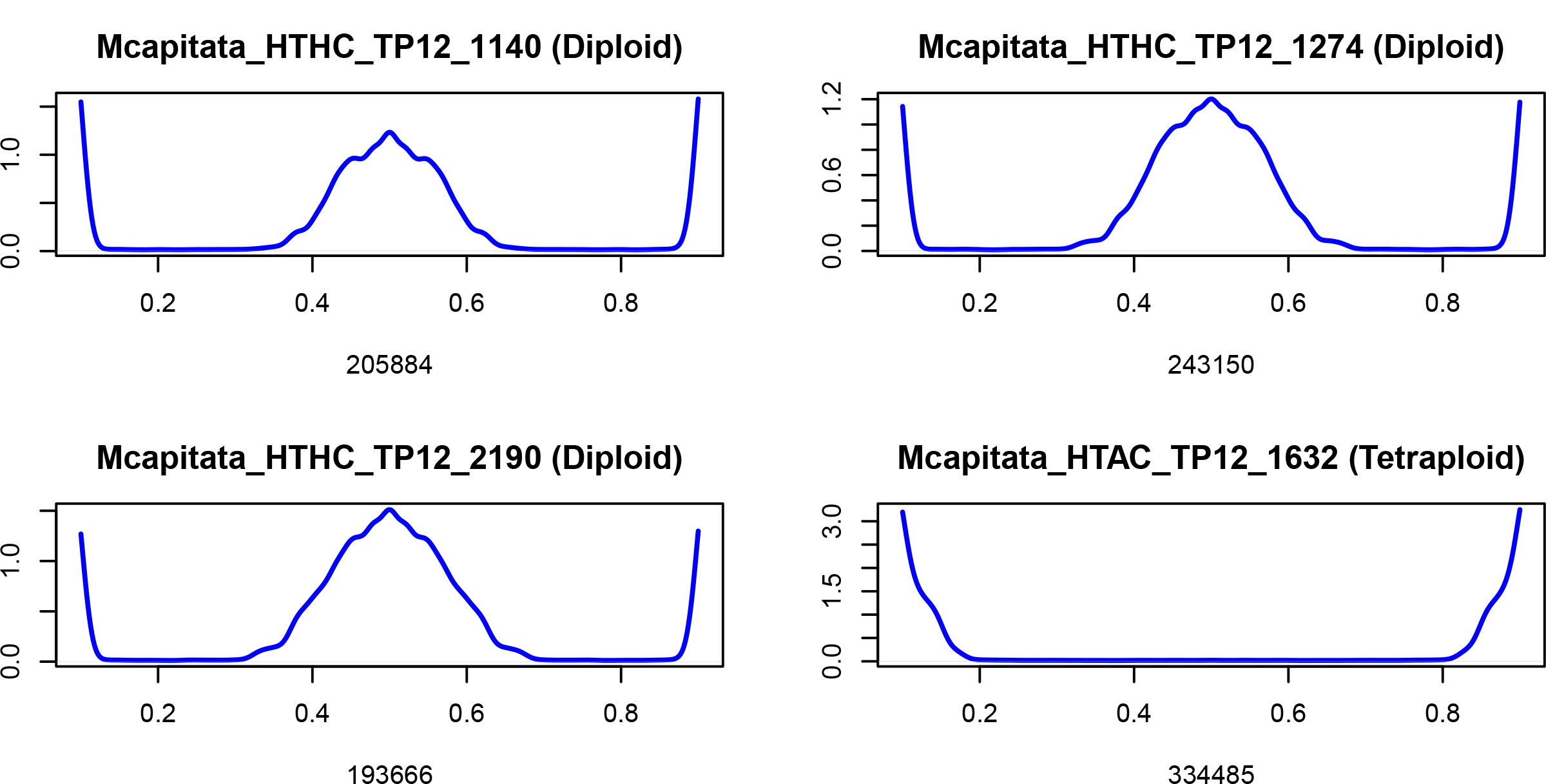
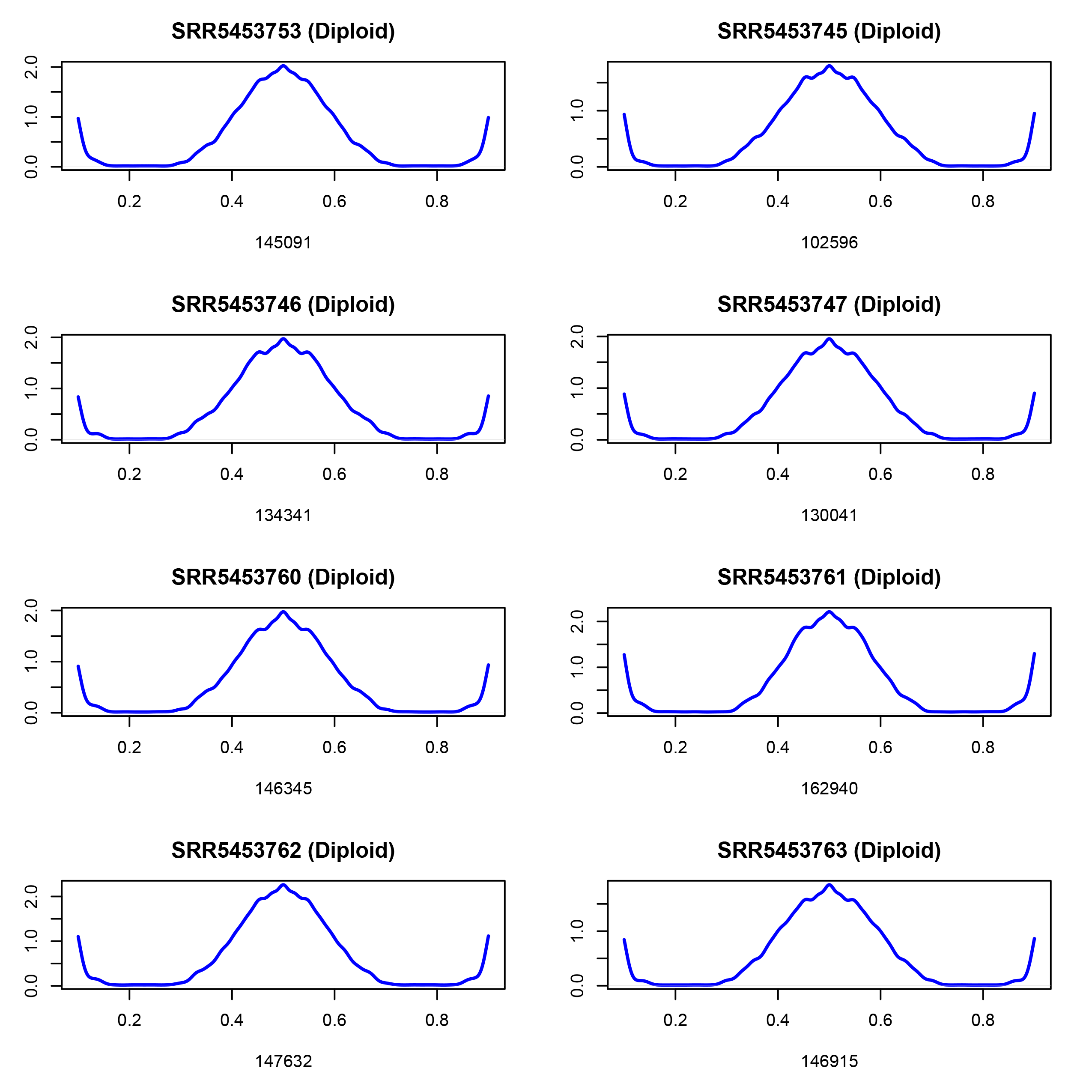
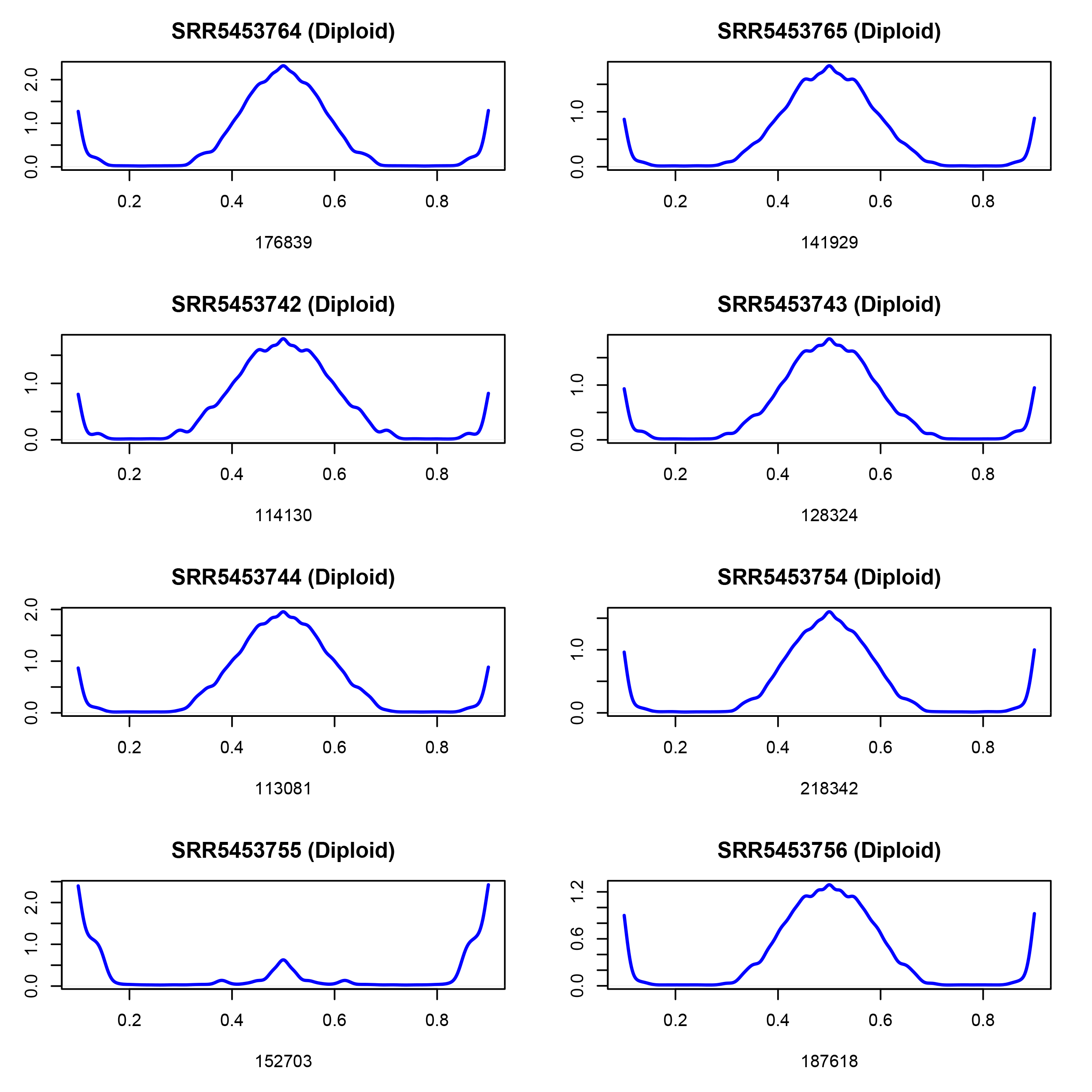
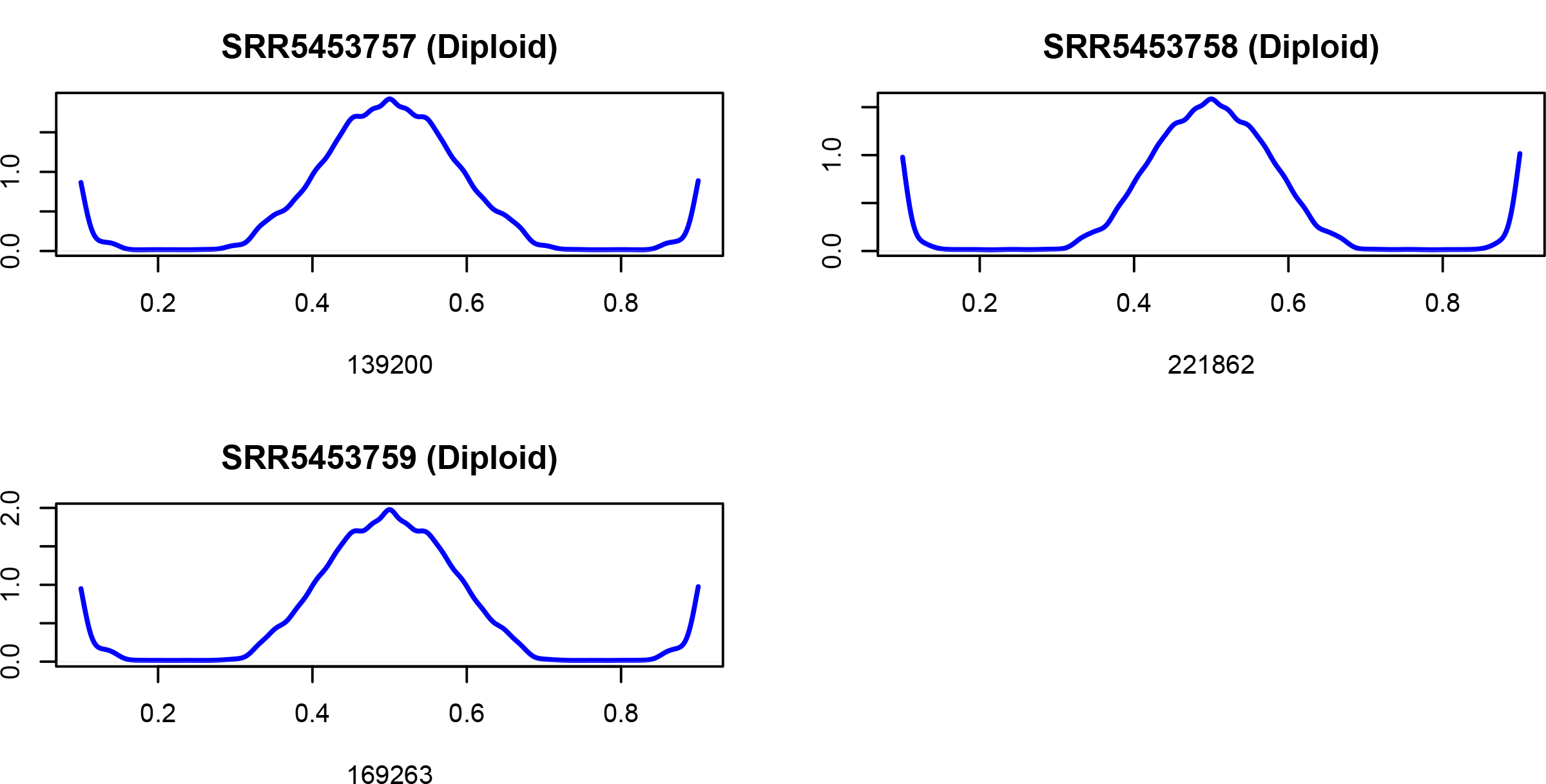

## Notes

### Competing Interest Statement

The authors have declared no competing interest.

### Summary of Updates

Sections of the manuscript have been removed because they were not part of the core story or they were published as standalone papers. Additional analysis has also been conducted based on comments made by Reviewers during several rounds of submission to scientific journals.

