## Supplementary figures and images for "Ploidy variation and its implications for reproduction and population dynamics in two sympatric Hawaiian coral species"

### Figure_1.jpg

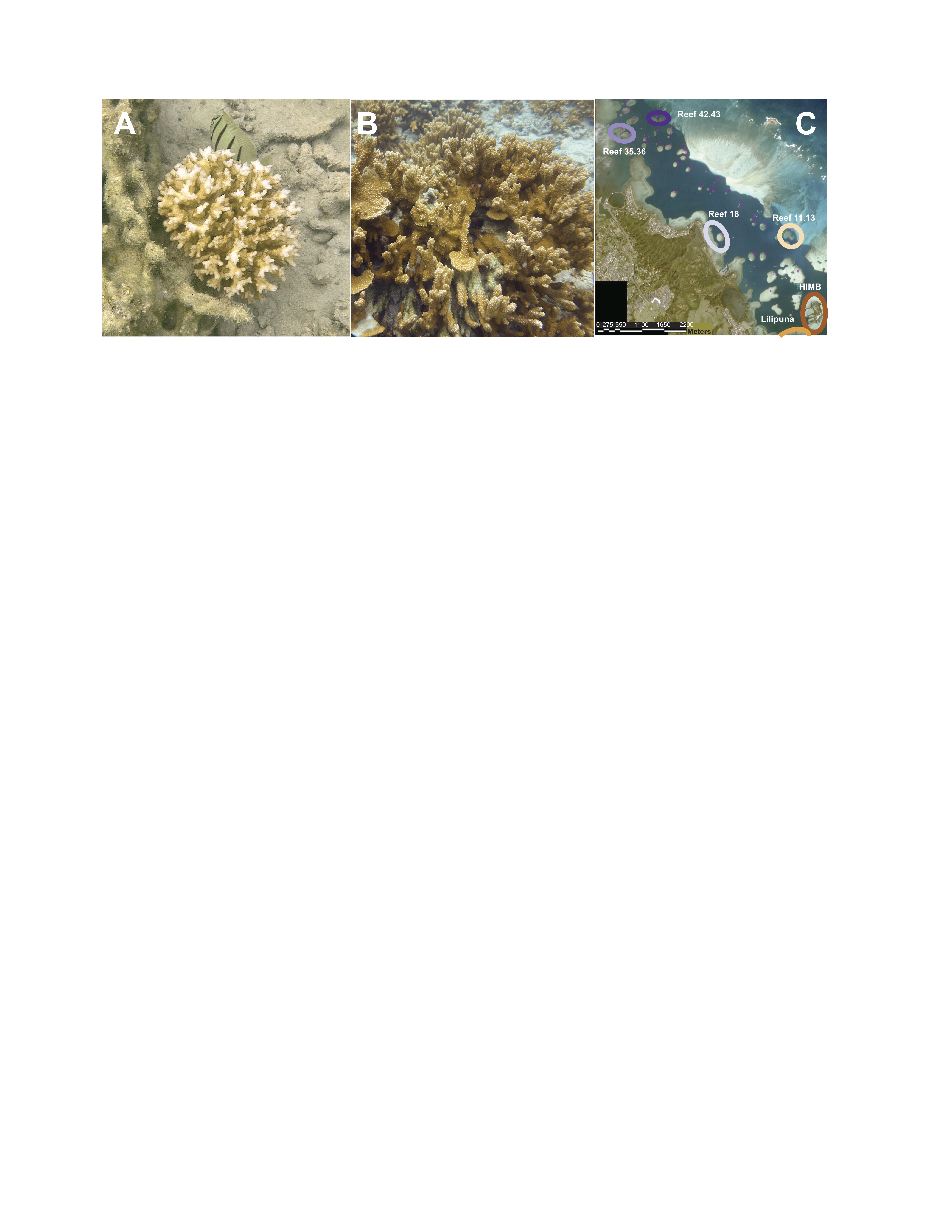

### Figure_2.pdf

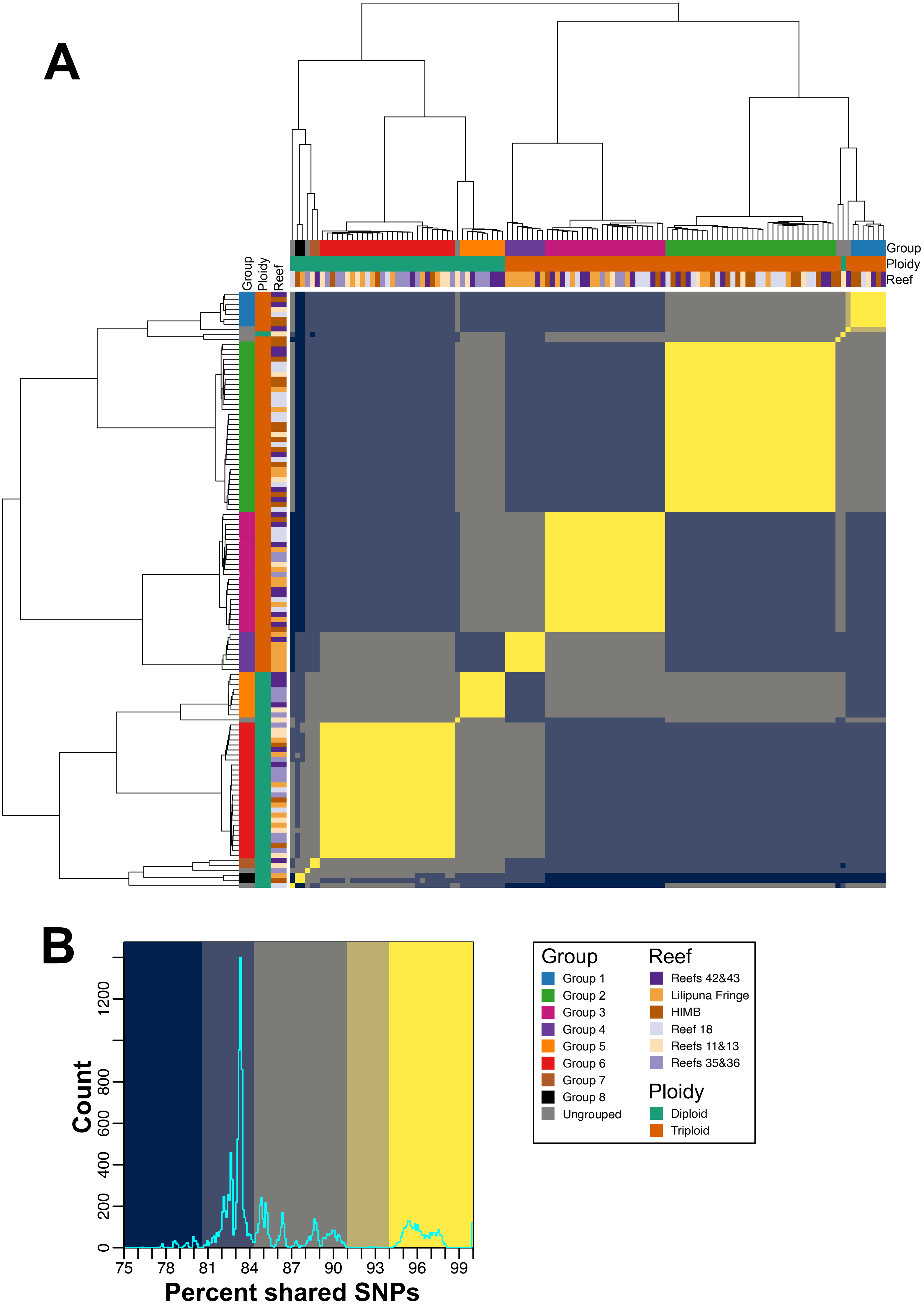

### Figure_3.pdf

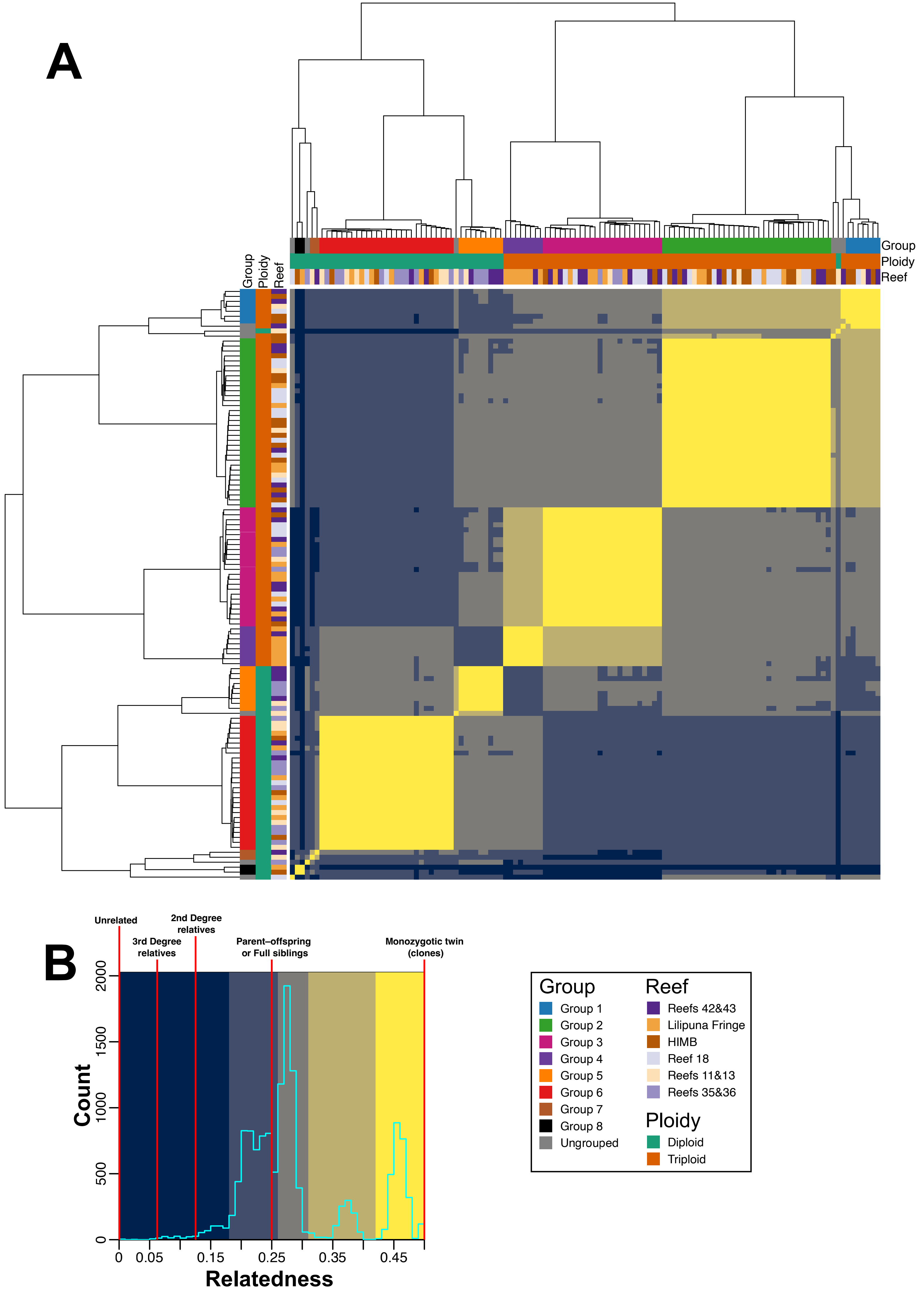

### Figure_4.pdf

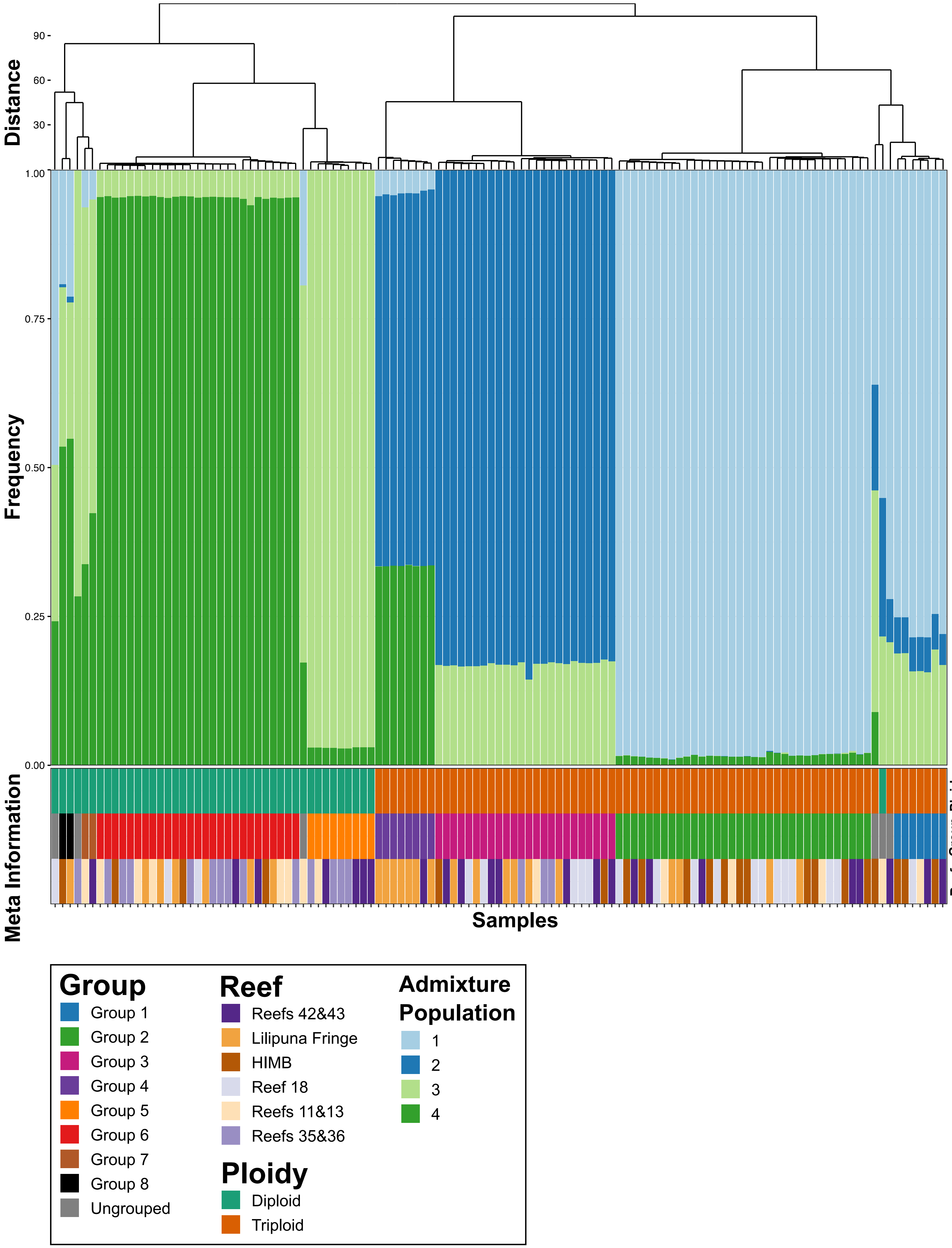

### Figure_5.pdf

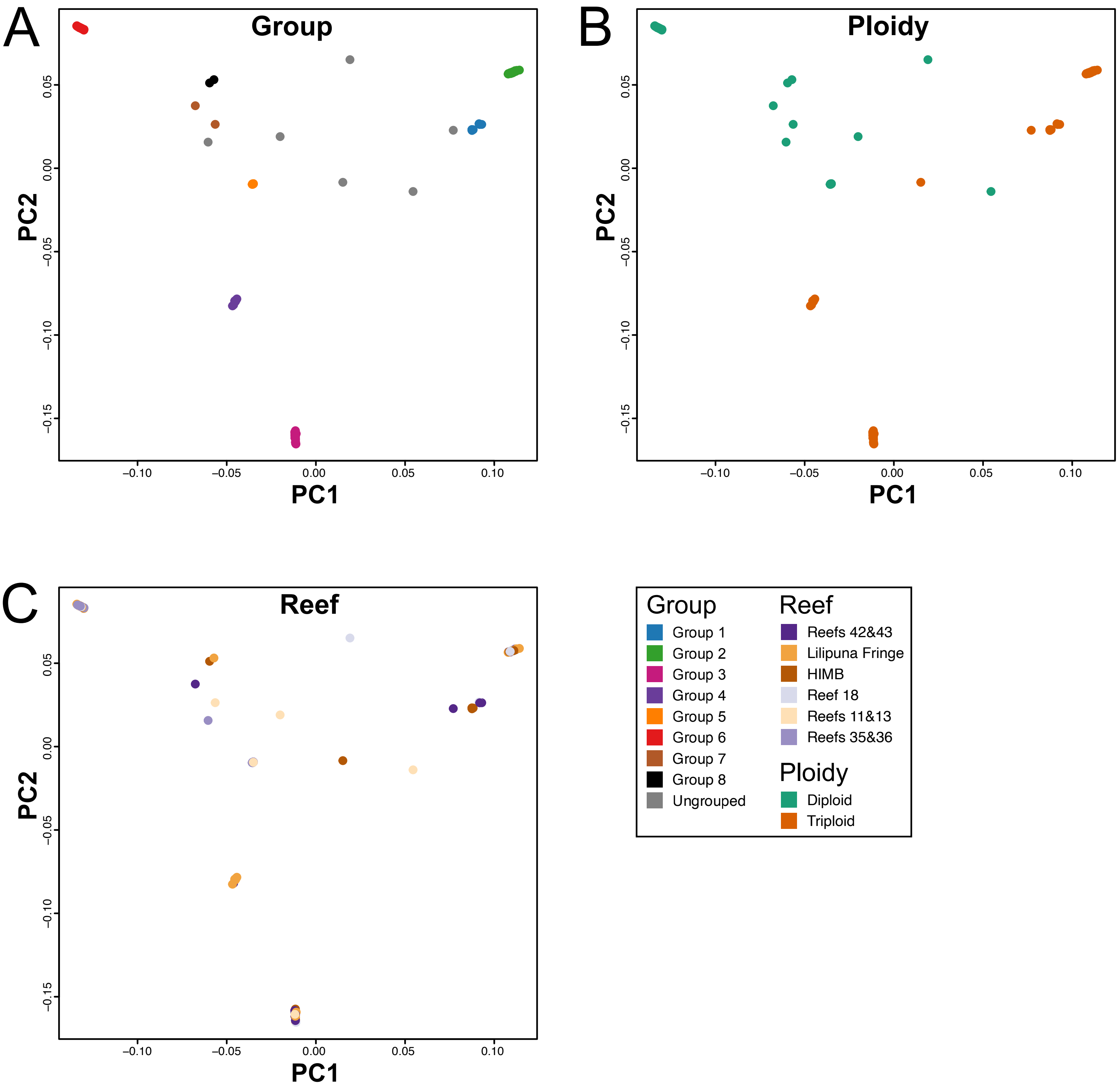

### Figure_6.pdf

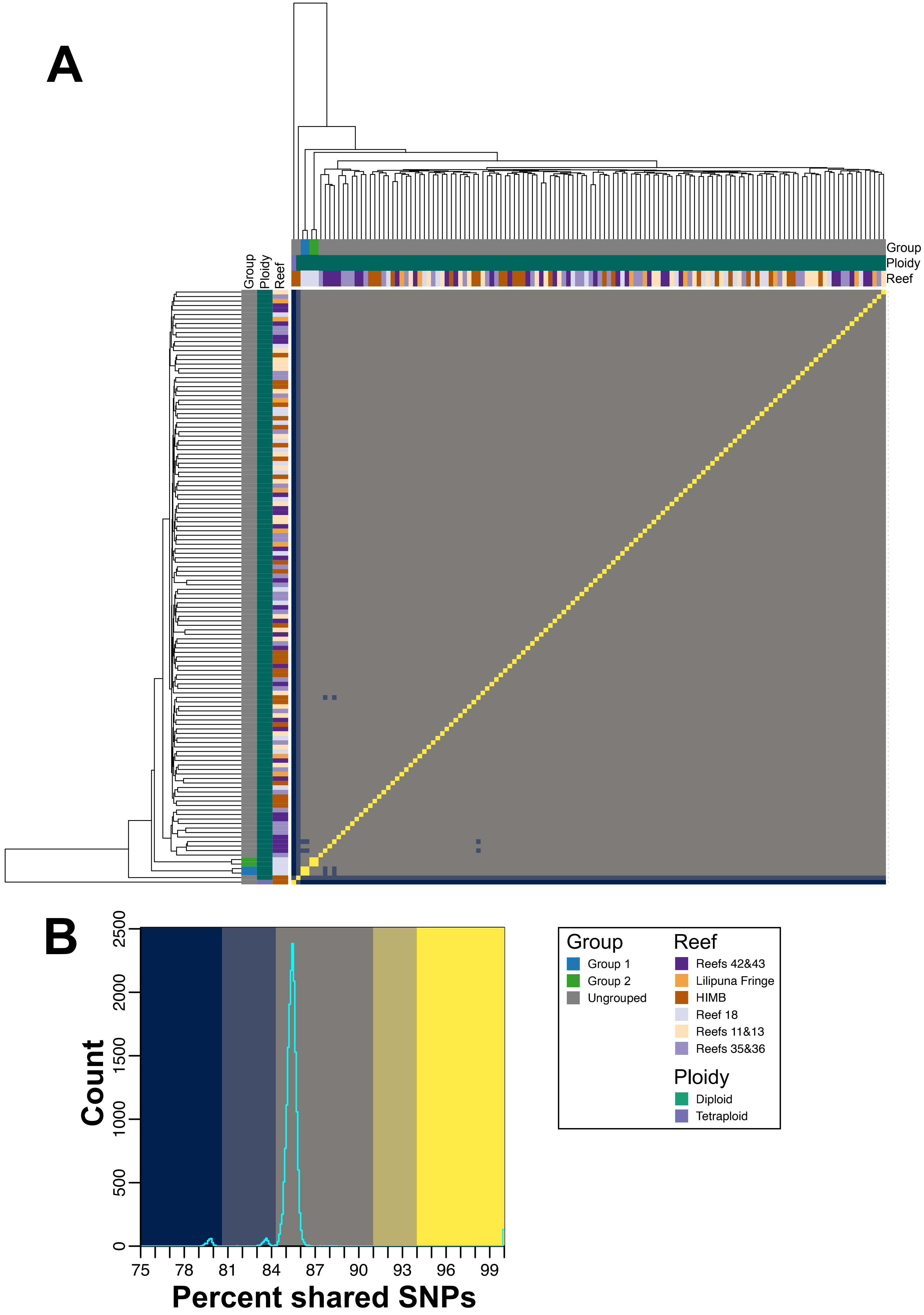

### Figure_7.pdf

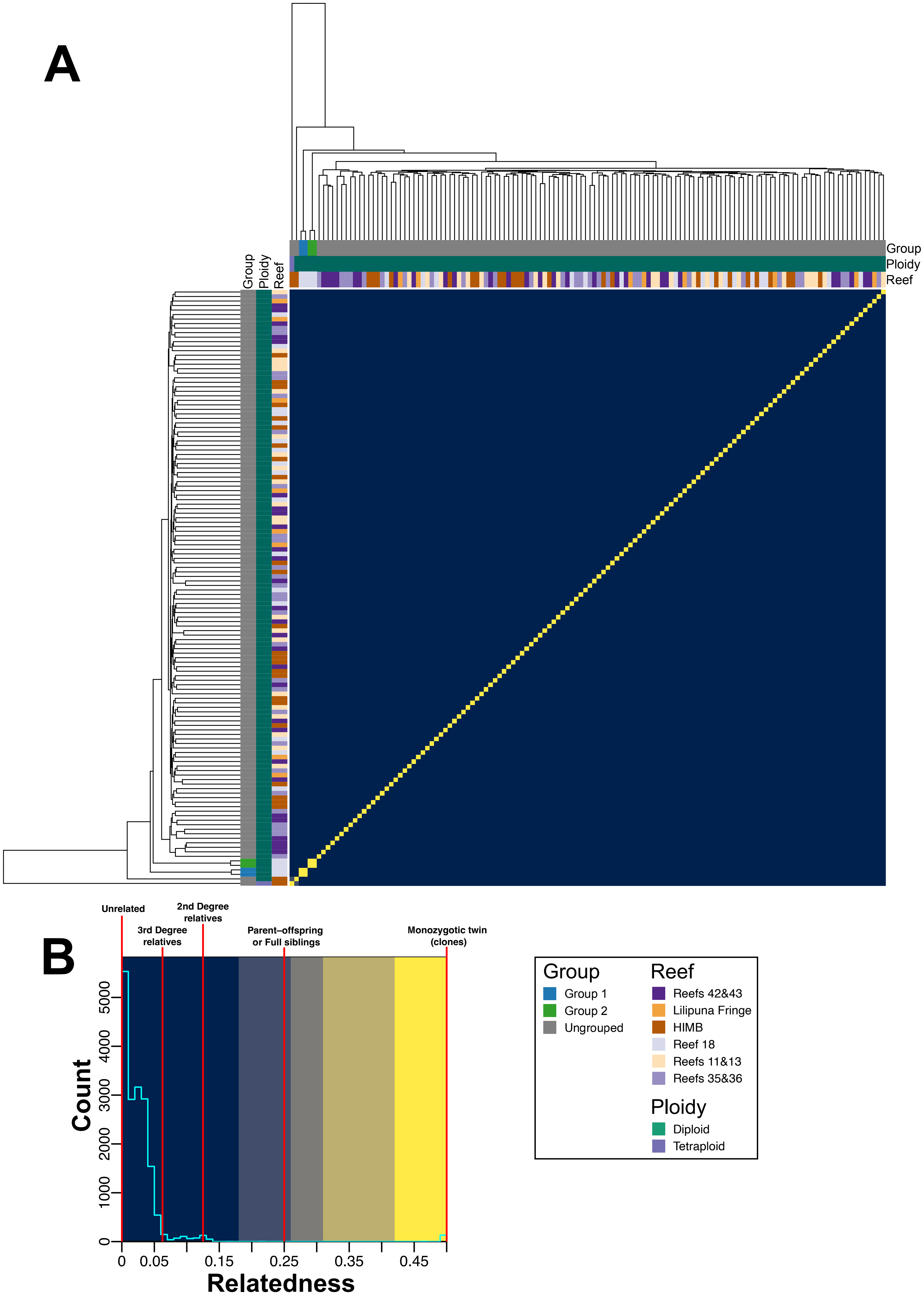

### Figure_S1.pdf

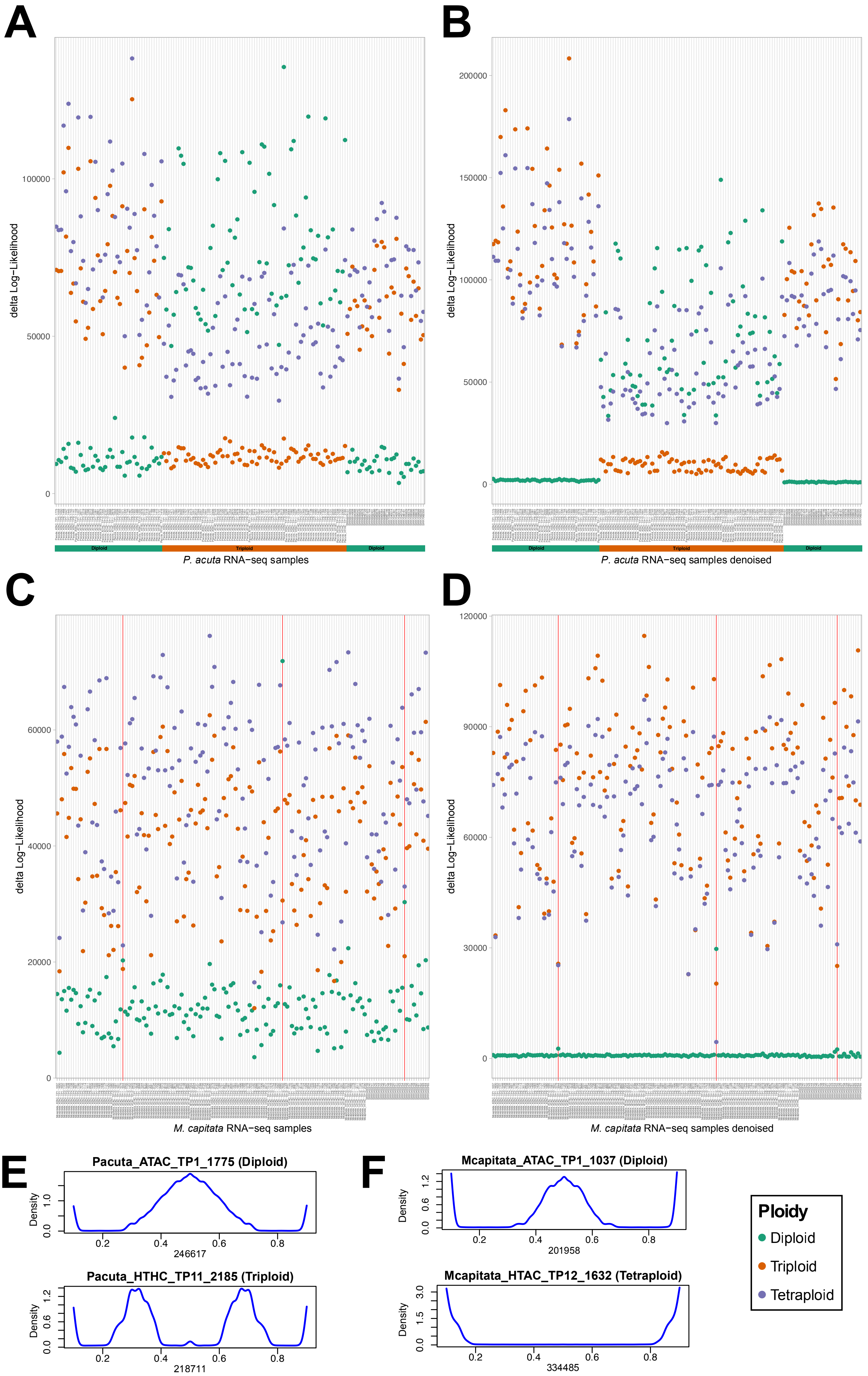

### Figure_S2.pdf

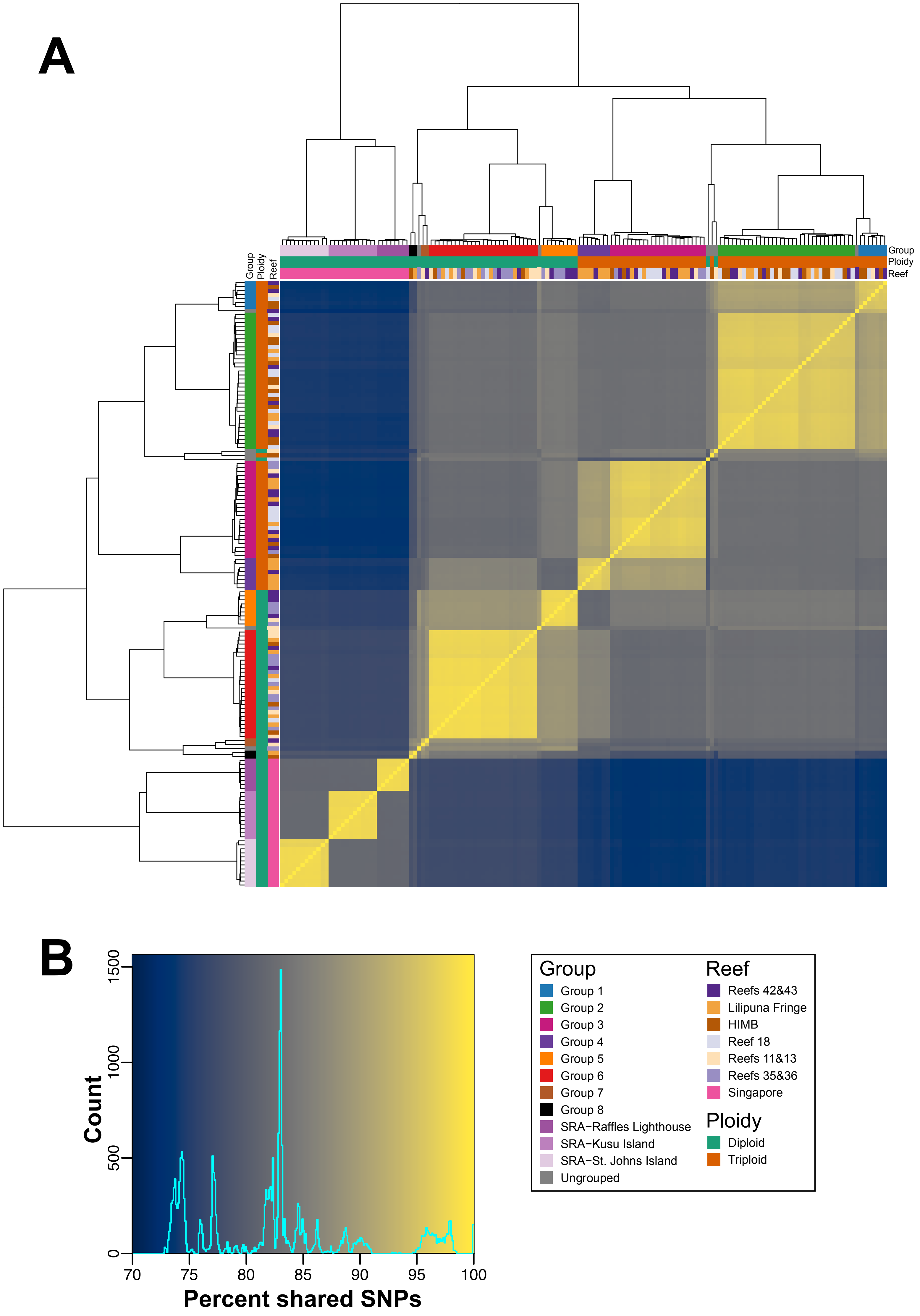

### Figure_S3.pdf

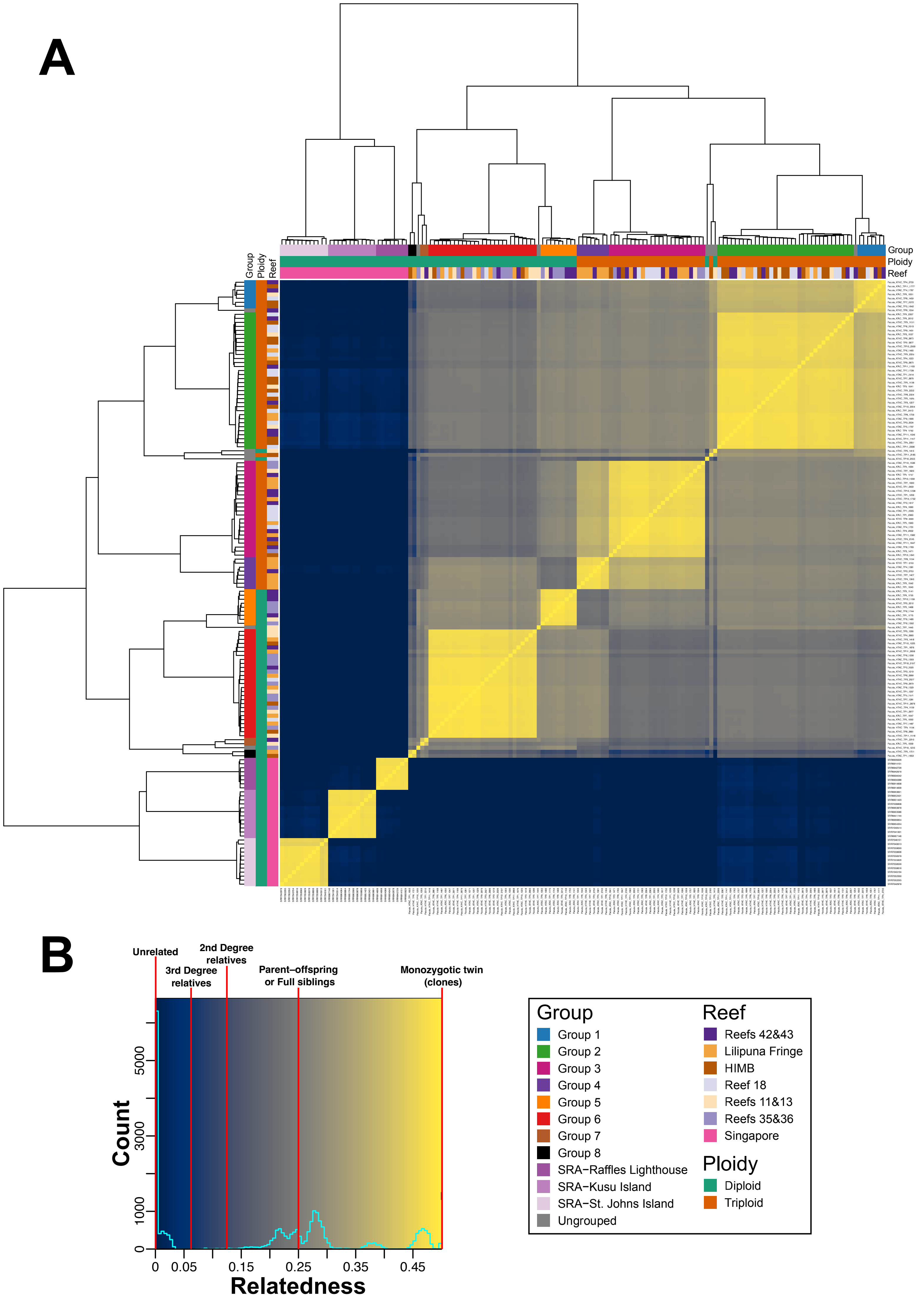

### Figure_S4.pdf

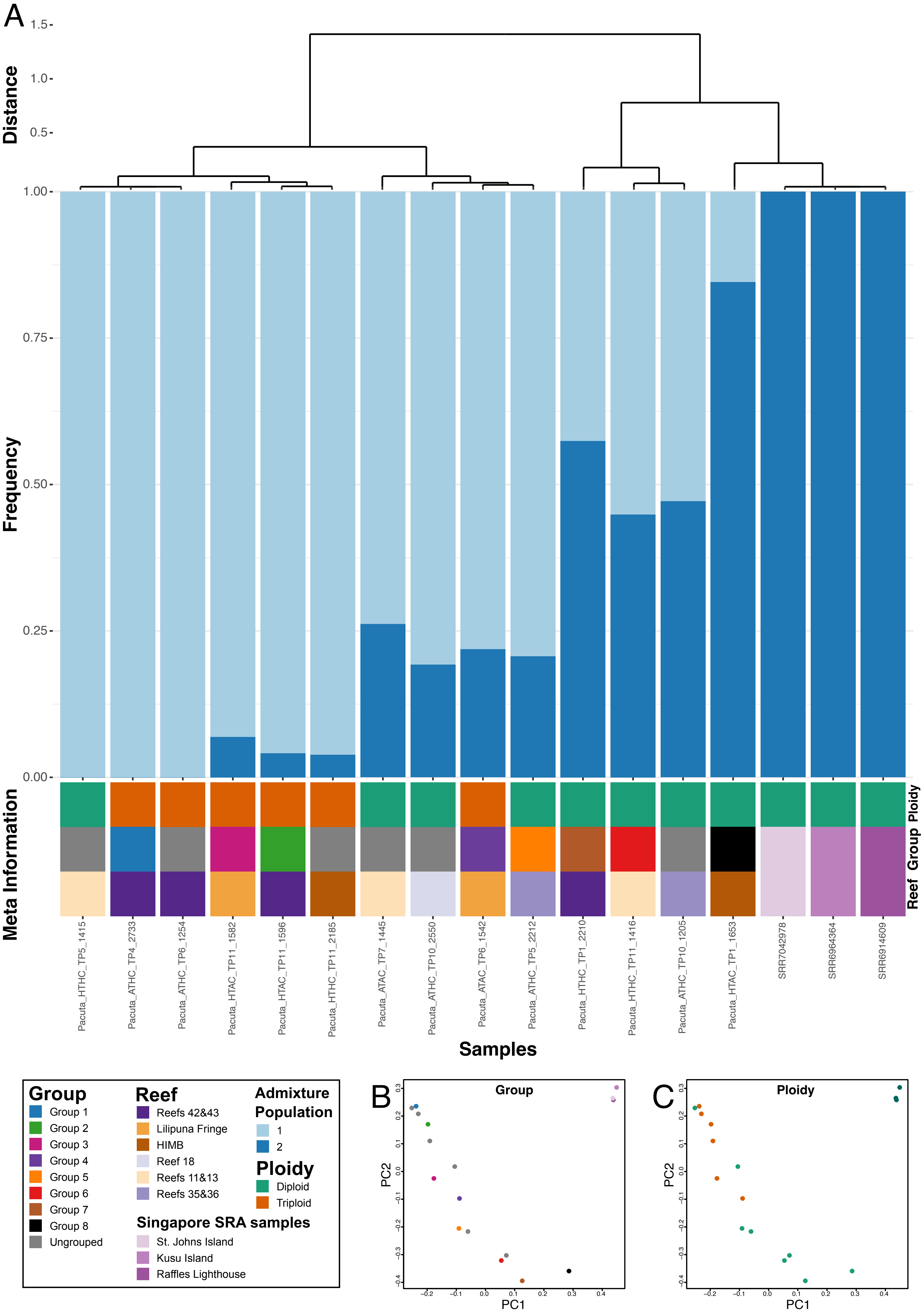

### Figure_S5.pdf

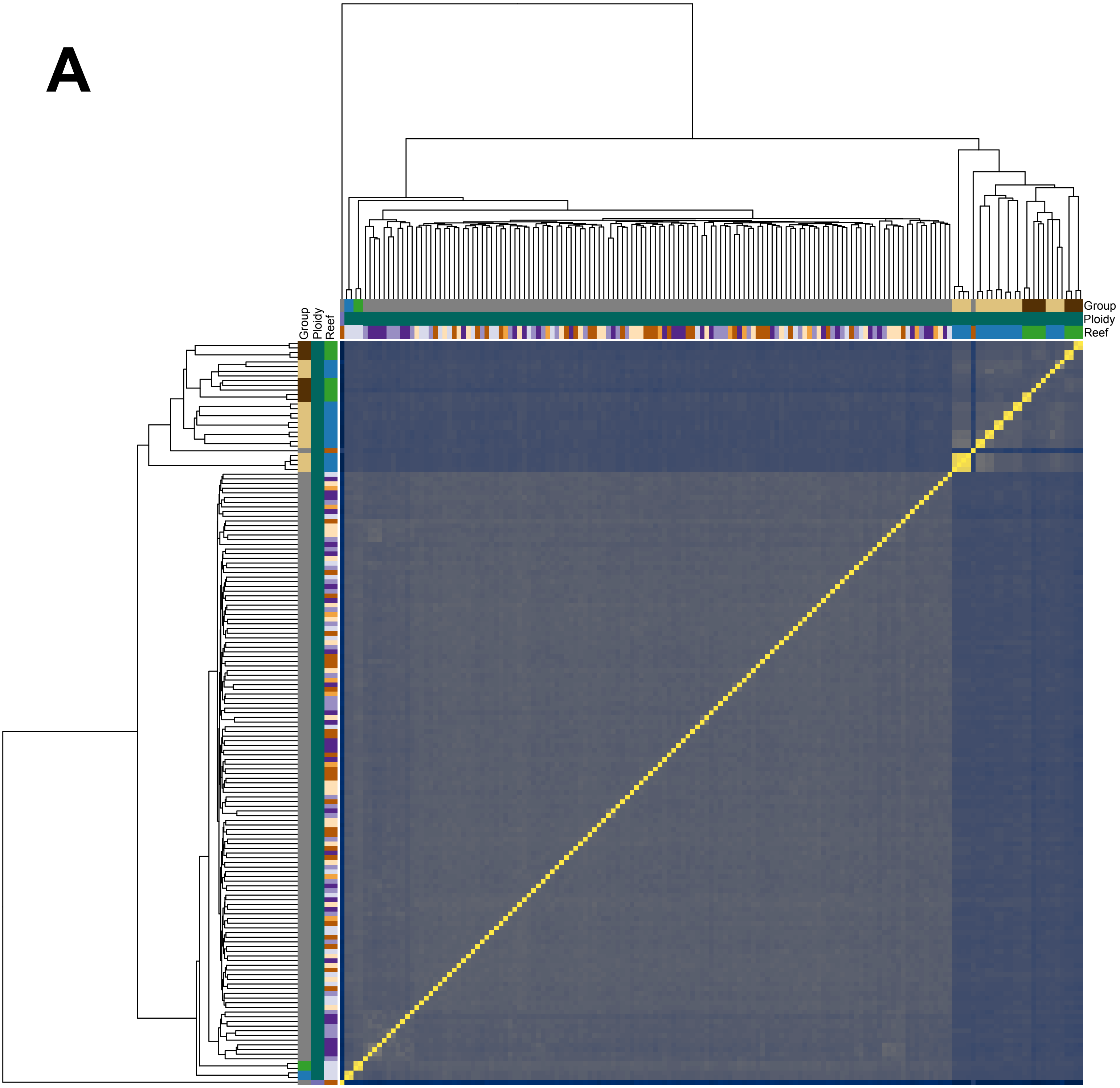

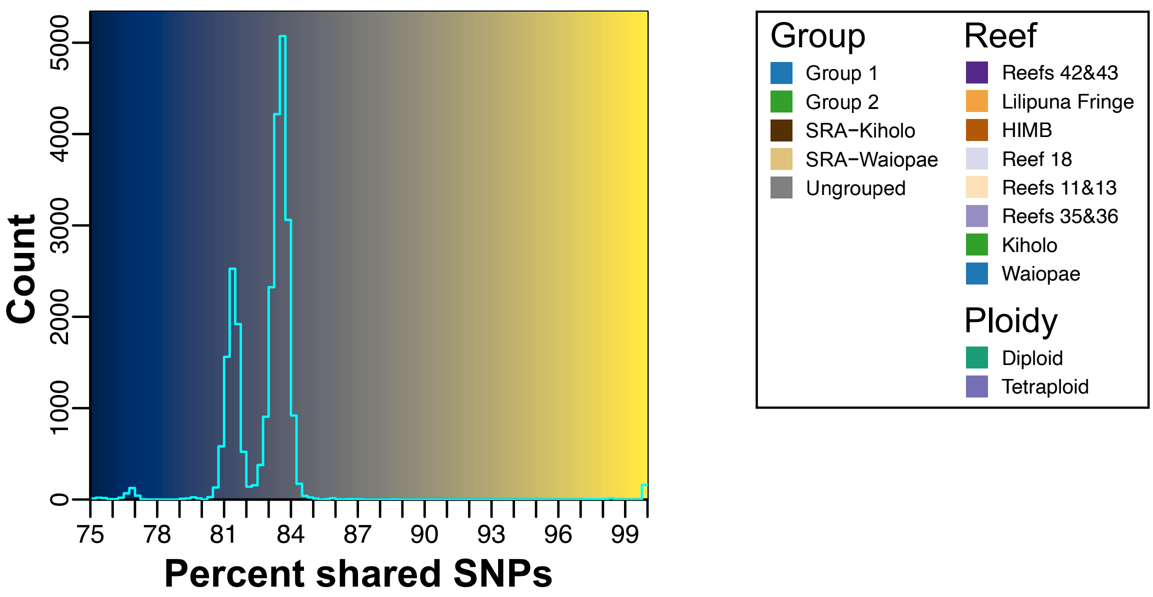

### Figure_S6.pdf

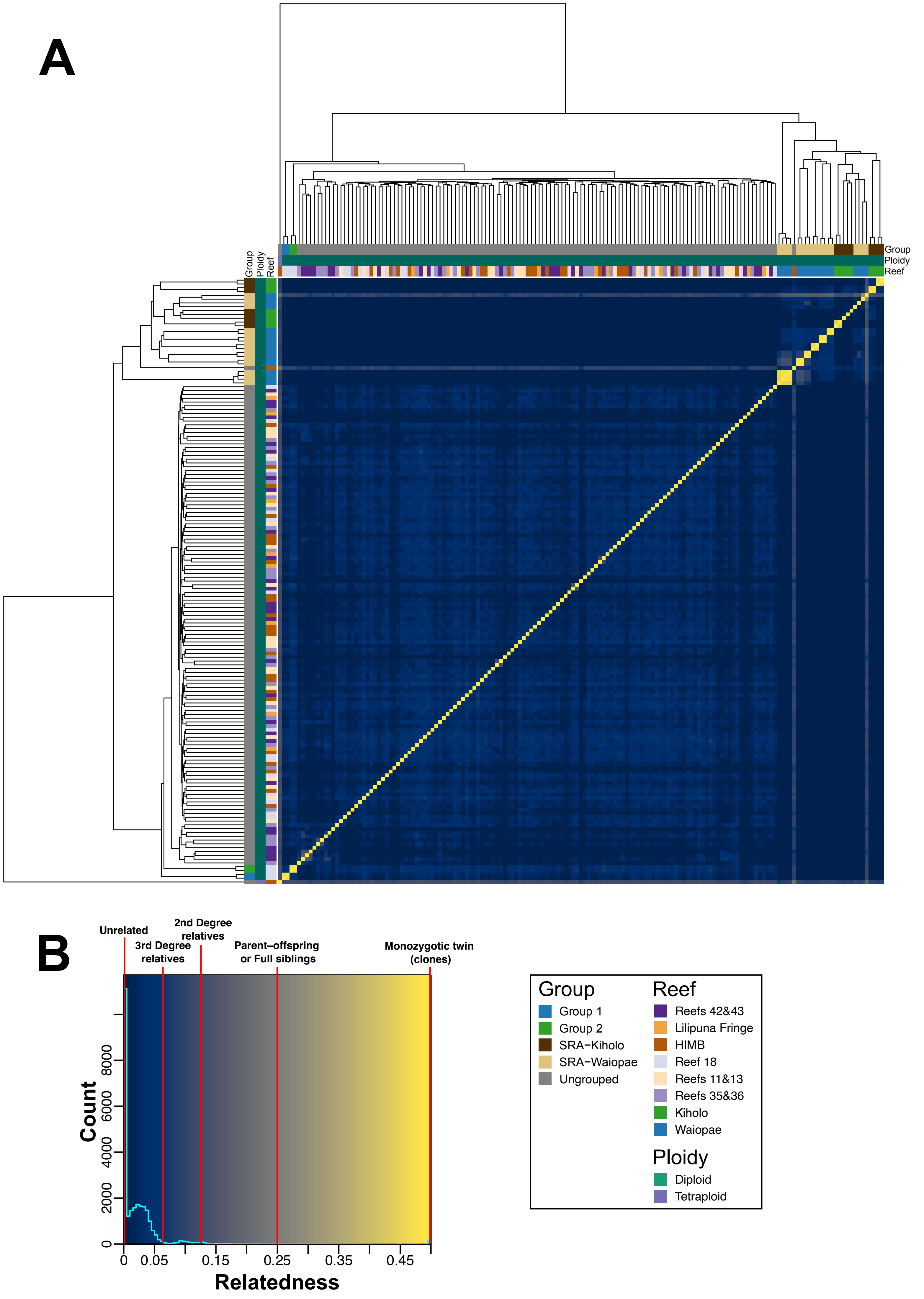

### Figure_S7.pdf

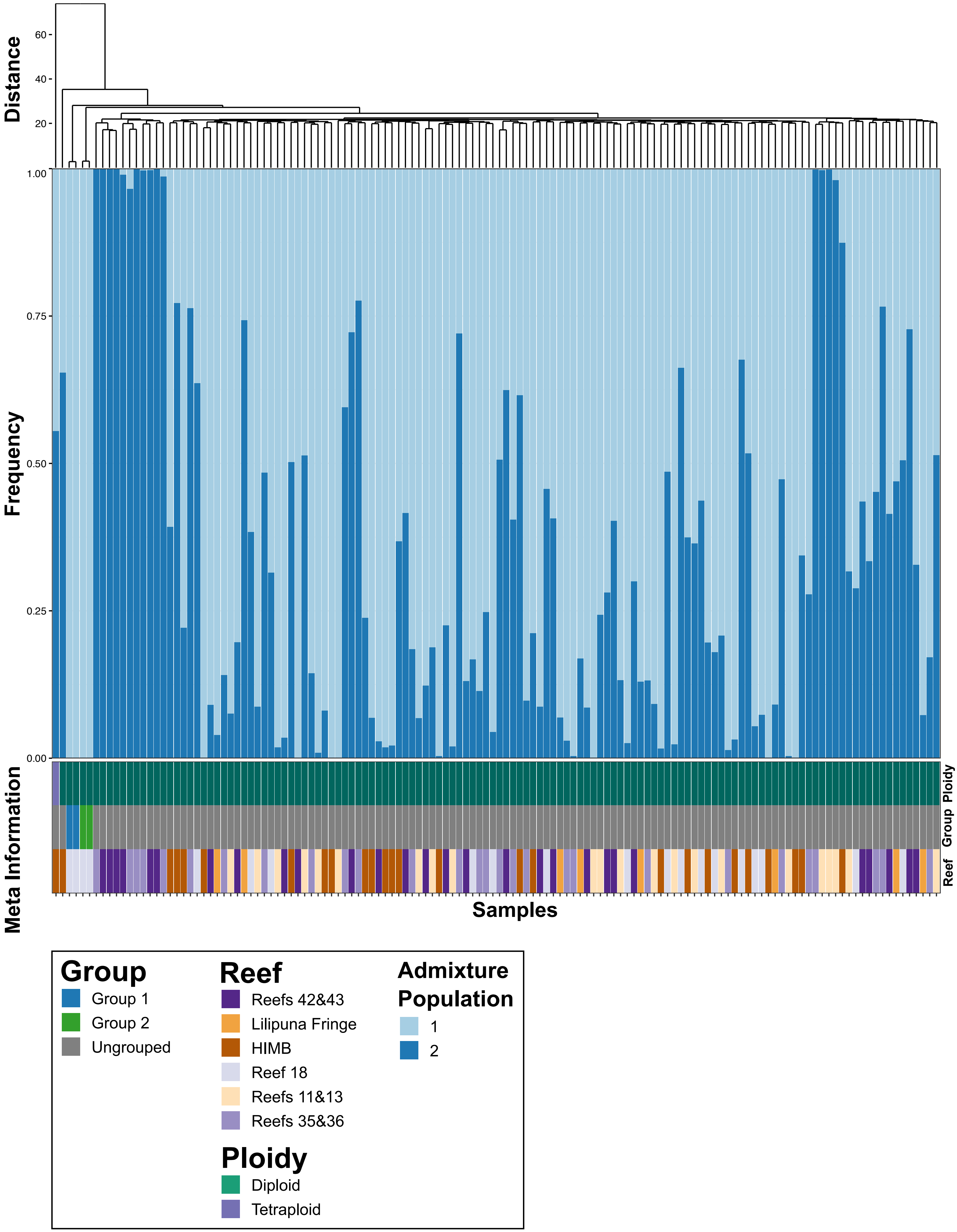

### Figure_S8.pdf

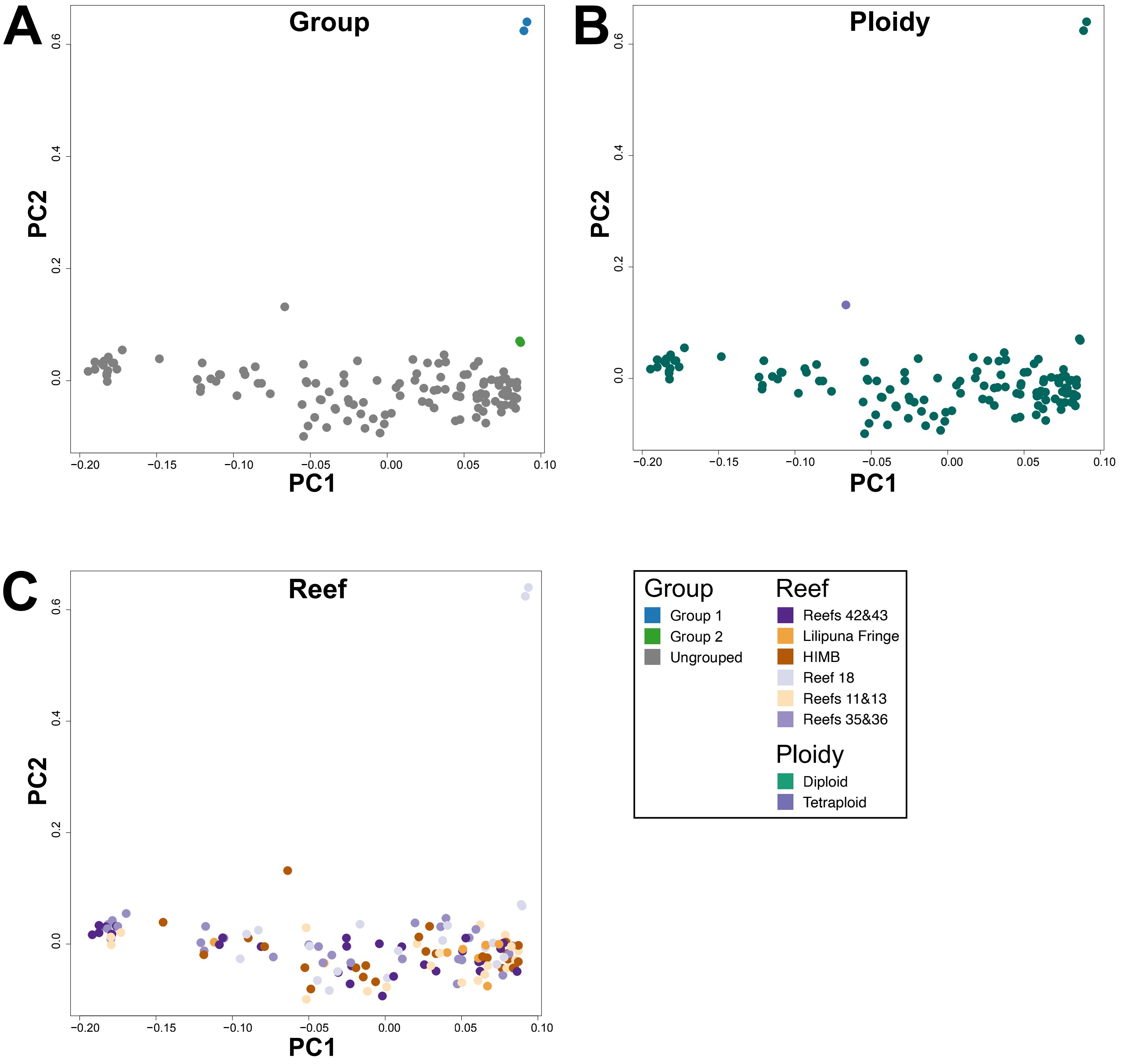
