## Supplementary Figures for "Ploidy variation and its implications for reproduction and population dynamics in two sympatric Hawaiian coral species"

**This PDF file includes:**

Figs. S1 to S8

**Other Supplementary Materials for this manuscript include the following:**

Tables S1 to S8
Data S1 to S4

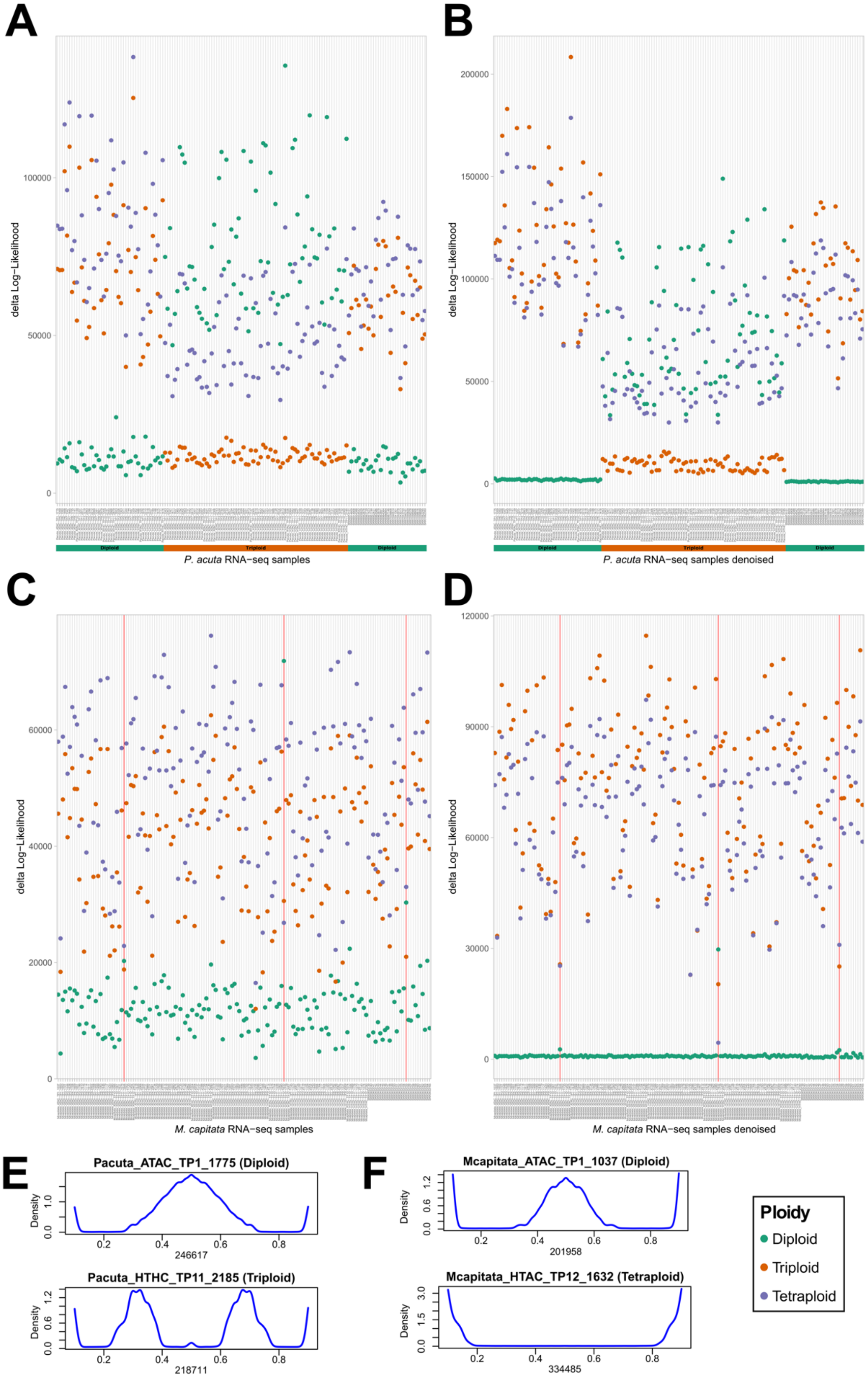
A
B
C
D
E
F
delta Log−Likelihood
P. acuta RNA−seq samples
P. acuta RNA−seq samples denoised
M. capitata RNA−seq samples
M. capitata RNA−seq samples denoised
Diploid
Triploid
Diploid
Pacuta_ATAC_TP1_1775 (Diploid)
Pacuta_HTHC_TP11_2185 (Triploid)
Mcapitata_ATAC_TP1_1037 (Diploid)
Mcapitata_HTAC_TP12_1632 (Tetraploid)
Density
246617
218711
201958
334485
Ploidy
Diploid
Triploid
Tetraploid

**Fig. S1.**

**nQuire delta-logliklihood values for diploid, triploid, and tetraploid models.** Delta-logliklihood values for *P. acuta* RNA-Seq samples (from this study and SRA) (**A**) before and (**B**) after denoising, and the *M. capitata* RNA-Seq samples (this study and SRA) (**C**) before and (**D**) after denoising. Red lines in (**C**) and (**D**) denote samples with unexpected ploidy results (putative samples with multiple genotypes present). A legend explaining the colors is shown in the bottom right of the figure. The colored bars at the bottom of panels (**A**) and (**B**) highlight the predicted ploidy of each sample. (**E**) Density distribution of SNP alleles supported by a given proportion of aligned reads for selected *P. acuta* and *M. capitata* samples.

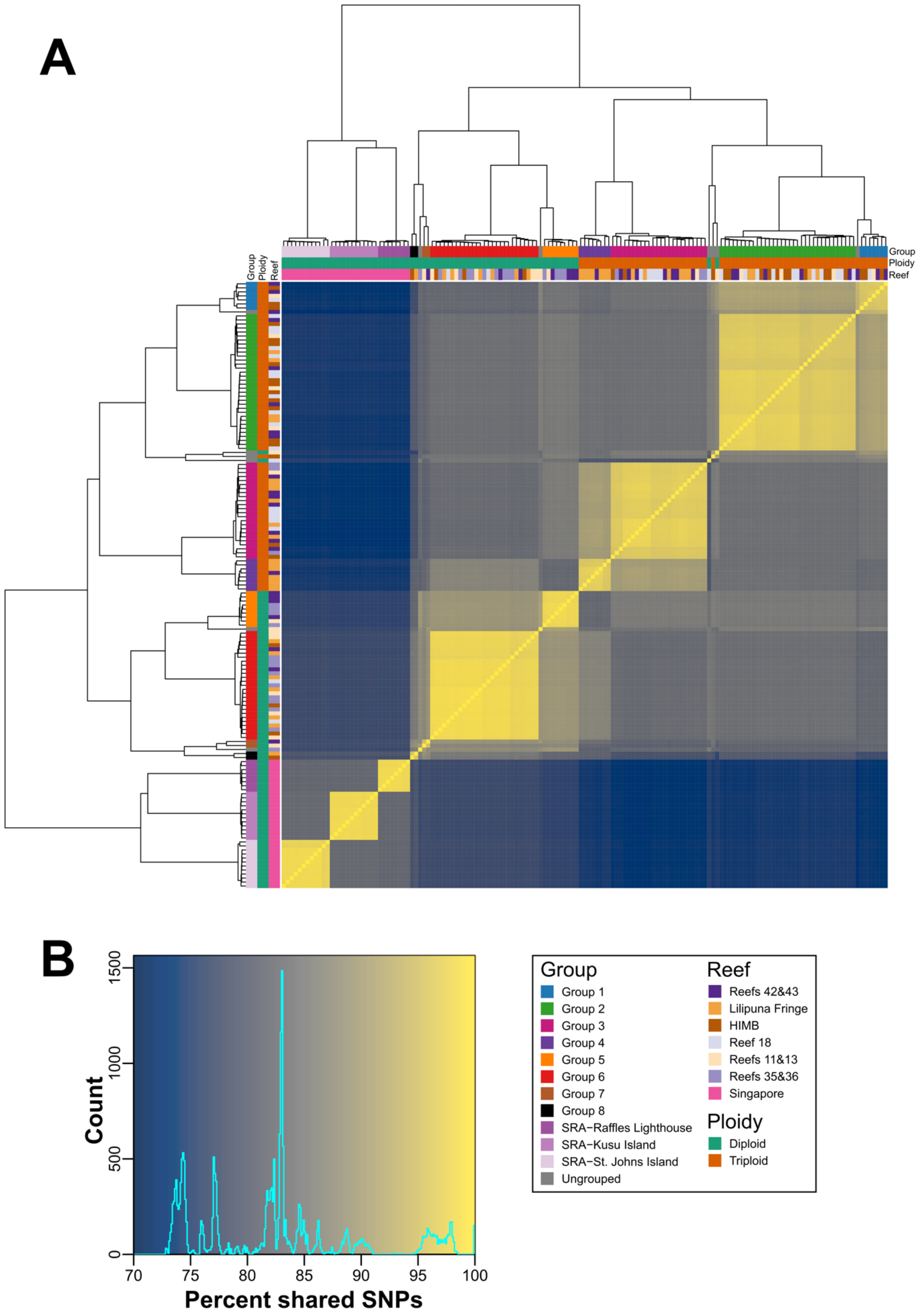
A
Group
Ploidy
Reef
B
Count
Percent shared SNPs
0
500
1000
1500
70
75
80
85
90
95
100
Group
Group 1
Group 2
Group 3
Group 4
Group 5
Group 6
Group 7
Group 8
SRA−Raffles Lighthouse
SRA−Kusu Island
SRA−St. Johns Island
Ungrouped
Reef
Reefs 42&43
Lilipuna Fringe
HIMB
Reef 18
Reefs 11&13
Reefs 35&36
Singapore
Ploidy
Diploid
Triploid

**Fig. S2.**

**Proportion of shared SNPs between each combination of *P. acuta* samples.**

(**A**) Heatmap displaying the proportion of shared SNPs between all pair-wise combinations of *P. acuta* RNA-seq samples sequenced in this study and downloaded from SRA. The putative clonal groups identified in this study (>94% shared SNPs) are shown as colored bars along the top and left sides of the heatmap; a legend describing the color of each group is presented on the bottom right side of the image. The order of the column and rows, and the dendrograms presented on the top and left sides of the heatmap, were generated by hierarchical clustering of the proportion of shared SNPs. (**B**) Histogram of the proportion of shared SNPs in the heatmap; the background colors used in the histogram correspond to the colors used in the heatmap presented in (**A**).

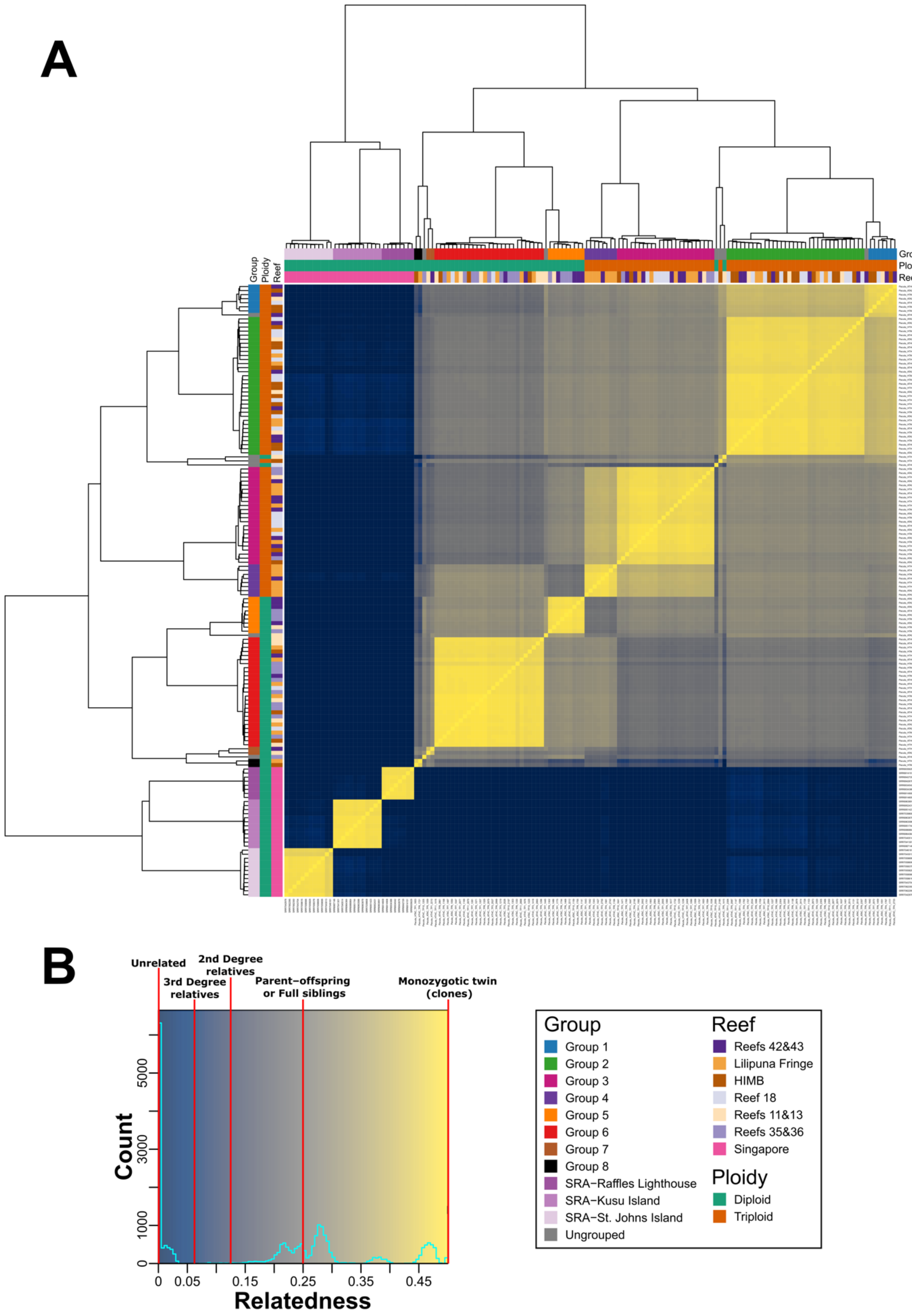
A
Group
Ploidy
Reef
B
Unrelated
2nd Degree relatives
3rd Degree relatives
Parent–offspring or Full siblings
Monozygotic twin (clones)
Count
5000
3000
1000
0
0
0.05
0.15
0.25
0.35
0.45
Relatedness
Group
Group 1
Group 2
Group 3
Group 4
Group 5
Group 6
Group 7
Group 8
SRA–Raffles Lighthouse
SRA–Kusu Island
SRA–St. Johns Island
Ungrouped
Reef
Reefs 42&43
Lilipuna Fringe
HIMB
Reef 18
Reefs 11&13
Reefs 35&36
Singapore
Ploidy
Diploid
Triploid

**Fig. S3.**

**Pair-wise relatedness between each combination of *P. acuta* samples.**

(**A**) Heatmap displaying the relatedness scores (proposed by Manichaikul, et al. 2010) of all pair-wise combinations of *P. acuta* RNA-seq samples sequenced in this study and downloaded from SRA. The putative clonal groups identified in this study are shown as colored bars along the top and left sides of the heatmap; a legend describing the color of each group is presented on the bottom right side of the image. The order of the column and rows, and the dendrograms presented on the top and left sides of the heatmap, are based on the dendrogram used in Figure S2. (**B**) Histogram of relatedness scores in the heatmap; the background colors used in the histogram correspond to the colors used in the heatmap presented in (**A**). The relatedness values that correspond to a particular relationship between samples (described in Manichaikul *et al.* (*13*)) are annotated on the histogram using vertical red lines. The colors used in the heatmap and histogram were manually chosen to highlight the distinct sets of relatedness scores observed in the histogram presented in (**B**).

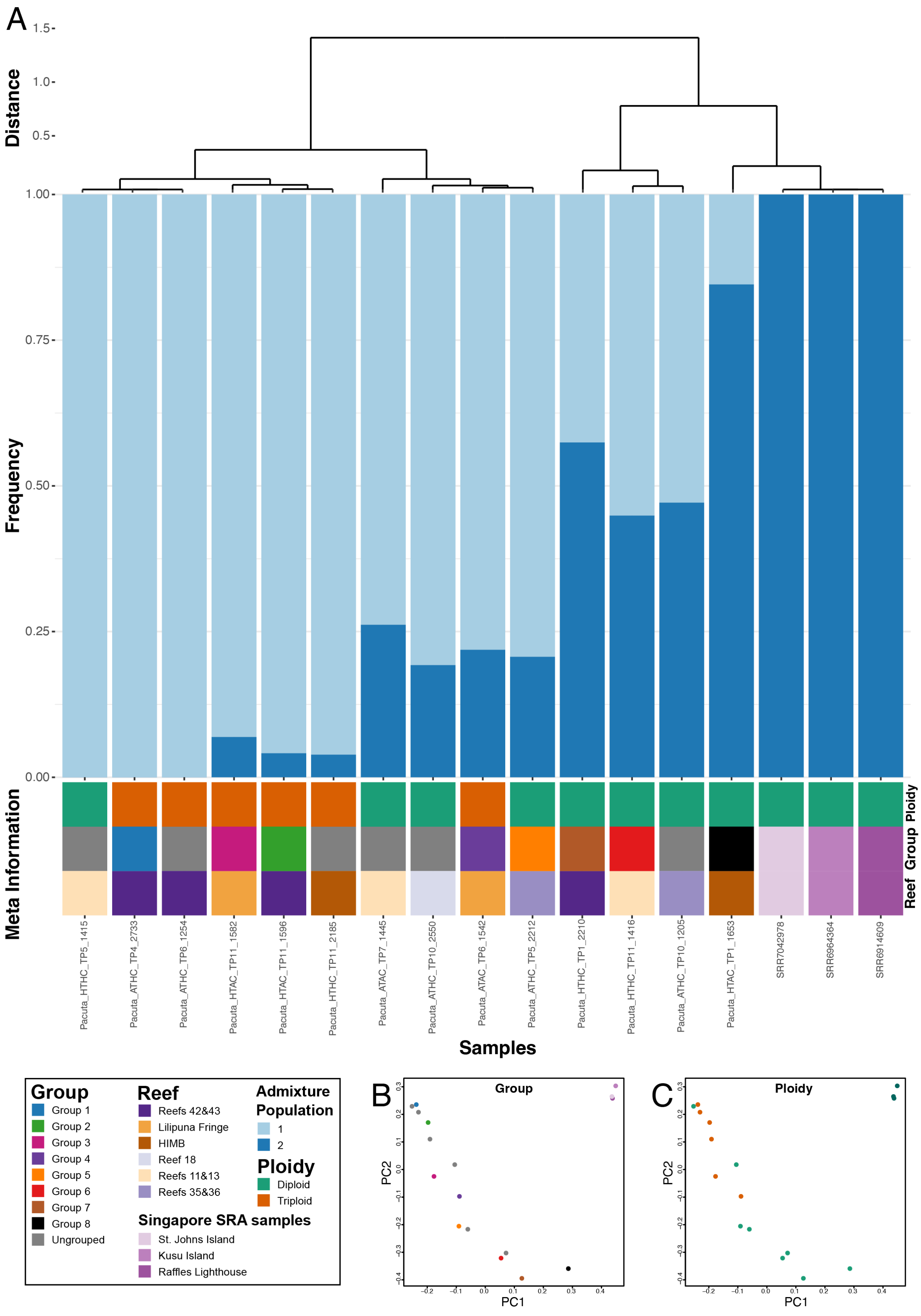
A
Distance
1.5
1.0
0.5
Frequency
1.00
0.75
0.50
0.25
0.00
Meta Information
Ploidy
Group
Reef
Pacuta_HTHC_TP5_1415
Pacuta_ATHC_TP4_2733
Pacuta_ATHC_TP6_1254
Pacuta_HTAC_TP11_1582
Pacuta_HTAC_TP11_1596
Pacuta_HTHC_TP11_2185
Pacuta_ATAC_TP7_1445
Pacuta_ATHC_TP10_2550
Pacuta_ATAC_TP6_1542
Pacuta_ATHC_TP5_2212
Pacuta_HTHC_TP1_2210
Pacuta_HTHC_TP11_1416
Pacuta_ATHC_TP10_1205
Pacuta_HTAC_TP1_1653
SRR7042978
SRR6964364
SRR6914609
Samples
Group
Group 1
Group 2
Group 3
Group 4
Group 5
Group 6
Group 7
Group 8
Ungrouped
Reef
Reefs 42&43
Lilipuna Fringe
HIMB
Reef 18
Reefs 11&13
Reefs 35&36
Singapore SRA samples
St. Johns Island
Kusu Island
Raffles Lighthouse
Admixture Population
1
2
Ploidy
Diploid
Triploid
B
Group
PC2
PC1
C
Ploidy
PC2
PC1

**Fig. S4.**
**Admixture and PCA analysis of *P. acuta* samples with a single representative per clonal group.**
(A) A stacked bar chart showing the proportion of each *P. acuta* sample's ancestry that is from each of the two estimated (by PCAngsd) ancestral populations. The ploidy, putative clonal group, and reef that the sample was collected from are shown as colored bars at the bottom of the plot. The dendrogram and order of the columns is based on the hierarchical clustering of the admixture results. The representative samples were selected based on their mapping rates to the reference genome, that is, the selected samples had the highest proportions of mapped reads amongst all samples from that clonal group. PCA of *P. acuta* samples colored by (B) group and (C) ploidy. Plots are based on the covariance matrix produced by PCAngsd with estimated individual allele frequencies and show PC1 (24.87% variance explained) and PC2 (7.29%). A legend describing each of the colors used in the figure is presented at the bottom of the plot.

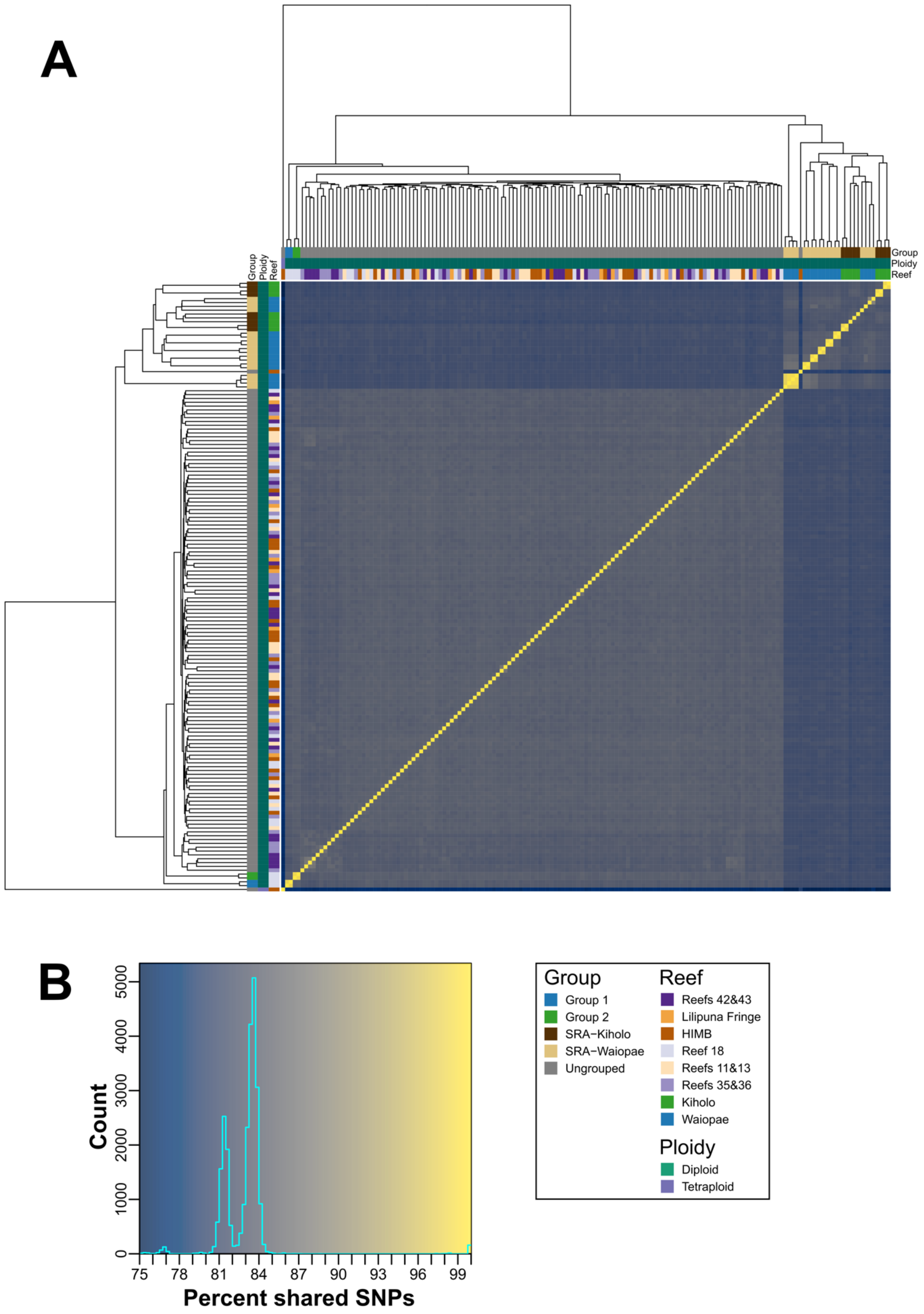
A
Group
Ploidy
Reef
B
Count
Percent shared SNPs
Group
Group 1
Group 2
SRA−Kiholo
SRA−Waiopae
Ungrouped
Reef
Reefs 42&43
Lilipuna Fringe
HIMB
Reef 18
Reefs 11&13
Reefs 35&36
Kiholo
Waiopae
Ploidy
Diploid
Tetraploid

**Fig. S5.**

**Proportion of shared SNPs between each combination of *M. capitata* samples.**

(**A**) Heatmap displaying the proportion of shared SNPs between all pair-wise combinations of *M. capitata* RNA-seq samples sequenced in this study and downloaded from SRA. The putative clonal groups identified in this study (>94% shared SNPs) are shown as colored bars along the top and left sides of the heatmap; a legend describing the color of each group is presented on the bottom right side of the image. The order of the column and rows, and the dendrograms presented on the top and left sides of the heatmap, were generated by hierarchical clustering of the proportion of shared SNPs. (**B**) Histogram of the proportion of shared SNPs in the heatmap; the background colors used in the histogram correspond to the colors used in the heatmap presented in (**A**).

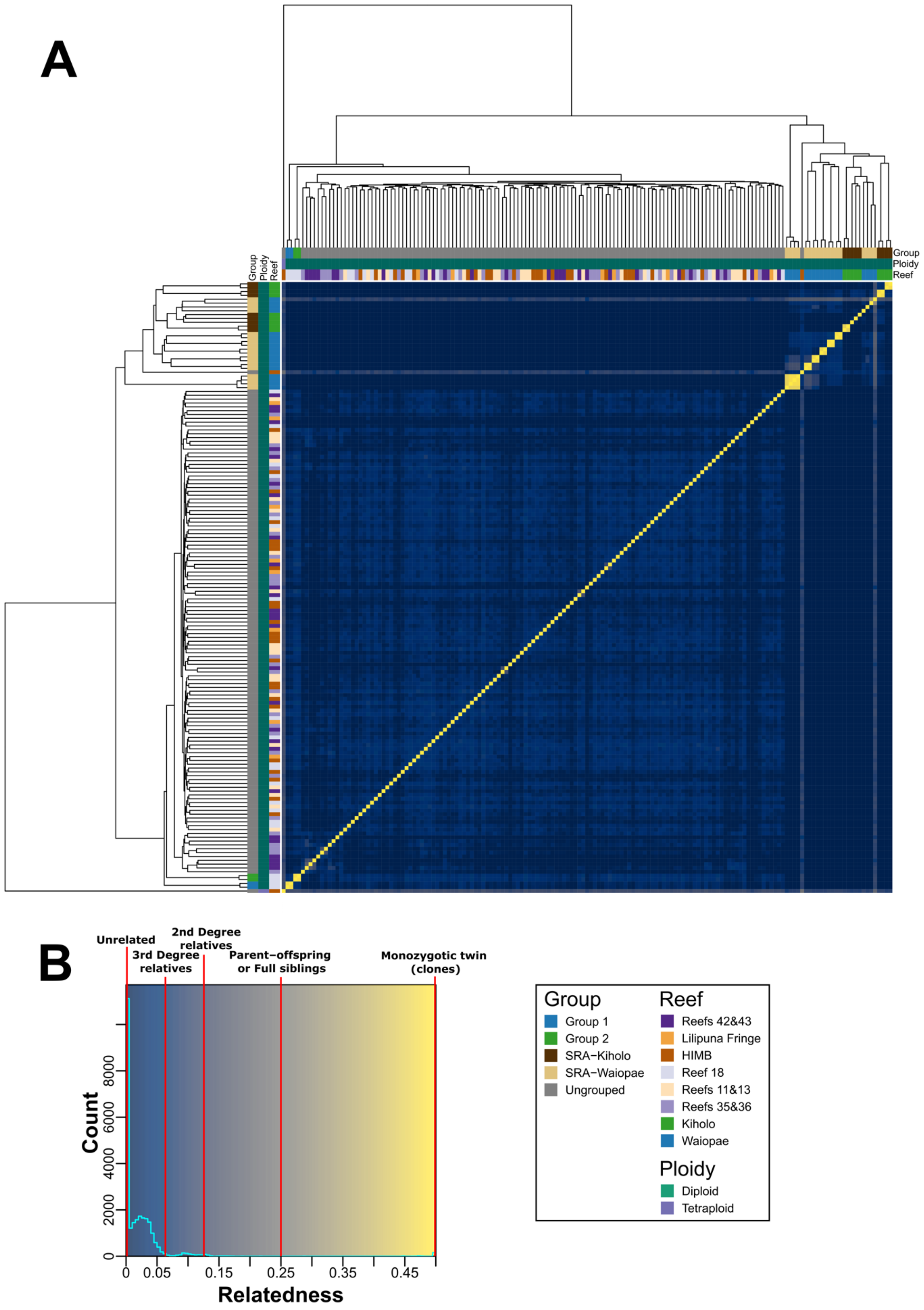
A
Group
Ploidy
Reef
B
Unrelated
3rd Degree relatives
2nd Degree relatives
Parent−offspring or Full siblings
Monozygotic twin (clones)
Count
0
2000
4000
6000
8000
0
0.05
0.15
0.25
0.35
0.45
Relatedness
Group
Group 1
Group 2
SRA−Kiholo
SRA−Waiopae
Ungrouped
Reef
Reefs 42&43
Lilipuna Fringe
HIMB
Reef 18
Reefs 11&13
Reefs 35&36
Kiholo
Waiopae
Ploidy
Diploid
Tetraploid

**Fig. S6.**

**Pair-wise relatedness between each combination of *M. capitata* samples.**

(**A**) Heatmap displaying the relatedness scores (proposed by Manichaikul, et al. 2010) of all pair-wise combinations of *M. capitata* RNA-seq samples sequenced in this study and downloaded from SRA. The putative clonal groups identified in this study are shown as colored bars along the top and left sides of the heatmap; a legend describing the color of each group is presented on the bottom right side of the image. The order of the column and rows, and the dendrograms presented on the top and left sides of the heatmap, are based on the dendrogram used in Figure S5. (**B**) Histogram of relatedness scores in the heatmap; the background colors used in the histogram correspond to the colors used in the heatmap presented in (**A**). The relatedness values that correspond to a particular relationship between samples (described in Manichaikul, et al. 2010) are annotated on the histogram using vertical red lines. The colors used in the heatmap and histogram were manually chosen to highlight the distinct sets of relatedness scores observed in the histogram presented in (**B**).

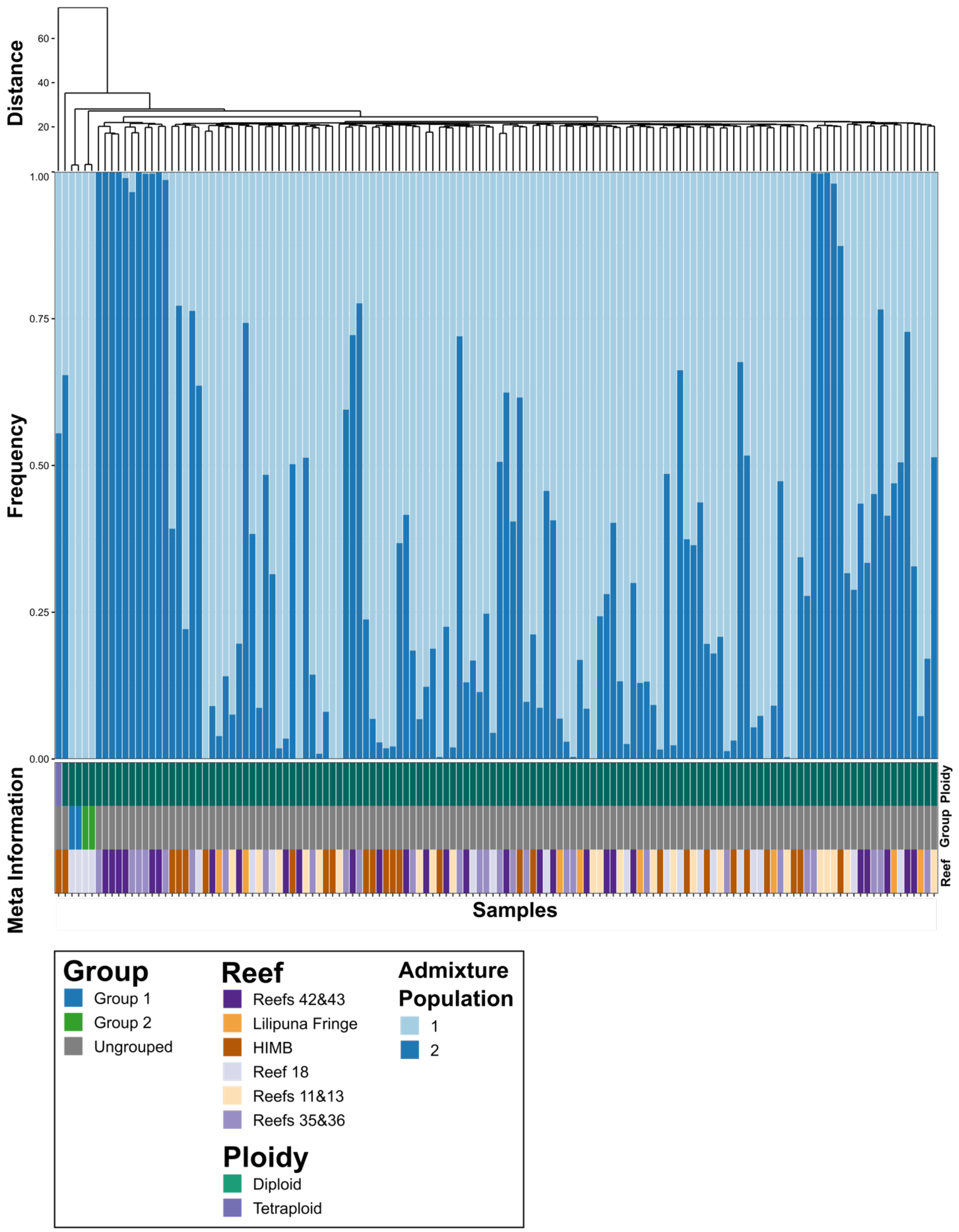
Distance
60
40
20
Frequency
1.00
0.75
0.50
0.25
0.00
Meta Information
Ploidy
Group
Reef
Samples
Group
Group 1
Group 2
Ungrouped
Reef
Reefs 42&43
Lilipuna Fringe
HIMB
Reef 18
Reefs 11&13
Reefs 35&36
Ploidy
Diploid
Tetraploid
Admixture Population
1
2

**Fig. S7.**
**Admixture analysis of *M. capitata* samples.**
A stacked bar chart showing the proportion of each *M. capitata* sample's ancestry that is from each of the two estimated (by PCAngsd) ancestral populations. The ploidy, putative clonal group, and reef that the sample was collected from are shown as colored bars at the bottom of the plot; a legend describing each of the colors used in the figure is presented at the bottom of the plot. The dendrogram and order of the columns is based on the hierarchical clustering of the allelic similarity scores presented in Figure 6.

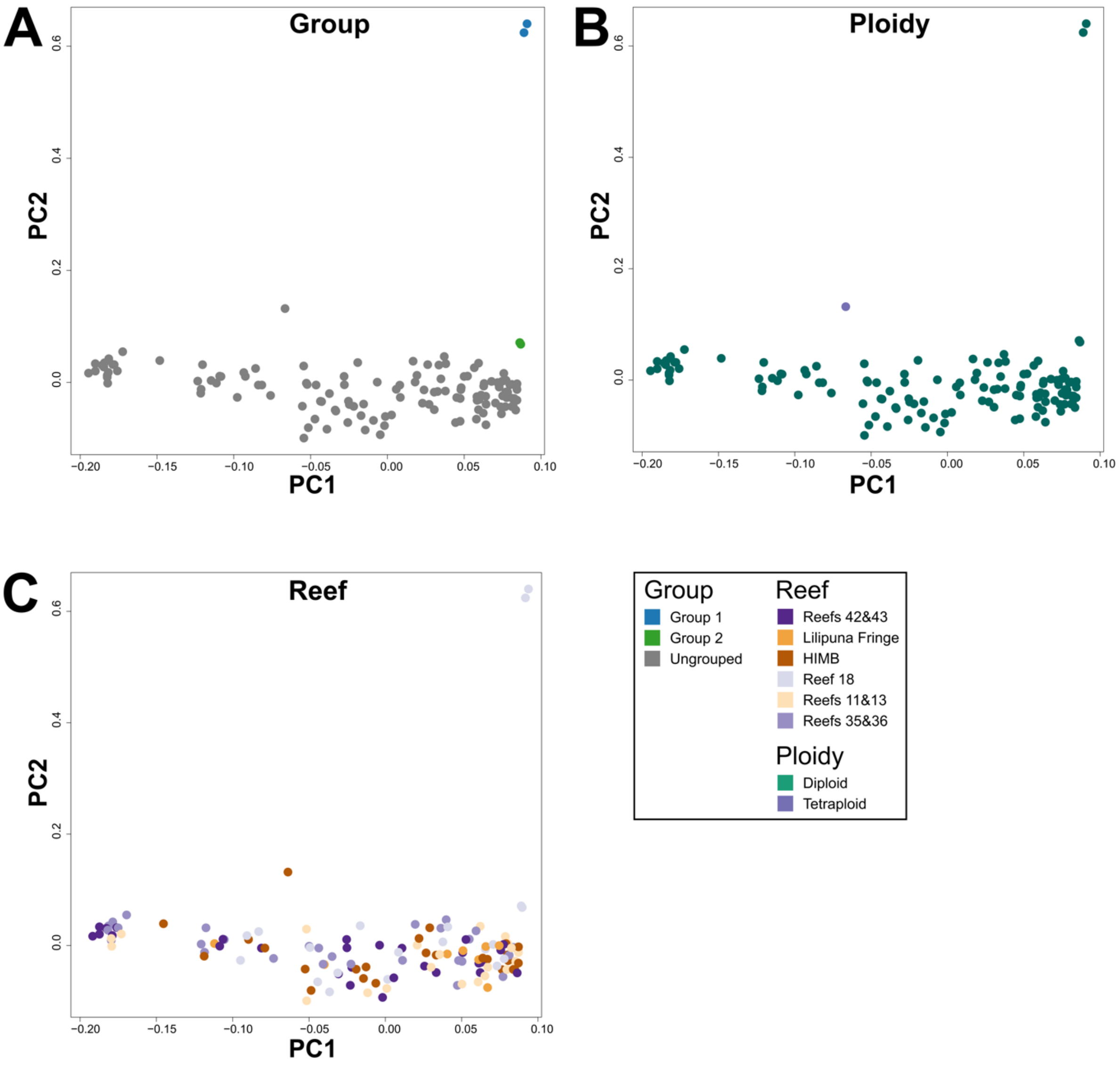
A
Group
B
Ploidy
C
Reef
PC1
PC2
0.0
0.2
0.4
0.6
−0.20
−0.15
−0.10
−0.05
0.00
0.05
0.10
Group
Group 1
Group 2
Ungrouped
Reef
Reefs 42&43
Lilipuna Fringe
HIMB
Reef 18
Reefs 11&13
Reefs 35&36
Ploidy
Diploid
Tetraploid

**Fig. S8.**
**PCA of *M. capitata* sample relatedness.**
PCA of *M. capitata* samples colored by (**A**) group, (**B**) ploidy, and (**C**) reef. A legend describing each color is presented in the bottom right of the figure. Plots are based on the covariance matrix produced by PCAngsd with estimated individual allele frequencies and show PC1 (9.54% variance explained) and PC2 (0.76%).

**Table S1. (separate file)**
Information on each coral fragment what was included in the RNA-Seq analysis.

**Table S2. (separate file)**
Statistics for the raw and quality-controlled RNA-seq data generated from each sample.

**Table S3. (separate file)**
nQuire delta log-likelihood values for each sample.

**Table S4. (separate file)**
SRA metadata and read counts for additional samples for *P. acuta* and *M. capitata* from SRA that were not collected from Kāne'ohe Bay.

**Table S5. (separate file)**
Proportion of shared SNPs between each pairwise combination of *P. acuta* samples from this study and SRA.

**Table S6. (separate file)**
Relatedness values between each pairwise combination of *P. acuta* samples from this study and SRA.

**Table S7. (separate file)**
Proportion of shared SNPs between each pairwise combination of *M. capitata* samples from this study and SRA.

**Table S8. (separate file)**
Relatedness values between each pairwise combination of *M. capitata* samples from this study and SRA.

**Data S1. (separate file)**
Density distribution of SNP alleles, from nQuire denoised bi-allelic sites, supported by a given proportion of aligned reads for the 119 *P. acuta* RNA-Seq libraries that were analyzed in our study. For each plot, the x-axis is the proportion of reads that support a given SNP allele out of all reads that align to that site, and the y-axis is the frequency of SNP alleles with a given proportion of supporting reads. The name and putative ploidy of each sample is presented above its distribution; the number of denoised bi-allelic sites analyzed in each sample is presented below its distribution. The x-axis of these plots was limited to between 0.1 and 0.9 to allow for clear visualization of the main distribution of sites.

**Data S2. (separate file)**
Density distribution of SNP alleles, from nQuire denoised bi-allelic sites, supported by a given proportion of aligned reads for the 32 *P. acuta* RNA-Seq libraries (not from Kāne'ohe Bay) that were downloaded from SRA. For each plot, the x-axis is the proportion of reads that support a given SNP allele out of all reads that align to that site, and the y-axis is the frequency of SNP alleles with a given proportion of supporting reads. The name and putative ploidy of each sample

is presented above its distribution; the number of denoised bi-allelic sites analyzed in each sample is presented below its distribution. The x-axis of these plots was limited to between 0.1 and 0.9 to allow for clear visualization of the main distribution of sites.

**Data S3. (separate file)**

Density distribution of SNP alleles, from nQuire denoised bi-allelic sites, supported by a given proportion of aligned reads for the 132 *M. capitata* RNA-Seq libraries that were analyzed in our study. For each plot, the x-axis is the proportion of reads that support a given SNP allele out of all reads that align to that site, and the y-axis is the frequency of SNP alleles with a given proportion of supporting reads. The name and putative ploidy of each sample is presented above its distribution; the number of denoised bi-allelic sites analyzed in each sample is presented below its distribution. The x-axis of these plots was limited to between 0.1 and 0.9 to allow for clear visualization of the main distribution of sites.

**Data S4. (separate file)**

Density distribution of SNP alleles, from nQuire denoised bi-allelic sites, supported by a given proportion of aligned reads for the 27 *M. capitata* RNA-Seq libraries (not from Kāne'ohe Bay) that were downloaded from SRA. For each plot, the x-axis is the proportion of reads that support a given SNP allele out of all reads that align to that site, and the y-axis is the frequency of SNP alleles with a given proportion of supporting reads. The name and putative ploidy of each sample is presented above its distribution; the number of denoised bi-allelic sites analyzed in each sample is presented below its distribution. The x-axis of these plots was limited to between 0.1 and 0.9 to allow for clear visualization of the main distribution of sites.
